# Buried in two places: Lineages from elite Maya tombs also found in distant caves

**DOI:** 10.64898/2026.06.07.724338

**Authors:** Esther S. Brielle, Elizabeth Dorgay, Douglas J. Kennett, Jose Mes, Emily Moes, Nadia C. Neff, Anna C. Novotny, Esteban Rangel, Erin E. Ray, Mark Robinson, Amy E. Thompson, Monica Warner, Ali Akbari, Kim Callan, Ella Caughran, Romain Fournier, Trudi Frost, Lora Iliev, Aisling Kearns, Jack Kellogg, Ann Marie Lawson, Iosif Lazaridis, Matthew Mah, Nihal Manjila, Mariam Nawaz, Iñigo Olalde, Jonas Oppenheimer, Iris Patterson, Lijun Qiu, Kendra Sirak, Gregory Soos, J. Noah Workman, Swapan Mallick, Nadin Rohland, David Reich, Keith M. Prufer

## Abstract

Classic Maya societies (250-900 CE) carved out urban centers in the rainforest reaching population heights and political complexity previously unknown in the Mesoamerican tropics. Kinship was a central feature of the social fabric, and rulership was legitimized through claims of direct descent from mythical ancestors. Mortuary practices kept the dead close to the living, and ancestor veneration sometimes produced complex deposits of disarticulated remains that are difficult to identify and whose biological relationships to one another cannot be understood without genetic data. We screened 487 human tooth and bone samples and successfully generated genome-wide data for 430 of them. Of these, 341 samples representing 107 distinct individuals were recovered from both elite and non-elite tombs in a Classic period kingdom (250-900) CE in the rugged Maya Mountains of Belize. Remarkably, 24 of these individuals have skeletal elements in both an elite tomb and in a ritual tooth cache in a distant cave located 26.5km away on the other side of the Maya Mountains. These results indicate that elite lineages created ancestors from their deceased relatives in geographically expansive ways and highlights the importance of caves in the belief system of Classic Maya elites.

## Introduction

Classic period (250-900 CE) Maya-speaking communities lived in hundreds of cities spread across southern Mexico, Belize, Guatemala, and northern Honduras^1^. They spoke distinctive Maya languages, but had common political, economic, and religious institutions. These Maya speaking communities had no documented open-air cemeteries. Instead, the deceased were commonly placed beneath the floors of houses, a practice that extended to all political and economic classes. Because social status was tied to lineal descent, burying relatives beneath the house helped keep ancestors physically present and reinforced a household’s claims to lineage and status. For political and economic elites, legitimacy was derived from their assertions of direct descent from powerful ancestors who had, in death, crossed back into the mythical underworld and become deities^2^.

The archaeological evidence shows that the living often kept the dead close^3^, allowing people to interact with their ancestors^4^. Mortuary rituals, therefore, could be both complex and elaborate^5^. Tombs included primary interments of one or multiple individuals^6^ and could also include secondary deposits of disarticulated skeletal elements^7^ or sacrificial victims^3^. Engaging with the ancestors was a recognized practice, and it often involved manipulating their skeletal remains and, at times, moving some elements to new locations^8^. Sealed burial contexts were sometimes reentered and skeletal elements were removed^9^. To access the dead, some tombs had dedicated entryways or formal stairways^10^. Resulting assemblages may represent multiple mortuary events with numerous individuals that accumulated over generations^11^.

In the Classic Maya worldview, tombs could be conceived as human-constructed caves^12^. Caves were considered access points to communicate with ancestors^2,13^, and thus had deep social meaning for Classic Maya speaking communities. But people did not live in caves, and they were less common places for burials than residential tombs. When human remains are found in caves, they often appear as isolated or a group of disarticulated skeletal remains rather than complete burials^14,15^. Archaeologists have typically interpreted articulated cave burials as either reverential interments or sacrificial deaths. Interpretation is largely dependent on context, evidence of peri- or post-mortem treatment, and age, with children frequently considered sacrificial^16^. In contrast, isolated skeletal elements that appear to be intentionally placed in caves are generally thought to have been moved from primary locations and redeposited as offerings related to ancestor veneration^17,18^.

Teeth are sometimes the most frequent elements in secondary deposits. Tooth caches are documented as offerings in architecture at Maya cities and are usually intentional deposits of small numbers of teeth accompanied by ceramics and other artifacts^10,19^. Occasionally these caches included hundreds of teeth^20^. More generally, teeth are excellent archives of genetic material and can be used to accurately estimate the number of individuals in an archaeological deposit.

Without genetic data, it is challenging to discern how disarticulated skeletal elements from archaeological contexts biologically relate to each other and greater Maya society. It is also difficult to determine relationships between individuals found in primary and secondary deposits^5^ and assess the role of ancestry and kinship in mortuary practices and the cultural meanings associated with caches of teeth and other disarticulated remains. Ancient DNA offers a powerful means of reconstructing and understanding the biological and cultural dimensions of Maya mortuary practice.

We successfully generated genome-wide data for 341 human skeletal and tooth samples (Methods), representing at least 107 individuals, recovered from elite and non-elite tombs in an ancient kingdom located in a valley high in the rugged Maya Mountains of southern Belize. We coanalyzed the data from tombs with additional genome-wide data for 89 skeletal elements, representing at least 36 individuals, recovered from a nearby rockshelter and two distant caves. With these, we reconstructed three lineages spanning four, six, and eight generations from otherwise disarticulated and comingled remains. Skeletal remains from 24 individuals were simultaneously buried in two places, namely teeth interred in an elite residential tomb beneath their house and additional teeth from the same 24 individuals placed in a remote funerary cave 26.5 km away from this residence (Figure 1)^21^. These teeth were placed around and above the pelvis of a primary burial of an adult female in the cave who was also accompanied by high status grave goods. This primary individual was a 4^th^ degree ancestor to one of the lineages as well as other individuals interred in the elite tombs.

**Figure 1:**
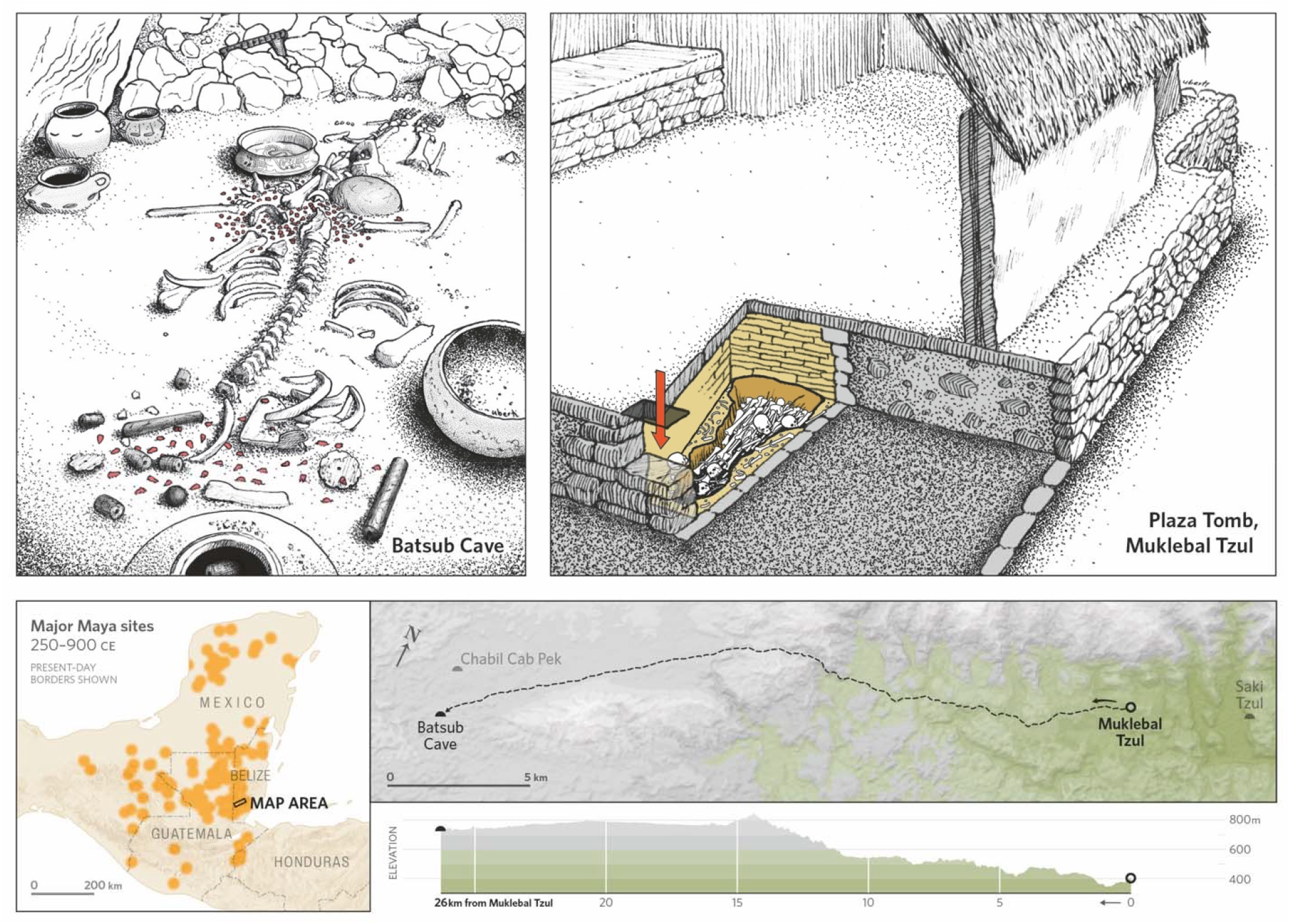
Upper panels, left to right: Artist rendition of the Bats’ub cave (BS) primary burial and assemblage and of the Muklebal Tzul (MKB) Plaza Tomb (PT) (SI Discussion 1). Lower panels, left to right: Map of the Maya lowlands with approximate locations for Maya population centers and a rectangle indicating the study area. Map showing the least cost path between the MKB and BS sites with elevation profile (SI Discussion 2). Chabil Cab Pek cave (CCP) and Saki Tzul (ST) are also shown in the least cost path map.

### Site descriptions

**Muklebal Tzul (MKB)** is a Classic period Maya population center that lies in an interior valley of the rugged Maya Mountains, situated in the upper watershed of the Bladen Branch of the Monkey River (SI Discussion 1 and Figures. S1.1 and S1.2) ^22^. The site core of MKB is a large stelae plaza connected to elite residential compounds where the two large tombs, Tomb 1 (T1) and Plaza Tomb (PT), were excavated in 1995 and 1996 (Figures S1.2-7)^22^. The non-elite settlements at MKB are primarily located to the west and south of the site core. Most settlements include at least one tomb, with several containing multiple tombs (Figure S1.5). In addition to the two large elite tombs, we include individuals from three settlement group tombs West 5 Structure 36 (W5 S36), West 1 Structure 4 (W1 S4), and West 1 Structure 6 (W1 S6) (Figures S1.8-10).

**Bats’ub Cave (BS)** is a nondescript cavern on the side of a small hill (Figures 1 and S1.11), contemporaneous with MKB. The primary burial in this cave was the skeleton of an adult female placed in a 20–25 cm deep pit cut through the cave’s gravelly floor. The head of this individual was removed, and in its place was the bottom half of an undecorated vessel containing a single jade bead. Fragments of a poorly preserved cranium, possibly her own head, and fragmented mandible with no teeth were found in the area of the pelvis. A bowl containing five cacao seeds had also been placed inverted above the pelvis^23^. At least 144 teeth were recovered in a cluster from this same area around the pelvis while another small cluster of teeth was found near the thoracic vertebra. Other teeth were found scattered throughout the burial context for a total of 226 teeth. There were high status grave goods accompanying the primary burial, indicating elite status and evoking an underworld scene from the Maya creation story^24^.

**Chabil Cab Pek (CCP)** is a cave located 1.7 km north of BS, and is contemporaneous with the early occupation period of MKB. Two older female adult burials were recovered from CCP. Both individuals were incomplete and were placed next to a small stone altar along with a painted Early Classic basal flanged bowl (Figure S1.14).

**Saki Tzul (ST)** is a large rockshelter on a hillside overlooking Ek Xux, a neighboring valley site lying 3.7 km east of MKB within the Maya Mountains (Figure 1 and S1.1). ST is one of two neighboring rockshelters in the upper Bladen River that are the oldest mortuary sites known in the Maya-speaking world with burials dating back to the Late Pleistocene^25,26^. Burials in the uppermost levels of the excavations date to the Classic period (Figure S1.13)^27^.

## Results

We sampled 367 ancient skeletal elements from five tombs in MKB. Two tombs (PT and T1) are in elite residences and date to the Classic period, while the other three tombs (W5 S36, W1 S4, and W1 S6) are in non-elite residences of MKB and date the mid to late Classic period (550-775 CE). We also sampled 5 skeletal elements from a nearby rockshelter ST, and 115 from two distant caves, BS and CCP. All work at these sites and subsequent analyses was done with permission from local authorities and consultation with local communities (Methods).

We used in-solution enrichment for more than 1.4 million single nucleotide polymorphisms (SNPs) to generate genome-wide ancient DNA from 487 skeletal elements, of which 430 gave working genome-wide data passing initial quality control (Table 1). To determine relatedness (SI Discussion 3), we used pairwise allelic mismatch rates (PMR, READv2^28^), pairwise number and total length of identity by descent segments (IBD, ancIBD^29^), and maximum likelihood pairwise linkage disequilibrium rates (reLD).

**Table 1:** There were 367 skeletal elements sampled from the tombs at MKB, 115 from the distant caves, BS and CCP, and 5 from the nearby rockshelter ST. Approximate distinct individuals is determined genetically. Minimum number of individuals (MNI) is determined osteologically with details in SI Discussion 1. Detailed osteological data are found in Online Tables 6-11. In the sub-table showing replicate individuals, context refers to the specific mortuary location or combinations of locations where skeletal elements were found. Total number of individuals will be less than the number of individuals found in each context if some of the individuals are replicates.

| SITE DESCRIPTION | SITE / CONTEXT | NO. SKELETAL ELEMENTS SAMPLED | NO. SKELETAL ELEMENTS PASSING INITIAL QUALITY CONTROL | APPROXIMATE DISTINCT INDIVIDUALS | APPROXIMATE DISTINCT INDIVIDUALS (>50,000 SNPs) | BONE MNI | TOOTH MNI |
| --- | --- | --- | --- | --- | --- | --- | --- |
| Elite tombs at Muklebal Tzul | MKB T1 | 3 | 3 | 3 | 3 | 3 | 8 |
|  | MKB PT | 336 | 314 | 144 | 88 | 41 | 28 |
| Non-elite / settlement tombs at Muklebal Tzul | MKB W5 S36 | 24 | 21 | 19 | 17 | 12 | 21 |
|  | MKB W1 S4 | 2 | 2 | 1 | 1 | 2 | 2 |
|  | MKB W1 S6 | 2 | 1 | 1 | 0 | 0 | 2 |
| Total at MKB |  | 367 | 341 | 166* | 107* |  |  |
| Distant caves | CCP | 2 | 2 | 2 | 2 | 2 | 1 |
|  | BS | 113 | 82 | 39 | 30 | 4 | 15 |
| Total in caves |  | 115 | 84 | 41 | 32 |  |  |
| Rockshelter | ST | 5 | 5 | 4 | 4 | - | - |
| Total |  | 487 | 430 | 193* | 118* |  |  |

\* all totals are calculated after removing replicate instances of the same individual

| CONTEXT 1 | CONTEXT 2 | NO. OF CONTEXT 1 IND. | NO. OF CONTEXT 2 IND. | TOTAL NO. OF IND. | NUMBER OF CROSS-SITE REPLICATES | CONTEXT 1 IND. NOT IN CONTEXT 2 | CONTEXT 2 IND. NOT IN CONTEXT 1 |
| --- | --- | --- | --- | --- | --- | --- | --- |
| MKB PT | MKB T1 | 88 | 3 | 89 | 2 | 86 | 1 |
| MKB PT | BS | 88 | 30 | 94 | 24 | 64 | 6 |
| MKB PT & T1 | BS | 89 | 30 | 95 | 24 | 65 | 6 |
| MKB PT | CCP | 88 | 2 | 89 | 1 | 87 | 1 |
| MKB PT & T1 | BS & CCP | 89 | 32 | 96 | 25 | 64 | 7 |
| MKB all tombs | BS, CCP, ST | 107 | 36 | 118 | 25 | 82 | 11 |

The majority of samples with working data came from MKB PT, with 314 out of 336 teeth producing genetic data (see Table 1). While we expected that any one individual might have as many as 32 teeth in the tomb, it was surprising that we still identified a minimum of 88 distinct individuals at PT based on genetic data. This represents the largest number of unique individuals in a single tomb documented in Mesoamerica.

### Replicate burials

Even more surprising was how frequently elements from the same individual were recovered from multiple, geographically distant sites. We had no reason to expect any connection between BS and MKB given the considerable distance between them and no prior archaeological evidence of ties between the sites. Strikingly, many of the genetic links between these sites did not simply reflect shared lineage but revealed that different skeletal elements of the same individuals were found in both the elite family tombs at MKB and in the underworld scene at BS (SI Discussion 1). There are 89 distinct individuals found at PT (88) and T1 (3), with 2 individuals having teeth in both elite tombs. There are 32 distinct individuals found at BS (30) and CCP (2), 25 of whom were cross-site replicates, namely individuals who have teeth interred in both an elite tomb and a distant cave. Of these cross-site replicates, 24 are from BS and 1 is from CCP (Table 1).

Of the 30 individuals recovered from BS, 24 are also found in PT, far more overlap than expected by chance (one-sided Fisher exact test *p* = 4.9E-7 for [24,6;24,64]; simulation-based right-tailed *p* = 0.04 under a null model that randomly assigns “BS” and “PT” element labels; SI 8). Those same 24 individuals, however, are only a minority of the much larger PT pool of at least 88 individuals, far less overlap than expected by chance (simulation-based left-tailed *p* = 4 · 10^−4^; SI 8). These results support an asymmetric relationship in which BS elements come selectively from a subset of PT individuals, rather than two parallel burial populations with random overlap. This is consistent with PT as the main lineage center and BS as a selective component of MKB elite mortuary practice, rather than a separate cemetery or community tomb in its own right. The subset of PT elites that had replicates in BS is not distributed uniformly across time. Before about 520 CE, individuals with cross-site replicates are rare (1 of 7), whereas after about 520 CE, being buried in both places is much more common (21 of 35), a significant temporal shift toward higher overlap (Fisher exact *p* = 0.0408; SI 8).

In total, we identified 118 distinct individuals across the sites (Table 1 and Online Table 2). This set includes individuals for whom at least one skeletal element yielded ≥50,000 SNPs or more. The additional 75 individuals at most, represented by 76 skeletal elements from BS and the MKB tombs, did not produce sufficient data (< 50,000 SNPs) to resolve identities or relationships although we report them (SI Table 1, Online Table 2).

### Lineages

We pooled the data from all replicate samples for each individual to increase resolution. Among the 118 unique individuals with high quality data, we identify close relatives for 59, with 42 of them resolved into three distinct pedigrees (Figure 2). Elite lineages 1 and 2 span 6 and 8 generations, respectively, with most individuals found primarily in the MKB PT and T1 tombs along with their replicate interments at BS. Of the 31 PT individuals that make up or are associated with these two lineages, 14 (45%) are interred at both PT and BS, and 2 (13%) are interred at T1, PT, and BS. These two lineages are connected to each other by distant relatives of approximately 4^th^ degree. From the non-elite settlement tomb W5 S36, we also document the separate 4 generation deep Lineage 3 without any detected relatedness to the PT or T1 tombs or those interred at the caves and rockshelter. There are 8 additional individuals whose positions cannot be resolved within a lineage, but who were interred at both PT and a cave. Of these, 7 were interred at PT and BS and 1 was interred at PT and CCP. One of the CCP females dates to 244-362 calCE (NOSAMS-204278, 1751±18 BP) and shows that the connection between MKB and distant caves arose early in the occupation of MKB. The primary burial at BS and the second individual at CCP date to 250-402 calCE (NOSAMS-203735, 1733±23 BP) and 255-402 calCE (UCIAMS-280706, 1725±15 BP) respectively, which are nearly identical dates that also align with the earliest known occupation at MKB. Among the remaining 51 individuals with sufficient coverage, 6 are from BS (one of these is the primary female burial genetically identified by a long bone and a tooth), 1 is from CCP, 32 are from PT, 1 is from T1, 10 are from W5 S36 (one of these is represented by 2 teeth), and 1 is from W1. There are many undiscovered or unsampled sites in the Maya Mountains so we do not exclude the possibility of additional replicate interments elsewhere.

**Figure 2:**
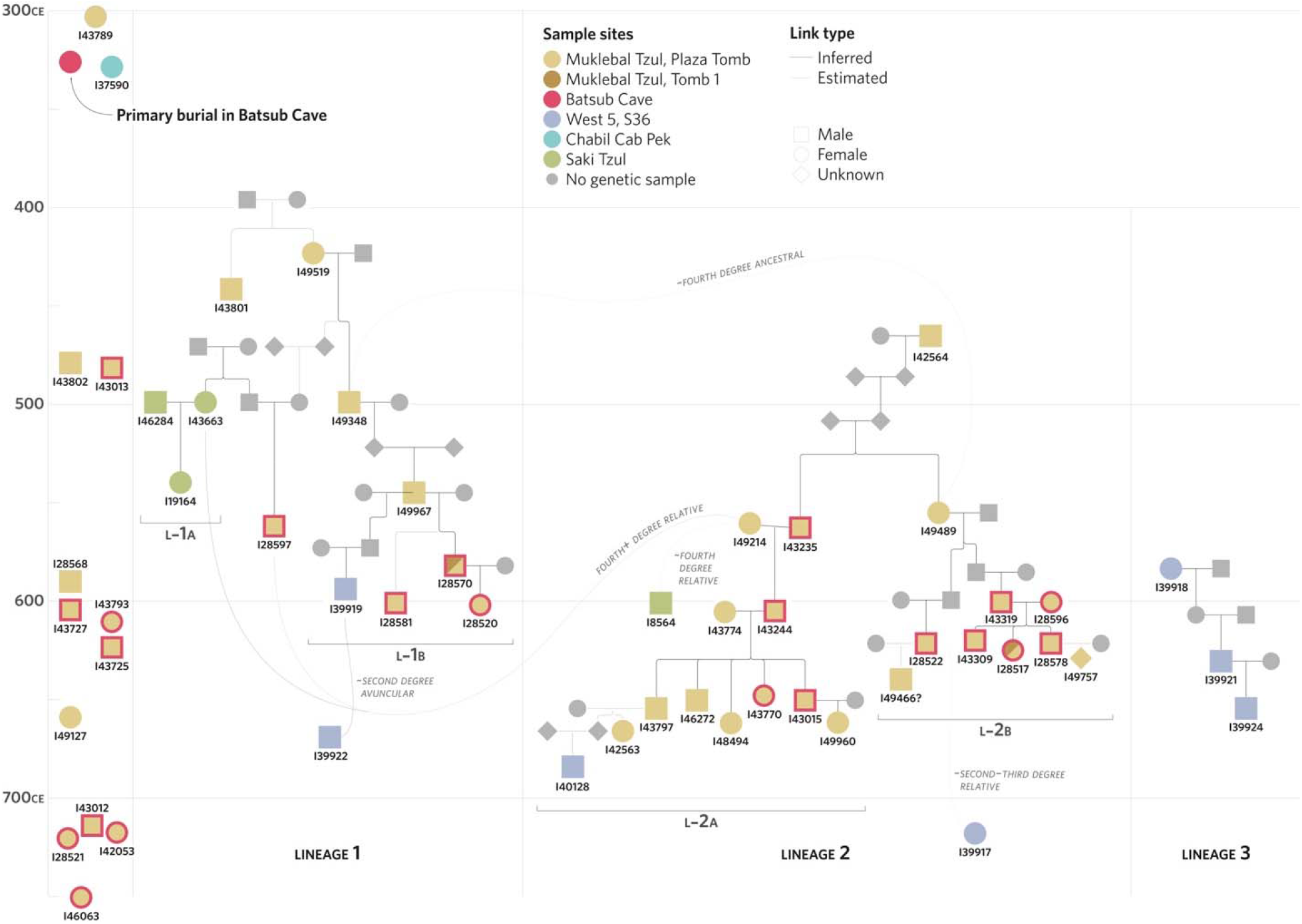
Three reconstructed lineages from ancient DNA samples recovered from MKB, BS, and ST. Individuals interred at MKB PT are tan, at MKB W5 S36 are blue, and at ST are green. Cross-site interments are indicated by an additional red outline for BS, a teal outline for CCP, and a split light and dark background for interments at both MKB T1 and PT. All individuals interred at BS in these lineages are also interred at MKB PT. The two individuals interred at MKB T1 are also interred at MKB PT and BS. There are two individuals interred at CCP, with one of them (I43789) also interred in MKB PT. Individuals are positioned at their approximate modeled dates, according to Figure S4.3 and Table S4.4. Gray dashed lines depict relationships that are not fully resolved due to insufficient coverage or distant pairwise relationships.

Lineage 1 (L1) is derived from the elite tombs at MKB, as well as from BS and ST. L1 contains two sub-lineages (L1-a and L1-b) connected by individual I28597, interred in both PT and BS. We model the duration of lineage L1 to 255±62 years, spanning six generations from 389±43 CE to 651±34 CE (see Extended Data Table 1). Individuals from the oldest four generations of L1-b were only interred in PT, while individuals from the final two generations were also interred in BS, with one of them interred in T1 as well. We also identified an unexpected link involving a male, I49967, from L1-b, who was interred in PT. This male has descendants interred in both PT and W5 S36, albeit from two different women. From one maternal line, we identified two sons and a granddaughter interred in PT. From a second maternal line, we identified a single grandson, I39919, interred in W5 S36. If I39919 was born to a woman who was considered part of the L1-b lineage, we would expect I39919 to be interred in PT. Instead, his burial in W5 S36 suggests a connection outside the core PT household. We therefore do not include I39919 as a member of L1-b. The L1-a sub-lineage is a family of both parents and their daughter buried at ST. Individual I28597, who connects the L1-a and L1-b sub-lineages, is related avuncularly to the mother in L1-a and is the great-grandson of the oldest generation woman in L1-b. The mitochondrial DNA haplogroups differ between connecting individual I28597 and family L1-a, meaning that I28597 is related to L1-a through at least one male instead of a shared maternal line. Since I43663 is a female, her sibling, who is also a parent of I28597, must be a male. This suggests that either I28597 or his father joined into this MKB L1 family, while their other close relatives were buried in ST, located in the neighboring valley, where the separate Ek Xux polity was located (Figure S1.1). I28597’s mother is likely the granddaughter of I49519, the oldest woman in the L2 lineage who was interred in PT, suggesting that I28597’s mother may herself have been part of a long-standing MKB family.

Lineage 2 (L2) is represented in both elite tombs and BS, and spans eight generations from 361±89 years from 396±51 CE to 775±58 CE (Table S4.4). Lineage 2 comprises two related five-generation sub-lineages, designated as L2-a and L2-b. These sub-lineages are related through their oldest generation, a brother in L2-a and his sister in L2-b. The great grandparent of these siblings was also found in PT, represented by 12 teeth and dating to 444-600 calCE (NOSAMS-204281, 1527±18 BP), but the following two generations were not identified. All members of L2-a and L2-b were interred in PT. Within Lineage 2, cross-site replicate prevalence differs between the two sub-lineages, with L2-b showing a higher rate than L2-a. In L2-a, 4 of 12 individuals are cross-site replicates, whereas in L2-b, 6 of 7 individuals are cross-site replicates, with one of them having an additional element interred at T1 as well. This difference is supported by Fisher exact tests and a simulation-based power analysis and is consistent with treating L2-a and L2-b as distinct sub-lineages (SI 5).

Lineage 3 (L3), associated with the non-elite tomb in the MKB settlement group W5 S36, spans four generations and is represented by 3 individuals from 19 sampled teeth. The duration of lineage L3 spans 195±76 years, from 516±47 CE to 726±46 CE. L3 thus broadly overlaps with lineages L1 and L2, although it begins later.

In contrast to the elite lineages, none of the individuals in the non-elite lineage L3 have replicate interments. While L2-b has a large proportion of cross-site replicates and differs significantly from L3, most lineage-level comparisons are underpowered (SI 5). We therefore focus on significant differences at the level of tombs and caves, which have increased sample sizes providing higher power. The proportion of cross-site replicates differs across all mortuary contexts (*χ*^2^ *p* = 2.6×10 □ □, SI 5). This indicates that cross-site replication is concentrated in a subset of contexts rather than being evenly distributed across the mortuary landscape. Cross-site replicates are present in the elite tombs PT and T1 but absent from the non-elite tombs W1 and W5. Among the 89 individuals from the elite tombs PT and T1, 25 are cross-site replicates, while among the 18 individuals in non-elite tomb contexts W5 and W1, there are 0 cross-site replicates (Fisher exact *p* = 0.006). The same pattern holds true when down sampling PT and T1 skeletal elements to match the same number of elements sampled from W5 and W1 (SI 5), and it is also seen when comparing PT alone to W5 alone (25 of 88 versus 0 of 17, Fisher exact *p* = 0.01). Overall, the statistics show that cross-site replication is strongly concentrated in elite tombs and essentially absent from the non-elite tombs we sampled (SI 5 for Fisher tests and SI 7 for *χ*^2^ tests).

There is also a significant association between individuals being interred in a distant cave and having a greater number of replicates at PT (*χ*^2^ test *p* = 3.98E-05, SI 7). In other words, the individuals represented at BS tend to be the same individuals represented by multiple elements within PT. This indicates non-random selection for cave deposition and may partly reflect chronology, with later generations in the elite lineages more often represented in both contexts. There is no association between the presence of cross-site replicates and molecular sex (*χ*^2^ test *p* = 0.1, SI 7). We also find that there is no significant association between males and females in any elite lineage (L1-b, L2-a, L2-b) and if they had any cross-site replicates in a distant cave (Fisher p = 0.4, SI 6).

Internal transmitters, individuals with both an ancestor and a descendant within the reconstructed elite lineages from MKB, are disproportionately male (Fisher exact test *p* = 0.01, SI 6). This is consistent with a pattern of women more often entering the lineage through marriage and men more often being born into the lineage and having their male children remain in the reconstructed pedigree and thus the lineal household. Relatedly, family membership is strongly associated with the mitochondrial haplogroup (*χ*^2^ test *p* = 1.71E-08), consistent with different maternal lines feeding into different sub-lineages. These tests support patrilineal descent and patrilocal residence, with outside females marrying into the lineage, and bringing different mtDNA into their respective family branches.

### Primary ancestor

A single long bone (tibia) from the primary BS burial was directly dated to 250-402 calCE (NOSAMS-203735, 1733±23 BP; Extended Data Table 1) and is a genetic match to a tooth recovered from BS from the scatter of teeth near the primary individual’s neck. We did not identify any first to third degree relatives of the primary individual at any site, which is not unexpected given that she predates the earliest reconstructed lineage members by 104 years, with a median date of 331 calCE for the primary BS burial versus 435 modeled years CE (modCE) for I49519, the earliest individual in L1. Instead, the primary BS individual, as well as the CCP burials, show distant relatedness (≥4th-degree) to multiple individuals at PT, BS, and ST (Online Table 2). Given the distant relatedness of the primary BS burial to PT and BS individuals, it is parsimonious to interpret her as an ancestor or ancestor-like relative whose real or fictive descendants venerated her with offerings of their teeth.

There are two possible scenarios to explain the use of BS as a mortuary site; first, a single funerary event after the date of death of, or with the extraction of a tooth from, the latest individual deposited in the cave; or second, a series of funerary events over time. Direct dating of teeth and organic samples from BS help inform the chronology of the site (Extended Data Table 1). At one end of the date range, two teeth, without associated genetic data, date to 45 calBCE-125 calCE (UCIAMS-14914, 1975±35 BP) and 26-219 calCE (UCIAMS-14913, 1910±35 BP), significantly predating the primary burial and making them the earliest known interments at BS and in the study. These teeth were not recovered from the main tooth caches near the pelvis or neck. At the other end of the date range, the latest dated tooth from BS cave has a lower bound of 666 calCE, significantly postdating the primary burial. Under a single-event scenario, the remains of the primary female, the tooth cache, and the grave goods would have been deposited together after 666 calCE. However, at least two pieces of evidence do not support this single-event scenario. First, radiocarbon dates of charcoal and wood torch fragments from BS predate several teeth in the main tooth caches, indicating use of the cave before the death of many of the people whose teeth were deposited in BS. Second, the primary burial is unlikely to be a secondary deposit because the parts of the skeleton in contact with the wet clay surface of the cave, including the spinal column and ribs, were in correct anatomical articulation when excavated. The most likely scenario is thus a series of funerary events, with the primary individual buried shortly after death and then later deposits of the teeth of real or fictive descendants were added during one or more subsequent visits to the cave over the following centuries.

## Discussion

Genetic identification of replicate interments provides direct evidence that the remains of a single individual were distributed across multiple mortuary sites. Most of these replicate interments belonged to deep elite lineages with patrilineal descent and patrilocal residence. Our data suggest an intentional strategy to anchor these lineages simultaneously in elite houses and in a cave transformed into an elaborate underworld funerary setting.

There is a considerable body of archaeological research that has documented the central role of caves and mountains as powerful religious landmarks in ancient Mesoamerica^30^. Caves long functioned as thresholds, marking the place of emergence of humans out of the mythic time and space where they were first constituted, and thus became places where the elite communicated with those ancestral forces^6^. By 1000 BCE, rulers linked their legitimacy to a divine ability to mediate with deities responsible for health, fertility, and rainfall who inhabited a vast mythological watery underworld inside mountains, which were accessible through caves^31^. Classic period rulers publicly proclaimed that they derived their supernatural authority from their ancestors^4^, passing it down to their descendants, and forming powerful lineages. They prolifically recorded their relationships with underworld-dwelling ancestral deities in hieroglyphic texts and as imagery on finely crafted ceramics, codices, and stone sculpture. Rulers frequently depicted themselves in cave-like settings, conjuring ancestors and presenting offerings of their own blood or, less frequently, the bodies of others to appease powerful deities. These same rulers constructed tombs and temples as physical manifestations of caves and mountains. In some well-preserved tombs carved into the bedrock beneath elite buildings, hieroglyphic writing on the walls describes these tombs as the interiors of mountain-caves ^32^.

While caves were important but infrequent funerary locations, the substructural spaces beneath houses, and specifically tombs, were also considered to be places where ancestors dwelled.

Archaeologists have long inferred that people living in the Maya-speaking region buried their dead below the floors of their houses to maintain relationships with ancestors to legitimize the rights of the living to specific local resources^3^. Elites further claimed descent from ancestors who dwelled in the underworld, thus additionally giving them exclusive rights to political and religious authority^4^.

Bats’ub Cave and its complex funerary assemblage are appropriate for a royal individual in an underworld setting (SI Discussion 1, Figure S1.12). It epitomizes the way elites connected their ancestors to caves and to rituals of death. The most elaborate vessel, a black and red on orange basal-flange bowl (Figure S1.12), characteristic of the Early Classic^33^ depicts a scene from the Maya underworld. The exterior shows two individuals—or possibly the same individual repeated on both sides of the vessel—lying face up, with protruding tongues and eyes as narrow slits, details suggestive of a death pose. In this scene, the head rests atop the abdominal region of the body. The feet abut what is likely a bloodletter bundle^21^, elsewhere described as containing stingray spines (bloodletting implements)^3^. The interior surface of the vessel is decorated with a mythical feathered hummingbird-serpent creature emerging from an object that could be a torch handle or a stingray spine. Death poses, skull-removal, bundles, torches, stingray spines, and feathered serpents are all themes associated with an underworld setting. CCP also contained a highly eroded Early Classic basal flange vessel depicting a ballplayer from a mythical underworld scene. Finally, a small stool carved from a single piece of rosewood and placed near the BS burial is the first of its kind found in a mortuary context, yet consistent with an underworld setting and as the possession of an elite^34^. In the creation myth, Popol Vuh, stools are described as property of underworld deities^23,35^.

Given that the elite and non-elite lineages in this study probably lived in and around MKB and were found there in residential tombs, the choice of BS (and CCP) was specific. There are many caves closer to MKB and the physical journey to BS would have required a multi-day trek across 26.5 km of extremely rugged terrain with jagged limestone hills and deep sinkholes. The deliberate transfer of selected skeletal elements of lineage members to this distant cave suggests that BS held particular religious and political connotations for the MKB elites, especially as they related to their ancestors.

Our data suggest that MKB elites did not confine mortuary practices to any single tomb or site, but they also interred their ancestors in caves. MKB non-elites, however, were more likely to manifest their relationships with ancestors within their household. This could reflect different social ideologies surrounding kinship or mortuary practices in regard to how ancestors are venerated between elites and non-elites. Even with these complex arrangements of skeletal elements interred between tombs and caves we have not accounted for the bodies of most of the individuals whose teeth are interred at MKB or BS.

We present the first empirical genetic and mortuary evidence for kinship organization, and for how descent was reckoned in Classic period elite families in the Maya-speaking lowlands. The composition of the lineages interred at PT and T1 indicates that descent in elite families at MKB was patrilineal. Males were interred significantly more often with their fathers and male ancestors along with their sons and male descendants, while significantly more non-descendent women (women who were not born into the lineage) are entering into these lineages through marriage. These interments are in household tombs, further suggesting patrilocal residence, namely that related males lived in the same residential compounds with non-descendent women who were marrying into these families. Females in the PT and T1 lineages have skeletal elements that are represented as offerings along with those of males at BS, suggesting that elements from both males and females played an important role in the centuries-long series of ancestor veneration rituals provided to the primary female ancestor, buried at BS.

This study reinforces the concept that where individuals were buried reflected their status in life, their role as members of families and lineages, and, in death, their status as ancestors to those still living^13^. The BS funerary assemblage centers on a primary female 4^th^ degree ancestor to the PT and T1 lineages. Teeth from at least 39 unique individuals were placed around her pelvis, suggesting that the descendants petitioned their ancestor over subsequent centuries. Distributing the remains of a person in both tombs and caves also suggests that ancestor veneration may have been more expansive than a single residential tomb. Viewed as a portal to the underworld, BS may have offered MKB elites a place to position their ancestors, and by extension themselves, as mediators with the supernatural forces that animate the world.

## Methods

### Inclusion and ethics

All ancient skeletons for this study were excavated and analyzed under permits issued by the Belize Institute of Archaeology (IA) and the Belize Forest Department. Skeletons of ancient individuals were exported under permits issued by the IA to K.M.P. in accordance with the laws of Belize and permission granted to conduct molecular DNA analyses. Research was conducted in close collaboration with the Ya’axché Conservation Trust, an internationally recognized Belizean NGO that is the co-manager of the Bladen Nature Reserve (BNR) with the Government of Belize. Ya’axché is locally managed and largely staffed by members of descendent Maya communities. As part of this collaboration, our research proposals are annually reviewed by the Ya’axché administrative and scientific staff. In 2016, 2018, 2020, 2022, and 2024 in coordination with Ya’axché, we invited indigenous leaders and community members from villages proximate to the BNR to consult on this research. We provided advance invitations and arranged transportation for up to 50 people from five villages to attend the public consultation. At each consultation we presented results of field and laboratory studies to the communities, and additional time was devoted to answering questions and clarifying the data and interpretations^1^. Local participants were asked to provide feedback and ideas for future collaborative efforts and to be presented at future consultations. Community members requested future public consultations to update them on additional research results, as well as copies of all study results in English with translations into Mopan and Q’eqchi’ languages.

### Least Cost Path Analysis

We computed the least cost path (LCP, SI Discussion 2)^2^ between MKB and BS using a 12.5m Digital Elevation Model (DEM). The LCP model indicates that the 26.5 km journey would have taken 6 to 7 hours. While the least cost path does account for the rough and steep terrain, the time calculation along this path does not account for how this terrain would wear a person down over time, especially if they were carrying a load. Given the mountainous terrain, the journey would take more than one day.

### Ancient DNA data generation

Ancient DNA skeletal sampling and library preparation was performed using established ancient DNA protocols in dedicated ancient DNA clean rooms. While the cochlea from petrous bones normally contains the highest endogenous DNA yield, when that was not available, teeth were sampled. Specifically, from MKB, there was a significantly greater minimum number of individuals (MNI) determined from teeth than the MNI determined from bones. We sampled 350 teeth from MKB along with 10 petrous, 2 temporal, 1 parietal, and 4 long bone samples. From BS, aside from the primary individual, which we represented with a tibia, the only skeletal elements were tooth caches, from which we sampled 113 teeth. The petrous bones and the rest of the skull of the primary individual from BS were too deteriorated to yield DNA. The four samples obtained from ST included 2 petrous bones and 2 carpals. We sampled two petrous bones from skeletal remains at CCP.

Sampling to collect powder for DNA extractions in our cleanrooms comprises of two steps, first, the sample cleaning and decontamination, and second, and the sampling itself. Teeth and non-petrous bone fragments are UVC irradiated in a cross-linker for 5-10 minutes before debris and the outer layer of the spot we selected for sampling are removed with a sanding disk attached to a dentistry drill; this is to minimize exogenous DNA contamination and dirt entering the DNA extraction. Since we sample from the inside of the petrous bone (we either isolate the cochlea using a sandblaster ^3^ or drill into the cochlea to collect powder) we only superficially clean away the dirt from petrous bones with air from the sandblaster; this sometimes dislodges auditory ossicles^4^ that we then use directly without damaging the bone. Teeth are drilled into with dentistry drill bits to obtain ∼40 mg of fine sample powder by targeting the outer layer of the teeth (cementum^5^), shown to contain the highest endogenous DNA content in teeth, as well as dentin. For non-petrous bone fragments, we select a compact part to drill into to collect ∼40mg of powder. For petrous bones, when possible, the cochlea portion is isolated by removing the surrounding less dense bone using a sandblaster and then ground with a mixer mill to obtain fine powder; ∼40 mg of this bone powder^3^ is used for DNA extraction.

DNA is extracted from ∼40mg of the bone powder using protocols optimized for short, damaged molecules^6,7^ using automation with magnetic silica beads. The DNA extract is then incubated with uracil-DNA-glycosylase (USER from NEB), an enzyme mix that cleaves deaminated cytosines (uracil bases) and cuts abasic double strands, but cleaves inefficiently at terminal uracils, which allows for authentication of the presence of ancient DNA segments. Barcoded single^8^ or double stranded^9^ libraries are prepared (ss.USER and ds.half in Online Table 1) for the samples.

For the majority of libraries, we used Twist(^™^□) Bioscience’s capture technology to enrich the DNA libraries for sequences containing target DNA segments. This capture reagent includes 1,434,155 million 80 base pair probes overlapping 1,352,535 targeted single nucleotide polymorphisms (SNPs) in the nuclear genome. The target SNPs were chosen to maximize information about population history, or to be relevant for Y haplogroup determination or associated with phenotypes^10^. A separate reagent of mitochondrial probes targeting the complete mitochondrial genome and helpful in mtDNA haplogroup determination was also added to the reagent. DNA segments that bind to these biotinylated probes are then captured on streptavidin beads, washed repeated times and subsequently sequenced from both directions (paired-end reads) on Illumina instruments.

Some of the libraries, particularly those created before the Twist Ancient DNA capture reagent was fully implemented (June 2021), were captured using the 1240K capture reagent, which targeted 1,200,343 SNPs on chromosomes 1-22 and X as well as 32,670 SNPs on chromosome Y^11^. These, too, were combined with mtDNA probes of the whole mt genome. Any DNA segments that hybridized to the probes of the biotinylated 1240K reagent were washed repeatedly and then sequenced with paired-end reads on Illumina instruments. To improve the rate of enrichment and therefore the number of targeted SNPs hit, most libraries were enriched in two consecutive rounds of capture; this was done for all libraries enriched with the 1240k protocol and for single-stranded libraries enriched with the Twist Ancient DNA kit (double-stranded libraries enriched with Twist only underwent a single round of capture). Several samples that were sequenced after capture with the 1240K reagent were also sequenced after capture with the Twist reagent, and in these cases, the sample’s DNA dataset includes a merge of the sequencing paired-end reads from both capture technologies.

Numerous DNA libraries were pooled for sequencing, and so the first bioinformatics step is to bin each sequencing read into individual libraries based on a library-specific barcode. The paired-end forward and reverse reads are merged into a single read if the length of the merged reads is at least 30 and if at least 14 out of 15 consecutive base pairs overlap for a base quality of at least 20 and at least 12 out of 15 consecutive base pairs overlap for a base quality below 20. For mismatched bases in the forward and reverse reads, the base with a greater base quality score is chosen. The barcodes and adapter sequences are trimmed for all reads. The reads are then aligned to the human reference genome GRCh37 (hg19)^12^ as well as the Reconstructed Sapiens Reference Sequence (RSRS)^13^ reference mitogenome using the `bwa aln` alignment command followed by the `bwa samse` alignment command of the Burrows-Wheeler Aligner (BWA) tool^14^ with parameters `-o 2 -n 0.01 -l 16500`. This 2-step process aligns reads to all possible mapping positions followed by randomly assigning the read to one of the mapping positions. The output BAM files with all aligned reads are then filtered to remove duplicates and to retain only reads that successfully aligned to a reference sequence.

For UDG partial (half) treatment DNA libraries, two base pairs were clipped from both the 5’ and 3’ ends of each read to remove nucleotides that typically have a high rate of deamination damage in ancient DNA. For UDG-minus DNA libraries, the adaptive pulldown script determines from the unclipped libraries what positions contain damage and thus what the optimal clipping would be, usually 5-10 base pairs.

From the BAM file for each library, ‘pseudo-haploid’ genotypes were created by selecting one allele for each SNP position. This allele was randomly selected from among the alleles of reads that overlap the SNP position and that have a base quality score of at least 20 and a mapping quality score of at least 10. This pseudo-haploid data was pulled down to three files, an ind file, snp file, and geno file. The *.ind file has three columns: individual or sample ID number, genetic sex, and population label. The snp file has six columns: SNP ID (often rsID), chromosome number (X chromosome is 23), genetic position of the SNP in Morgans, physical position of the

SNP in bases, reference allele, and variant allele. The geno file is a N_IND_ X N_SNP_ array with values of 0,1,2, and 9. Values of 0, 1, or 2 denote that at the respective SNP column and individual/sample row, the allele is respectively the homozygous reference allele, heterozygous for the reference and alternate alleles, or homozygous alternate allele. For pseudo-haploid data, values of 1 are not present and values of 0 and 2 refer to the single randomly selected SNP. Values of 9 indicate SNPs where there are no overlapping reads of sufficient quality and the allele is characterized as missing.

There are 2,142,271 SNPs for which we incorporate allelic information, including both target and off-target SNPs captured on the Twist capture reagent (which incorporates the 1240K capture reagent).^10^ We then removed 898,740 SNPs that were determined to bias co-analysis of libraries created from the different capture reagents and from shotgun data. This left a total of 1,243,531 SNPs on this compatibility SNP panel.^15^

We obtained DNA sequencing data passing quality control from 331 of the tooth samples from MKB along with 6 petrous bone samples and the 4 long bone samples. From BS, we obtained DNA of sufficient coverage from 82 teeth and 1 longbone. From CCP, we obtained sufficient coverage DNA from 2 petrous bone samples. From ST, we analyzed 5 skeletal elements; 3 petrous, 1 carpal, and 1 metatarsal, that provided sufficient coverage for analysis.

### Genetic sex and haplogroup determination

Genetic sex was determined by calculating the fraction of sex chromosome reads that aligned to the Y chromosome, namely 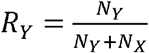, where *N*_y_ is the number of reads that aligned to the Y chromosome, and *N*_*X*_is the number of reads that aligned to the X chromosome.^16^ If *R*_*Y*_ > 0.32, then the individual was classified as male. If *R*_*Y*_ < 0.03, then the individual was classified as female. For the individuals where 0.03 < *R*_y_ < 0.32, the genetic sex was undetermined. After mtDNA alignment and clipping, variant calling is done using samtools mpileup with a minimum mapping quality of 30 and minimum base quality of 20, which is then piped into bcftools call -c -v --ploidy 1 and then output in vcf format. The mtDNA haplogroup was determined from this vcf file using Haplogrep 3.^17^

Y haplogroups were determined using a previously reported method^18^ which used the YFull phylogenetic tree v.8.09.^19^

### Ancient DNA authenticity and contamination assessments

Libraries with a low SNP count of fewer than 2,500 are removed from analysis.

We determined authenticity of the ancient DNA for each library by calculating the damage rate of all reads overlapping 1240K SNPs using PMDtools^20^. If the reads of the library have the typical ancient DNA damage of *C*→ *T* deamination in at least 1% in terminal nucleotides (before clipping), the library was determined to be potentially authentic ancient DNA. Libraries above 5,000 SNPs with less than 1% damage or libraries with 2,500-5,000 SNPs with less than 3% damage are removed from analysis.

But just because the ancient DNA is authentic does not mean that there is no contamination. To determine contamination, we used several independent analyses and flagged or removed from analysis any libraries that were determined to be contaminated.

All individuals receive their mtDNA only from their mothers. So all overlapping mtDNA reads are expected to be the same. For individuals with mtDNA coverage greater than 2x, we used contamMix-1.0-12^21^ (includes an option to set pseudo random number seeds) to determine the rate that mitochondrial reads matched the consensus sequence, and flagged as questionable any individuals whose lower bound for the 95% confidence interval for mtDNA heterogeneity (mismatch from consensus) was above 0.05 and we flagged/removed as contaminated when above 0.1.

Males have one X chromosome and so we expect no polymorphisms. For all individuals with a determined genetic sex of male, we calculated the percent of sites on the X chromosome that are polymorphic using hapCon^22^ (restricting to individuals with at least 2000 X chromosome SNPs covered at least once) or ANGSD v0.941-26-g6b5d906^23^ (restricting to individuals with at least 200 X chromosome SNPs covered at least twice). We flagged as questionable any individual whose lower bound of the confidence interval for percent of polymorphic SNPs was at least 1%, and we flagged/removed as contaminated when above 2% for hapConX or ANGSD.

We also flagged individuals with an undetermined genetic sex. An undetermined genetic sex can occur from contamination, but it can also be due to low sequencing coverage.

For individuals that did not pass quality control, and were flagged or assessed to be contaminated, we create a dataset comprising only SNP data from authentic damaged DNA molecules, namely that had deaminated terminal nucleotides. These libraries are denoted with “_d” to indicate that they only include data from reads with ancient DNA damage. Some libraries do not have sufficient coverage from damaged DNA molecules.

### Runs of homozygosity (ROH) detection

For libraries that had a minimum SNP coverage of 300,000 SNPs targeting 1240K sites, we used hapROH^24^ with default settings to infer runs of homozygosity. The total runs of homozygosity (sum of all ROH segments in cM) are reported in Figure 2 for select individuals with a high level of ROH segments greater than 20 cM.

### Relative detection

We employed three methods to infer the existence and degree of familial relationships between every two individuals. The first method used was pairwise mismatch rates between the BAM files of two individuals using the READv2^25^ software package. The second method used was the length and number of identity by descent segments using the ancIBD^26^ software package. The BAMs were first filtered to retain only reads of base quality 30 or greater and mapping quality 30 or greater. VCF files were then generated from the filtered BAMs using the `bcftools mpileup` command. We used the GLIMPSE 1.0^27^ software package and the 1000 Genome Project dataset as the reference panel in order to impute and phase the dataset. We then applied ancIBD with default parameters on our imputed dataset. The third method used was reLD, an internal linkage disequilibrium-based approach that measures, for a pair of individuals, how similar their genotypes are at pairs of loci and how that similarity changes as the genetic distance between loci increases. By using allele-frequency standardized genotypes and the distance-dependent decay in this genotype-similarity correlation, reLD infers the degree and type of relatedness.

### Radiocarbon dating

Radiocarbon dates were generated on the same skeletal elements (tibia from primary BC burial and teeth from all other individuals) as the genetic analysis, at the CSI facility at the University of New Mexico. Bone collagen or tooth dentin was extracted and purified using the modified Longin^28^ method followed by ultrafiltration^29^ or XAD amino acid purification^30^ depending on the preservation of the sample. The skeletal material was suspended in 0.5 N HCl at 5 °C for 24-48 hours until the matrix was demineralized, followed by two rinse cycles of UltraPure 18Ω H_2_O until neutrality. The resulting collagen pseudomorph was gelatinized for 12 hours at 60 °C in 0.01 N HCl and lyophilized for 48 hours. Samples were resuspended in UltraPure 18Ω H_2_O and transferred to pre-cleaned Sartorius Vivaspin 20 ultrafilters (to retain >30 kDa molecular weight gelatin) and centrifuged 6 times. Purified collagen samples were lyophilized for 48 hours and evaluated for crude gelatin yields. Samples with low collagen yields were further processed using XAD-purification to isolate amino acids in the ultrafiltered collagen^31^. The purified collagen was hydrolyzed in 1.5 mL 6 N HCl for 22 hours at 110 °C. The 2 mL collagen hydrolysate solution was transferred to a dropwise syringe and passed through a pretreated Supelco ENVI-Chrom® SPE (Solid Phase Extraction; SigmaAldrich) column and eluted with 10 mL 6 N HCl in a 20 mm culture tube. The purified hydrolysate was heated to 50 °C and dried under UHP N_2_ gas for ∼12 hours. Sample quality was evaluated by % crude gelatin yield, and %C and %N, C:N ratios to assess collagen preservation and contamination^38^. Approximately 0.7 mg of ultrafilter sample and 1.5 mg XAD-purified collagen were weighted into tin capsules and analyzed on a Thermo Scientific Delta V mass spectrometer with a dual inlet and Conflo IV interface connected to a Costech 4010 elemental analyzer (EA). Samples with Atomic C:N ratios between 3.0 and 3.4 progressed to sample combustion and graphitization^32^. Samples were combusted at 900°C in vacuum sealed quartz tubes along with prebaked CuO_2_ and Ag wire.

Resulting CO_2_ was cryogenically purified under vacuum and reduced using a Bosch reaction onto SigmaAldrich Fe powder at 550°C in an H_2_ atmosphere. The resulting Fe/graphite powder was pressed into custom 1mm Al targets and sent the NOSAMS (Woods Hole Oceanographic Institution) facility for AMS measurements.

## Data availability

The aligned sequences are available through the European Nucleotide Archive, accession PRJEBxxxxx [to be made available upon publication].

## Acknowledgments

Special thanks to Peter S. Dunham, Director of the Maya Mountains Archaeological Project (1992-2001) and MKB excavators and surveyors Courtney Laves, Rebecca Hays, and Andrew Kindon. We acknowledge Nasreen Broomandkhoshbacht, Elizabeth Curtis, and Fatma Zalzala for assistance with wet lab procedures at Harvard Medical School. At the University of New Mexico Center for Stable Isotopes, we thank Citlali Tierney, Laura Burkemper, and Viorel Atudorei. At the UNM Laboratory of Human Osteology we are grateful to Paige Lynch. Analysis at Texas Tech University was made possible with the assistance of Dr. Brandon Wagner and students Caitlyn Frazier, Kaitlyn Fulp, Sydney Hendrickson, Jordyn Knysz, Dennise Sanchez, Abigail Sink, Briana N. Smith, and Kaylie Waner. We acknowledge the continuing support of Dr. Melissa Badillo, Direct of the Belize Institute of Archaeology. Special thanks to all the Mopan and Qeq’chi’ communities in southern Belize that bound the Bladen Nature Reserve.

## Author contributions

E.S.B., D.R., and K.M.P. conceptualized the study. D.R. and K.M.P. supervised and acquired funding. E.S.B. formally analyzed the genetic data. E.S.B., D.R., and K.M.P. were involved in the genetic data investigation. K.C., E.C., T.F., L.I., J.K., A.M.L., N.M., J.O., I.P., and J.N.W. performed laboratory procedures under N.R.s supervision. A.K., M.M., M.N., G.S., and S.M., processed genetic data and performed bioinformatic curation. A.A., R.F., I.L., I.O., and K.S. provided further genetic processing. B.D., D.J.K, J.M., M.R., and K.M.P. provided archaeological resources and support. K.M.P. curated archaeological data. E.M., A.C.N., E.R., and M.W. performed osteological data analyses. A.E.T. conducted geospatial analysis. E.S.B., N.C.N., and K.M.P. performed chronological data analyses. E.S.B., D.R. and K.M.P. wrote the original draft. E.S.B., A.C.N., E.M., N.C.N., E.E.R., A.E.T., and K.M.P. contributed to the original supplementary information. All coauthors reviewed and edited the paper.

## Supplementary Information

The online version contains supplementary material available at [to be made available upon publication].

**Peer review information** to be added after review.

**Reprints and permissions information** is available at [to be made available upon publication].

**Extended Data Figure 1.**
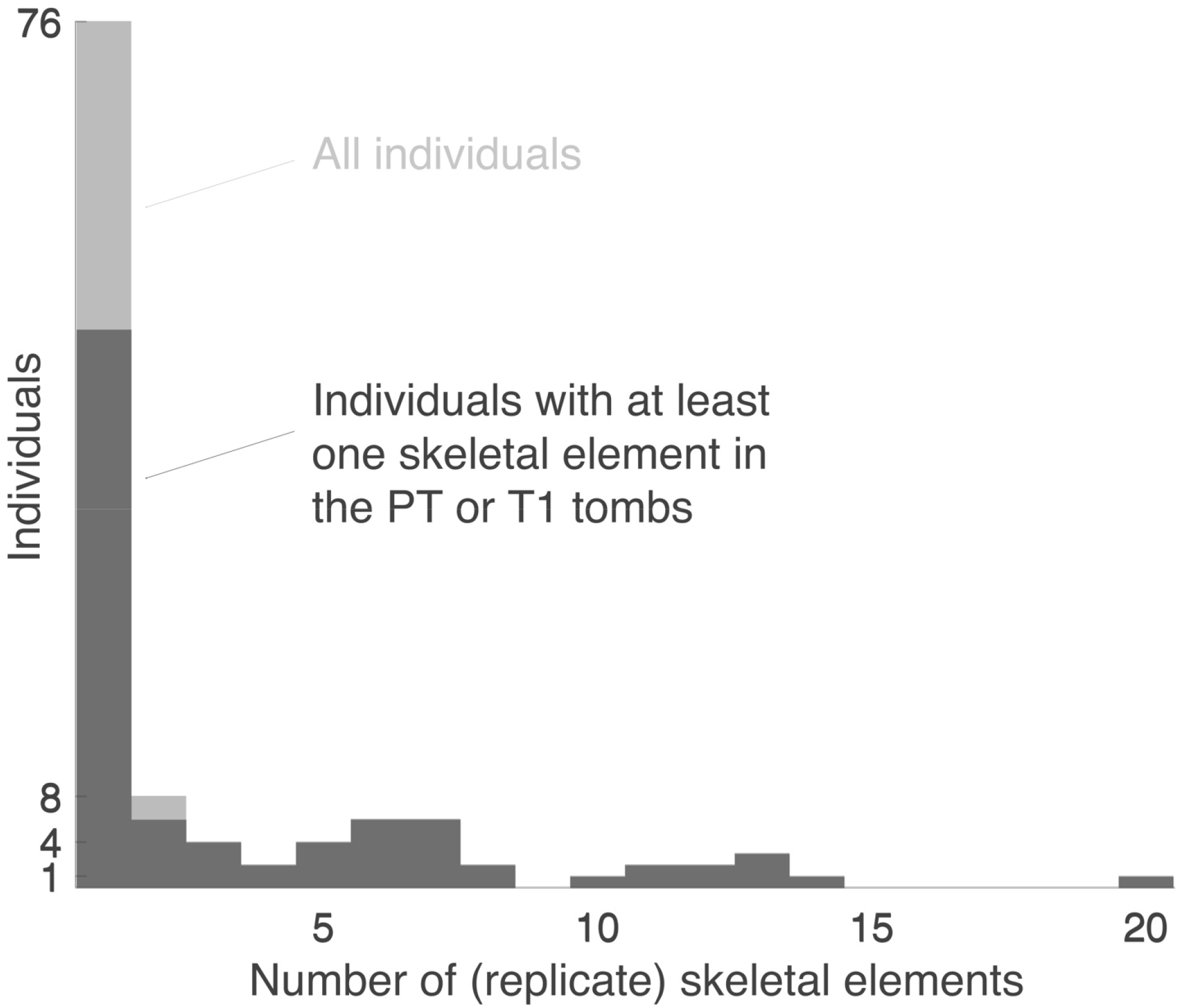
Histogram of the number of replicate skeletal elements recovered. The darker gray overlay are samples where at least one skeletal element was found at Muklebal Tzul. Replicates are considered if they are within site or between-site.

**Extended Data Table 1.**
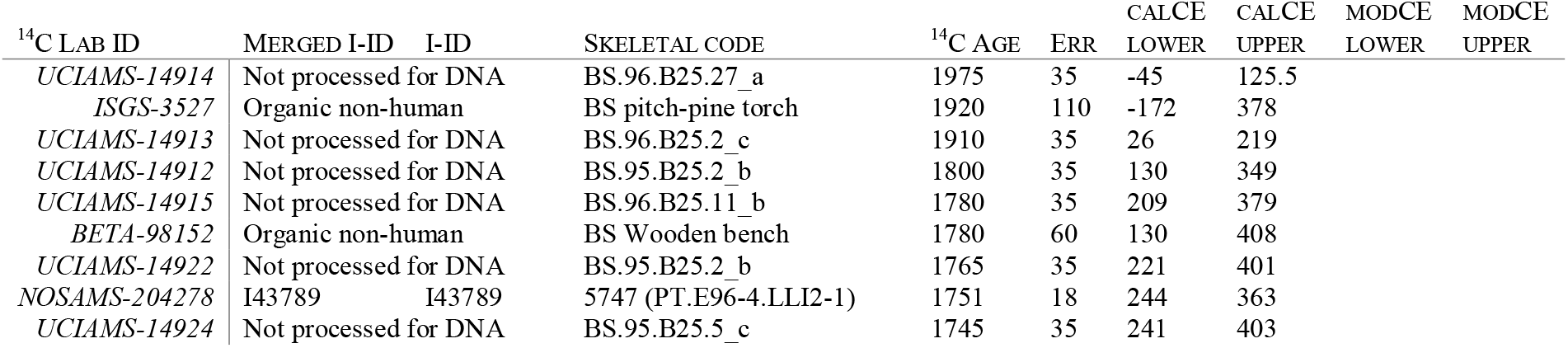
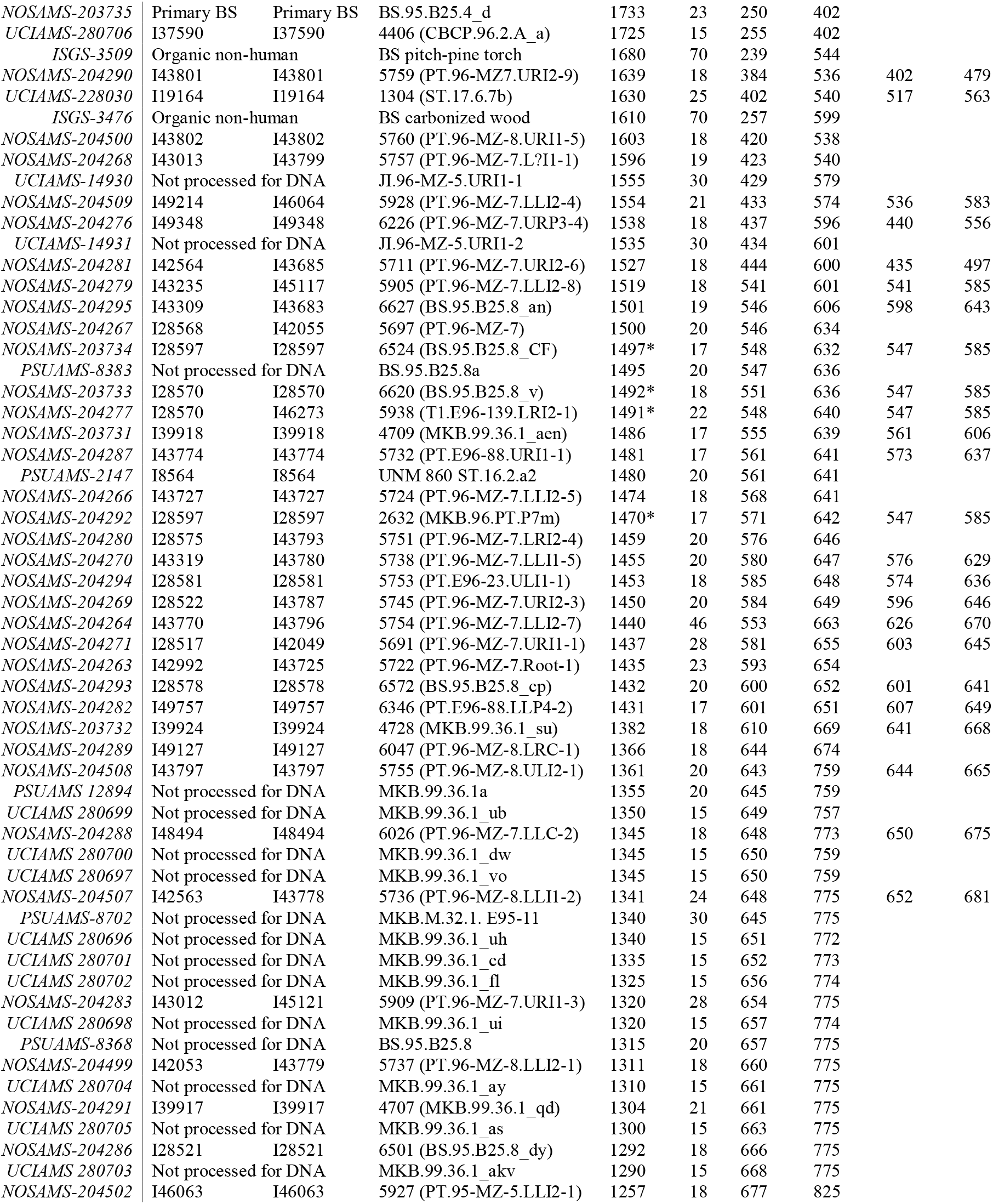
Uncalibrated and calibrated ^14^C dates and modeled dates. See SI Discussion 4, for further information on OxCal calibration and modeling. The only individuals listed here are those for whom we have direct ^14^C dates. All other individuals with modeled dates are included in Table S4.2. For individuals for whom DNA was processed, both the individual I-ID and skeletal code for the sample that underwent radiocarbon dating are listed as well as the merged I-ID for easy identification in Figure 2 in the case of replicates. The calibrated dates lower and upper 2σ (95.4% CI) bounds are in columns calCE lower and calCE upper, respectively. The modeled dates lower and upper 2σ (95.4% CI) bounds are in columns modCE lower and modCE upper. ^14^C dates with asterisks were first combined before modeling.

