## Supplemental Information for "Buried in two places: Lineages from elite Maya tombs also found in distant caves"

^1^Department of Human Evolutionary Biology, Harvard University, Cambridge, MA, USA. ^2^Broad Institute of MIT and Harvard, Cambridge, MA, USA. ^3^Ya’axché Conservation Trust, Punta Gorda, Toledo District, Belize. ^4^Department of Anthropology, University of California Santa Barbara, Santa Barbara, CA, USA. ^5^Uchben’kaj Kin Ajaw Association, Santa Cruz, Toledo District, Belize. ^6^Department of Anthropology, University of Texas at San Antonio, San Antonio, Texas, USA. ^7^Department of Anthropology, University of New Mexico, Albuquerque, NM, USA. ^8^Center for Stable Isotopes, University of New Mexico, Albuquerque, NM, USA. ^9^Department of Sociology, Anthropology, and Social Work, Texas Tech University, Lubbock, TX, USA. ^10^Department of Archaeology and History, University of Exeter, Exeter, UK. ^11^Department of Geography and the Environment, The University of Texas at Austin, Austin, TX, USA. ^12^Department of Genetics, Harvard Medical School, Boston, MA, USA. ^13^Howard Hughes Medical Institute, Harvard Medical School, Boston, MA, USA. ^14^Department of Zoology and Animal Cell Biology, University of the Basque Country EHU, Vitoria-Gasteiz, Spain. ^15^Ikerbasque-Basque Foundation of Science, Bilbao, Spain. *Corresponding authors. ‡These authors contributed equally.

### 1. Site Descriptions

#### Muklebal Tzul

Excavations at Muklebal Tzul were conducted between 1995-2000 as part of the Maya Mountains Archaeological Project (MMAP) that was directed by PS Dunham under permits issued by the Belize Department of Archaeology (DOA) and Forest Department (FD) and which co-lead author KMP was a project member. The two elite tombs, Plaza Tomb (PT) and Tomb 1 (T1) in the elite residential compound were excavated in 1995 and 1996 as salvage operations, as their entrances were exposed and they could have been subjected to looting. Human remains from Muklebal Tzul PT and T1 were exported by MMAP in the 1990s to the osteology laboratory of Dr. Frank and Julie Saul, and later transferred to KMP at the University of New Mexico in 2020. Tombs in settlement groups West 5 Str. 36, West 5 Str. 3, West 1 Str. 4, and West 1 Str. 6. are included in this study. This is just a small proportion of the 49 tombs documented in settlements at the site. Human remains from the settlement tombs at Muklebal Tzul were curated in Belize and exported under permits to KMP in 2020-2022 for further analyses, including osteological analysis, ^14^C dating, and genetic analysis.

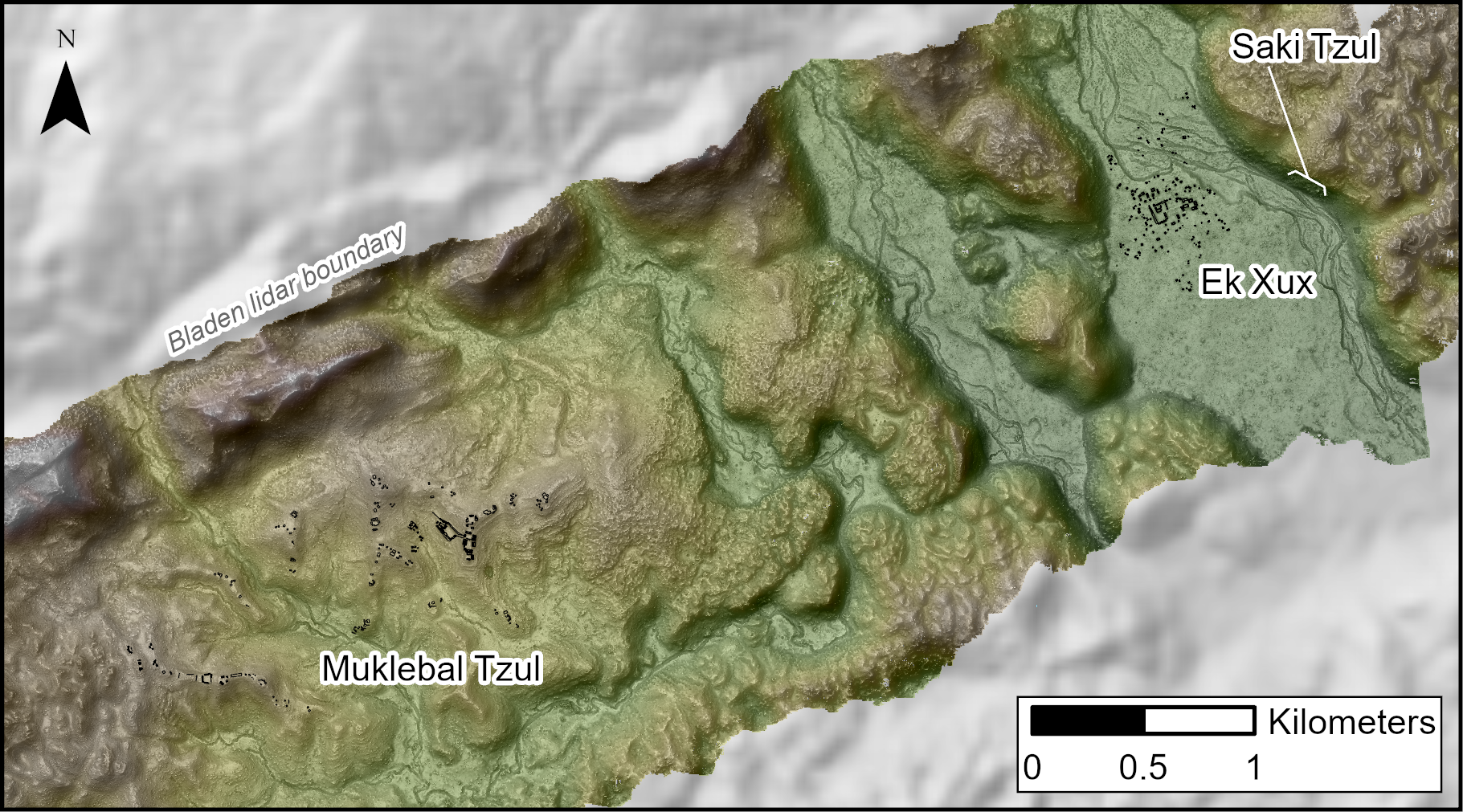

*Figure S1.1 Surface map of the upper Bladen Branch valley with three of the sites discussed in the text. Lidar derived map is a linear burn blend of a Digital Elevation Model (DEM) at 80% transparency and Visualization for Archaeological Topography (VAT) at 60% transparency.*

Muklebal Tzul is in the uppermost of the major valleys on the Bladen Branch of the Monkey River. The valley is also the largest in the Bladen Branch, measuring 4.5 x 2 km, with elevations ranging between 300-450 m (Figure S1.1). The Bladen Branch river is largely an underground system in this area, and its surface channel, on the south margin of the valley, is seasonally intermittent. The river channel enters and exits the valley through headwall caves. There is no permanent water within 1.25 km of the core. The topography of the valley system containing Muklebal Tzul does not consist of an alluvial bottomland but is a ridge-and-valley consisting of ridges and rolling hills composed of interbedded-limestone interspersed with swales with deep soils creating a high-productivity arable landscape. Most of the ridges were leveled and expanded (cut-and-filled) prior to the construction of the site core and settlements on the flattened areas (Figure S1.2). The layers of bedded limestone are well suited for use as architectural construction material.

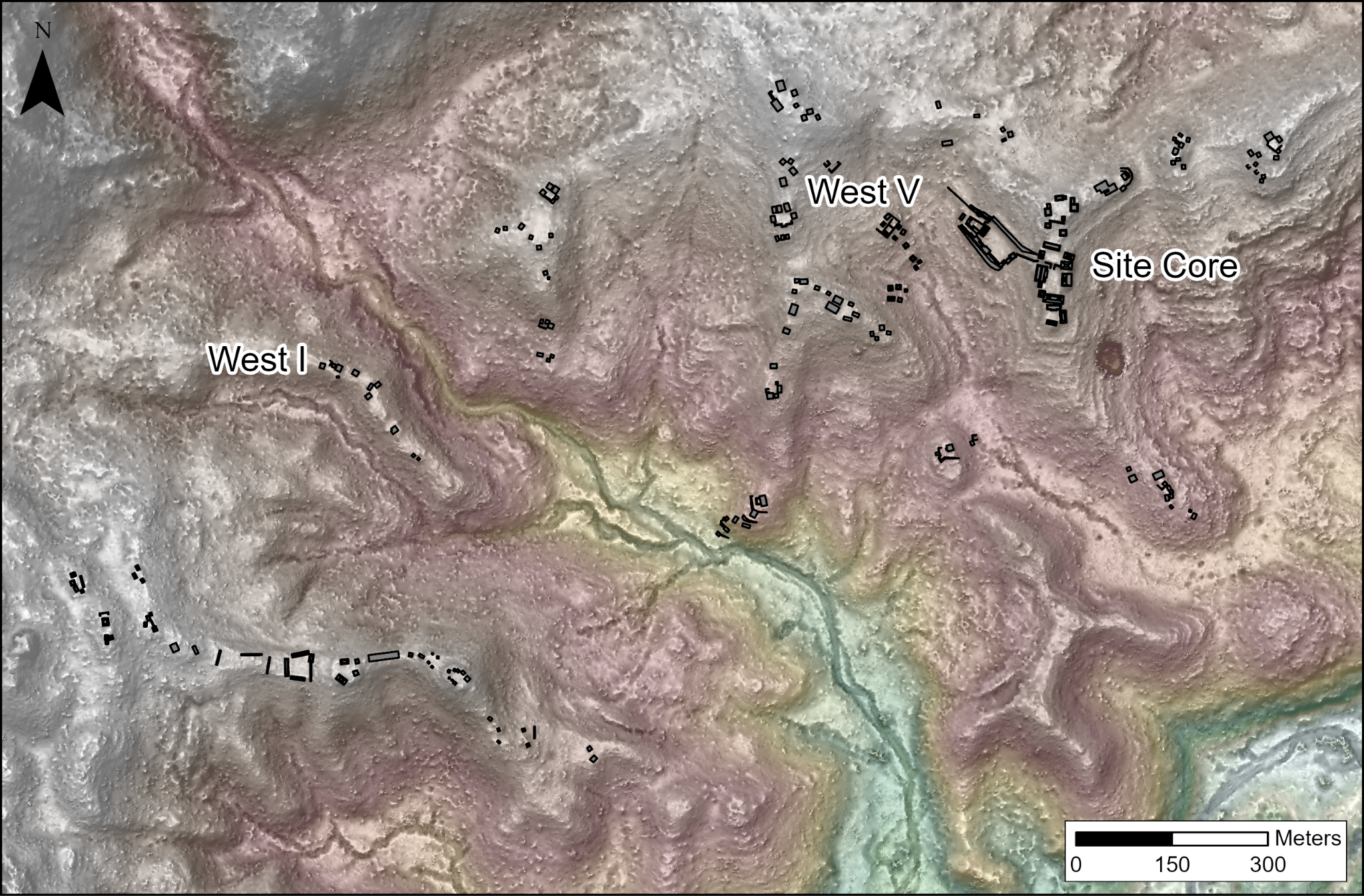

*Figure S1.2 Muklebal Tzul core and mapped settlements showing local topographic variation.*

The most striking cultural feature at Muklebal Tzul is a steep, modified ridge that cuts across the landscape (Figure S1.2). The site core was built on top of this ridge, and it is surrounded by numerous smaller ridges with narrow spines located to the west, east, and south. These ridges are separated by small seasonal creeks. The Muklebal Tzul site core is arranged on two levels (Figure S1.3). The upper level is along the ridge and consists of four conjoined residential plaza groups (Groups 1-4). The lower level is below the ridge and consists of a large platform with political and ceremonial architecture (Groups 5-6).

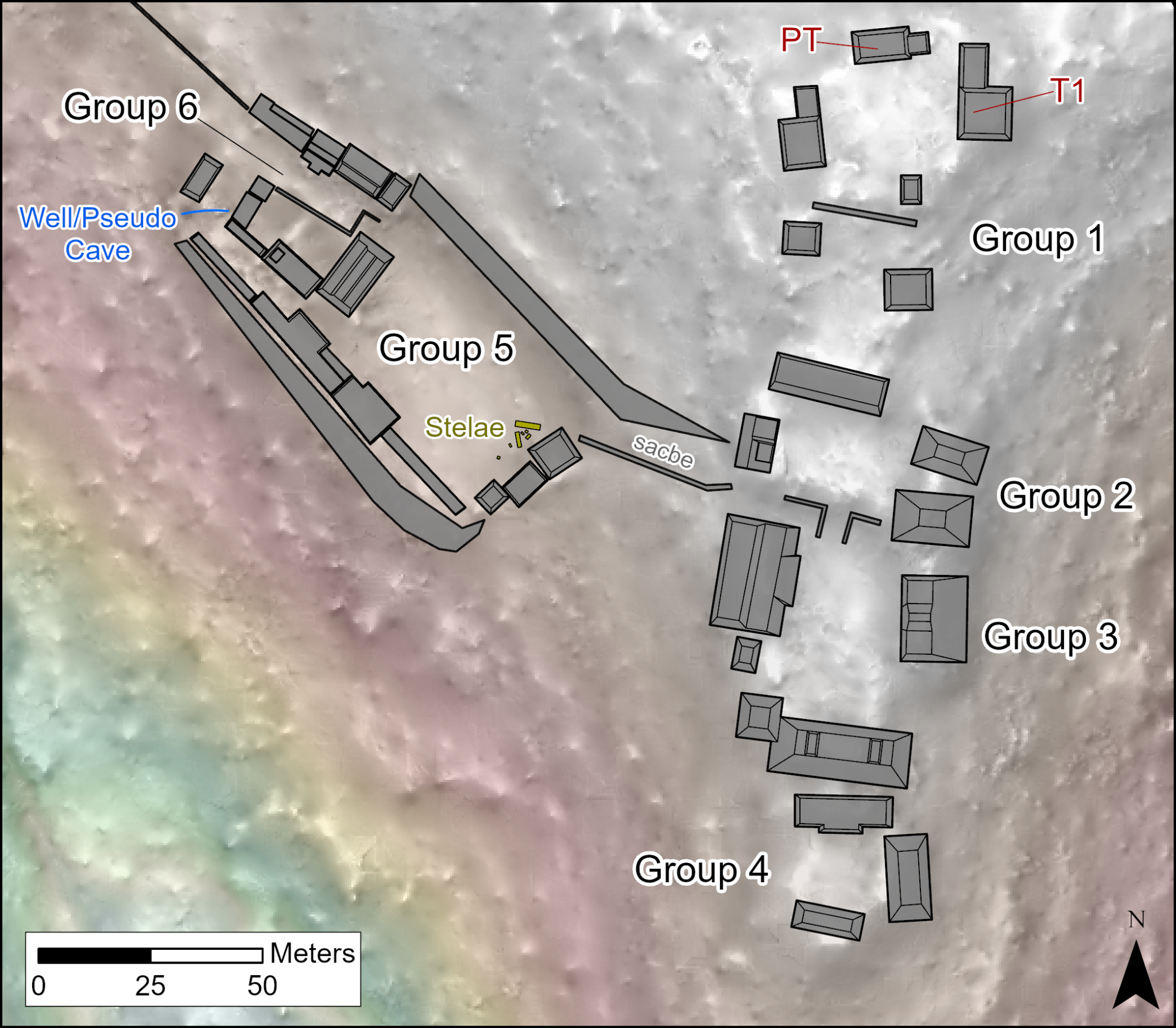

*Figure S1.3. Core areas of Muklebal Tzul include the elite residential plazas and the ceremonial and political groups.*

The upper-level groups occupy the highest points at the site and are arranged in a linear fashion along the spine of the ridge in such a way that the northwestern plaza group (Group 1) is the highest, and the other three (Groups 2-4), moving to the southeast, are progressively lower. These were likely residential compounds and ceremonial areas under the control of the elite members of the site. The location atop the ridge and lack of easy access to these groups suggests that this area was restricted. Group 1 was the location of Plaza Tomb and Tomb 1.

The lower level is a massive 4720m^2^ platform constructed on a flattened hillside 11 m below the ridge that supports the conjoined elite residential groups. This lower platform is connected to the plaza Group 3 by a 36.5 m long 12 m wide parapeted causeway leading from the southeast corner of the platform up the slope to Group 3. The summit of the great platform was divided into two main plazas in antiquity. The larger of the two plazas is a stelae plaza that occupied the southeastern two-thirds of the platform. This plaza is bounded on the northeast by the upslope wall, and by structures on the northwest, southwest and southeast. Directly in front of the easternmost and tallest structure on the platform, are six stelae and what may be one hemispherical altar. All of the stelae were broken, and only three of the six stela bodies remain, although all the bases appear to have retained their original placements. The smaller of the two plazas on the platform (Group 6) is crowded with structures and built features. This plaza is bounded by several low buildings built in its center that effectively worked to bisect it into two separate spaces: a smaller, interior plaza surrounded by a narrow C shaped plaza. The division of space suggests a much more restricted access to the activities carried out in this plaza area compared to the stelae plaza. The exterior plaza slopes steeply, indicating that access to both the smaller and larger plazas was primarily from the larger stelae plaza. It also contains a water-bearing well that is a 13-meter-long horizontal passage leading to a spring (Figure S1.4). It is accessed by descending a three-step stairway that leads to a small plaster-lined basin or pool^1^. The passage also has many features commonly associated with natural caves: a dark zone, an underground stream, and a twilight activity area. However, the volume of water it produces, 2.6 l/minute combined with its location under a large platform that houses a stelae complex, argues that the water from the well was used for ceremonial or other restricted activities or for the personal use of the residents of the elite residential groups.

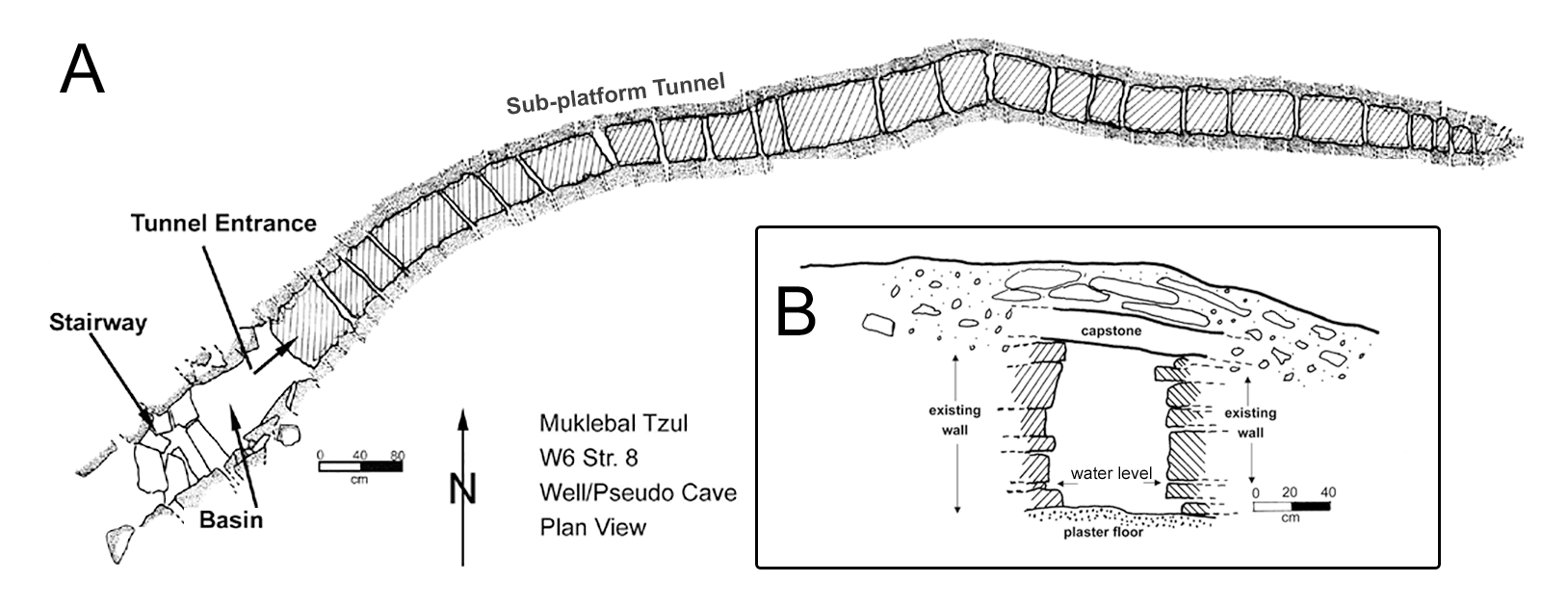

*Figure S1.4. A. Plan-view of MKB well/pseudocave well. B. Entrance doorway to the well.*

A total of 211 structures have been mapped in settlement zones at Muklebal Tzul during a partial survey made between 1996-2000. Residential buildings at the site are clustered into groups and these groups are further arranged into larger neighborhoods across the spines of the low ridges that dissect the valley. The ridges are distributed across much of the Muklebal Tzul valley and settlements are replicated in this fashion across the site, running down the main spines of ridges with a wide array of raised and parapeted causeways linking some of the more formal groups. The acquisition of lidar data covering the Muklebal Tzul Valley in 2019 allows for a more complete visualization of modifications to the landscape and the extent of settlements.

Typical architectural groups have structures arranged linearly along the spine of major ridges. Architecture is generally built along ridgetops, and as most ridges are narrow, the architecture extends from side to side, covering the entire width (Fig S1.2). Architectural arrangements are usually more formal on the upper reaches of ridges, with buildings concentrated on the highest points. This pattern replicates the arrangement found in the site core in the settlement groups.

Building platforms in the Muklebal Tzul settlements are typically small, ranging from 50 cm to several meters in height. Like elsewhere in the region, cut-stone facades were built along many of the slopes and hillsides, and these stone-covered natural rises are incorporated into the architecture to give it a grand appearance. All of the buildings excavated to date were built in single events. Construction at Muklebal Tzul is largely cut-block limestone masonry. Blocks were fairly easily fashioned from naturally occurring bedded limestone that is found on the ground and eroding from hillside surfaces throughout the site. This easy to access bedded limestone likely helped facilitate the construction of numerous tombs across the site. All settlement groups that were documented during survey have tombs of varying sizes built into residential platforms. Excavations indicated that the tombs were built at the time that buildings were constructed. It appears that masonry forming the walls of the tombs was constructed on the leveled bedrock ridges. The platform structures were then constructed around the tombs, with additional cut block facades on the platform exteriors and filled in with debris and rocks. Access to the interior tomb chambers was either through horizontal or vertical entrances, some of which were sealed.

To date, a total of 37 sub structural tombs were documented at the site (Figure S1.5). Except for one tomb that was found during an excavation, all tombs were documented based on the presence of collapsed roof slabs or visible entrance shafts or doorways^2^. There are likely many more that have not collapsed or that have entrances that are obscured. The vaulted construction of the platforms and tombs has likely preserved a large number that have not been identified. Less than 10 structures at the site have been looted, likely owing to the remote location, and those that have been looted all have tombs.

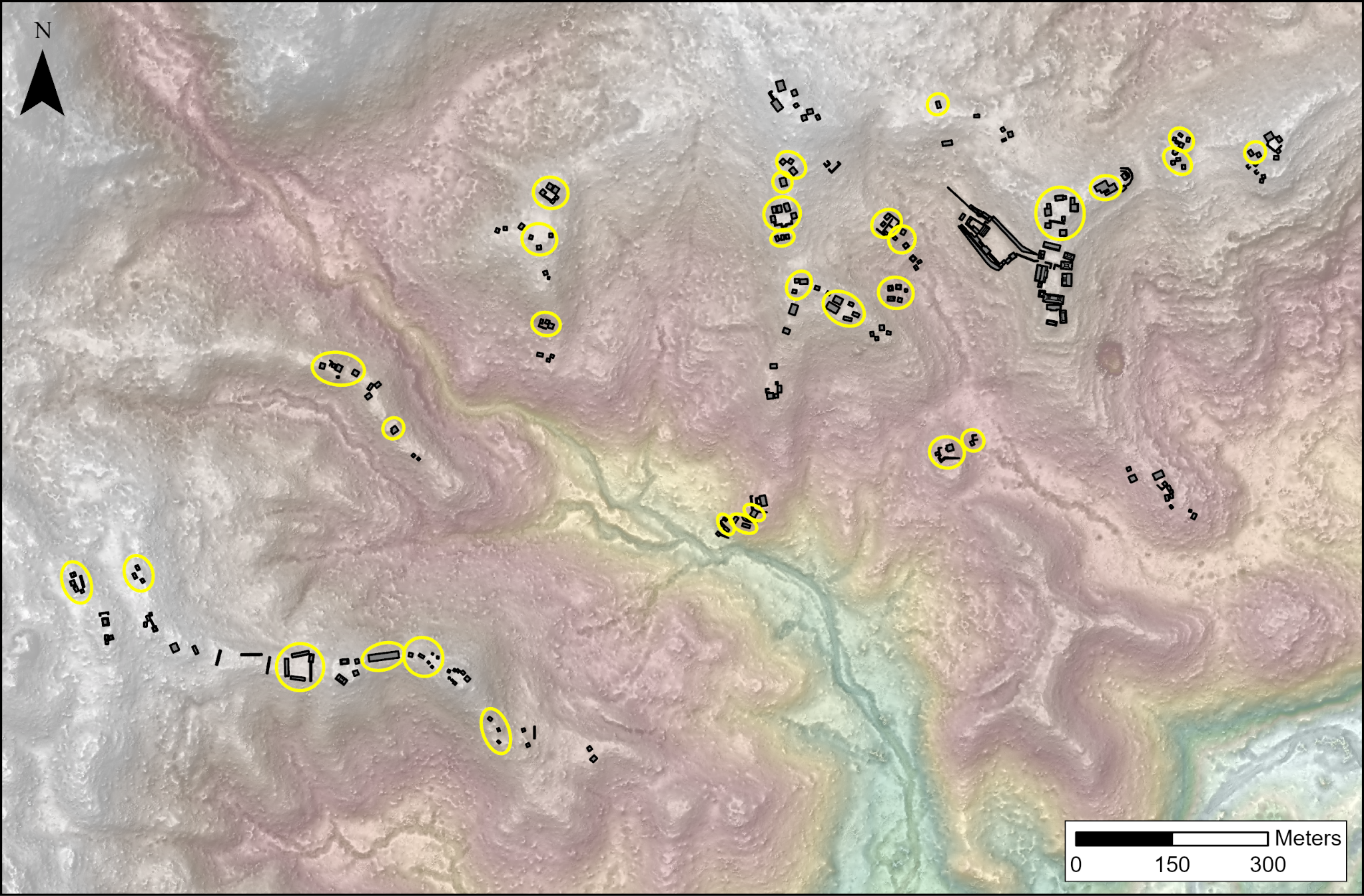

*Figure S1.5. Muklebal Tzul settlements. Yellow ovals mark 31 groups containing 37 previously documented tombs.*

A total of 13 tombs were excavated between 1995-2000, including the two tombs in the site core elite residential area (PT and T1), and tombs at the W1, W3, and W5 settlement groups (Figure S1.2). Only roughly half (6 of 13) of the excavated tombs contained skeletal material at the time when they were documented. Several have deposits of artifacts that suggest a pattern of deposition similar to that found in caves. Three (one excavated) settlement tombs and PT were natural fissures in the limestone around which tomb walls and platforms were constructed. Two fissure tombs at Plaza Tomb and at W5, Str. 36 contained complex deposits of disarticulated remains of multiple individuals. PT and T1 tombs have experienced repeated flooding due to seasonal rainfall and disturbance from tree roots that destabilized the architecture. By the time of excavation, the burials in both tombs were disarticulated and embedded in sediment.

##### i. Plaza Tomb

PT is a single chamber tomb located in Group 1 of Muklebal Tzul and was excavated by the MMAP during the 1995 and 1996 field seasons. Fieldnotes describe it as a walled modification to a natural crevice or fissure in the elite residential Group 1 overlooking the ceremonial site core. The long axis of the tomb was oriented along the east/west fissure, and it was constructed of rectangular stone block walls covered with thick capstones which had partially collapsed into the chamber (Figure S1.6). The chamber dimensions are about 1 m in height, 2.5 m long and about 0.9 to 1.2 m wide. The eastern wall is described as a rock slab in the field notes, possibly exposed bedrock. A low stone bench or step runs along the western wall. The tomb was constructed on bedrock which forms the floor of the chamber and has a corbeled ceiling. The east half had soil above the bedrock. A natural fissure in the floor is where the majority of human remains were recovered. The human remains were recovered throughout the chamber. There was one set of articulated remains described by excavators as a partially flexed burial. They note that the bones of the legs and feet were in the correct anatomical position. Otherwise, they emphasize the high volume of disarticulated bone. The field drawings indicated that at least two skulls were found in close proximity along the southern wall of the tomb and long bones were generally oriented east/west. Cultural materials found in the tomb include stingray spines, several carved shell ornaments, manos and metates used as grinding stones for corn and cacao, obsidian blades, shell and ceramic earspools, a carved bone “scoop”, a bone ring, a biface fragment, jade bead, as well as sherds. Subsequent analysis of recovered ceramics showed most of the typeable ceramics date to the Late Classic (500-800).

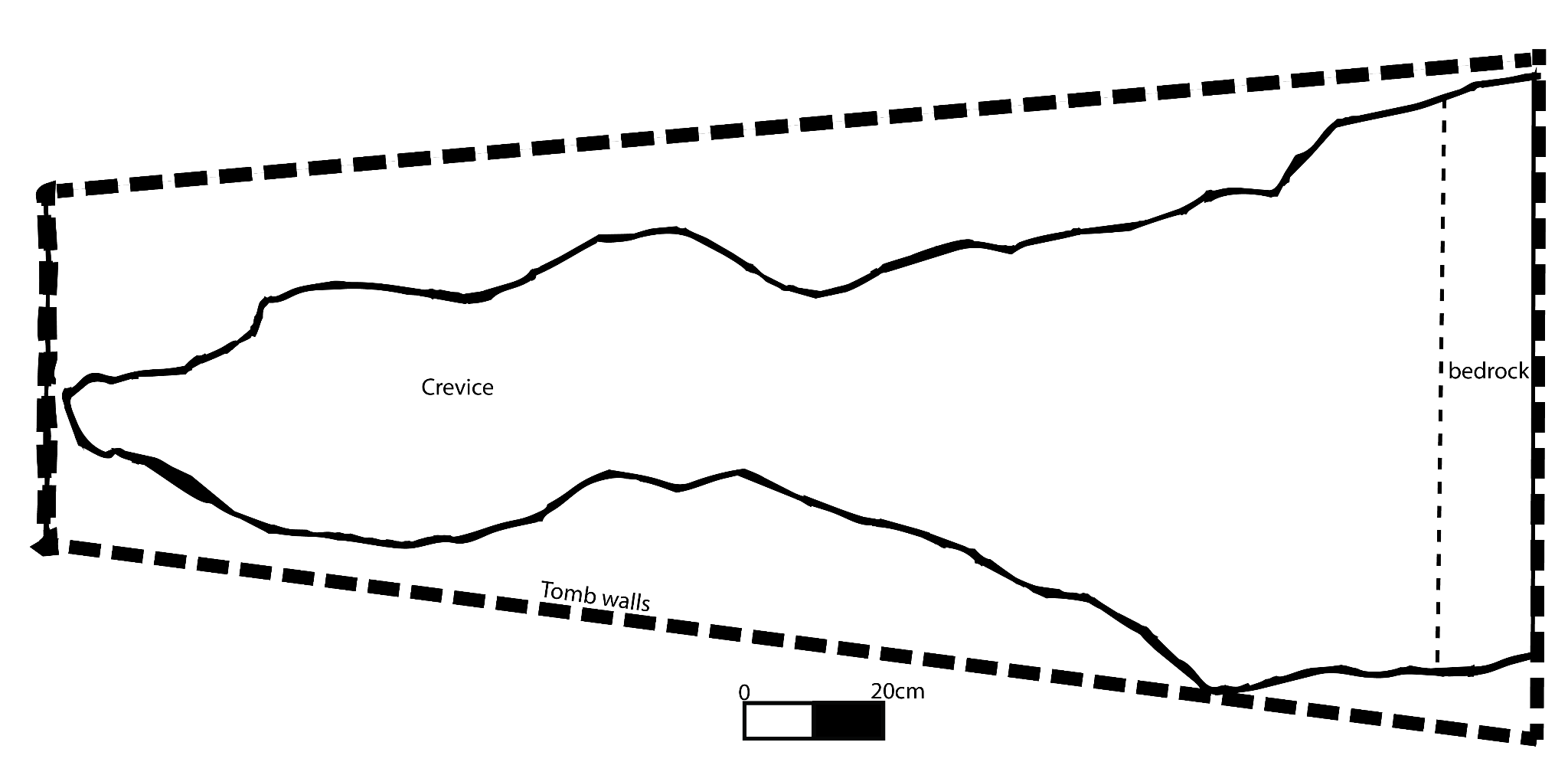

*Figure S1.6. Plan-view of the Plaza Tomb located in Group 1, the elite residential compound.*

Osteological analysis estimates the minimum number of individuals at PT as 41 based on bones and 28 based on teeth, with 8 subadults based on bone and 5 subadults based on teeth. While we expected that any one individual might have as many as 32 teeth in the tomb, it was surprising that we still identified a minimum of 88 distinct individuals at PT with genetic. Analysis of the all human remains from Muklebal Tzul was based on the physical remains and on limited fieldnotes from the MMAP excavations from 1995-2000. Additionally, some materials were lost or damaged following a hurricane that destroyed parts of the MMAP field camp in Belize in 2001.

PT Excavation notes and bag labels indicate a largely commingled context (Online Table 6). The notes do not indicate organization in terms of bone deposition (i.e. gathering of skulls or long bones in one part of the tomb).The human remains recovered include 505 teeth and hundreds of identifiable skeletal elements, with many bags of fragments too small or eroded to identify. The assemblage consists of bones from the axial and appendicular skeleton, including numerous small bones of the hands and feet. While degree of articulation is unknown, the elements present suggest that Plaza Tomb was the primary interment location for these individuals. The remains of 34 adult individuals were recovered based on the proximal one-third and midshaft of the right femur (Table 1). Skeletal pathology was evident on long bones in the form of fracture callouses and periosteal reaction, both active and healed at the time of death. There was also extensive dental pathology in the form of antemortem tooth loss. There were seven instances of dental modifications with anterior teeth inlaid with greenstone and eight teeth with the incisal edge filed.

Skeletal elements were recovered from eight subadults; five isolated nonadult teeth were also recovered. The nonadult age ranges include late childhood of ~7-10 years and adolescence of 11-18 years (Schaefer, Black and Scheuer 2009). All nonadult skeletal fragments were extremely small and only identified by the degree of sealing of the epiphyseal surfaces on damaged bone. There was no element repetition with the nonadult remains. No pathology was evident on the nonadult bones or dentition.

##### ii. Tomb 1

T1 in Group 1 of the site core consists of two chambers that were stacked vertically (Figure S1.7). The tomb was excavated during the 1995 and 1996 field seasons of the MMAP. The tomb was constructed of stacked stone blocks with a tunnel entrance from the plaza. The tunnel partially collapsed and was thus blocked. The capstones of the upper chamber of the tomb, however, had also partially collapsed, providing direct access to the upper chamber. The upper chamber was about 1.5 m by 2 m with a flat capstone ceiling. A doorway on the west side of the upper chamber leads to a passage with a ledge and three stone steps leading down to the lower chamber. The passage connecting the chambers was 2.17 m in height. The lower chamber was slightly larger than the upper chamber. The lower chamber measured approximately 2.6 m north/south and was wider along the eastern wall (2 m) than the western wall (1.6 m). The lower chamber was topped by a vaulted ceiling that rose 33 cm above the top of the supporting walls. The maximum height of the lower chamber was 1.6 m in the center of the chamber. Artifacts were recovered from both chambers and the shaft, including ceramics, shells, stone tool debris, a mano, greenstone or jadite, and faunal bone. An obsidian blade was found in the upper chamber and a fragment of a reddish chert blade and a mano were found in the shaft. Of significance is the high volume of lithic debris, waste from tool making, from the upper chamber described in the field notes, which is consistent with lithic deposits found in tomb contexts throughout the Maya lowlands^3^. The excavators frequently mention finding shells in both chambers, some of which were carved. Subsequent ceramic analysis describes a possible basal flange bowl, typically associated with the Early Classic (250-500). Other ceramics including a red-slipped jar rim are indicative of the Late Classic.

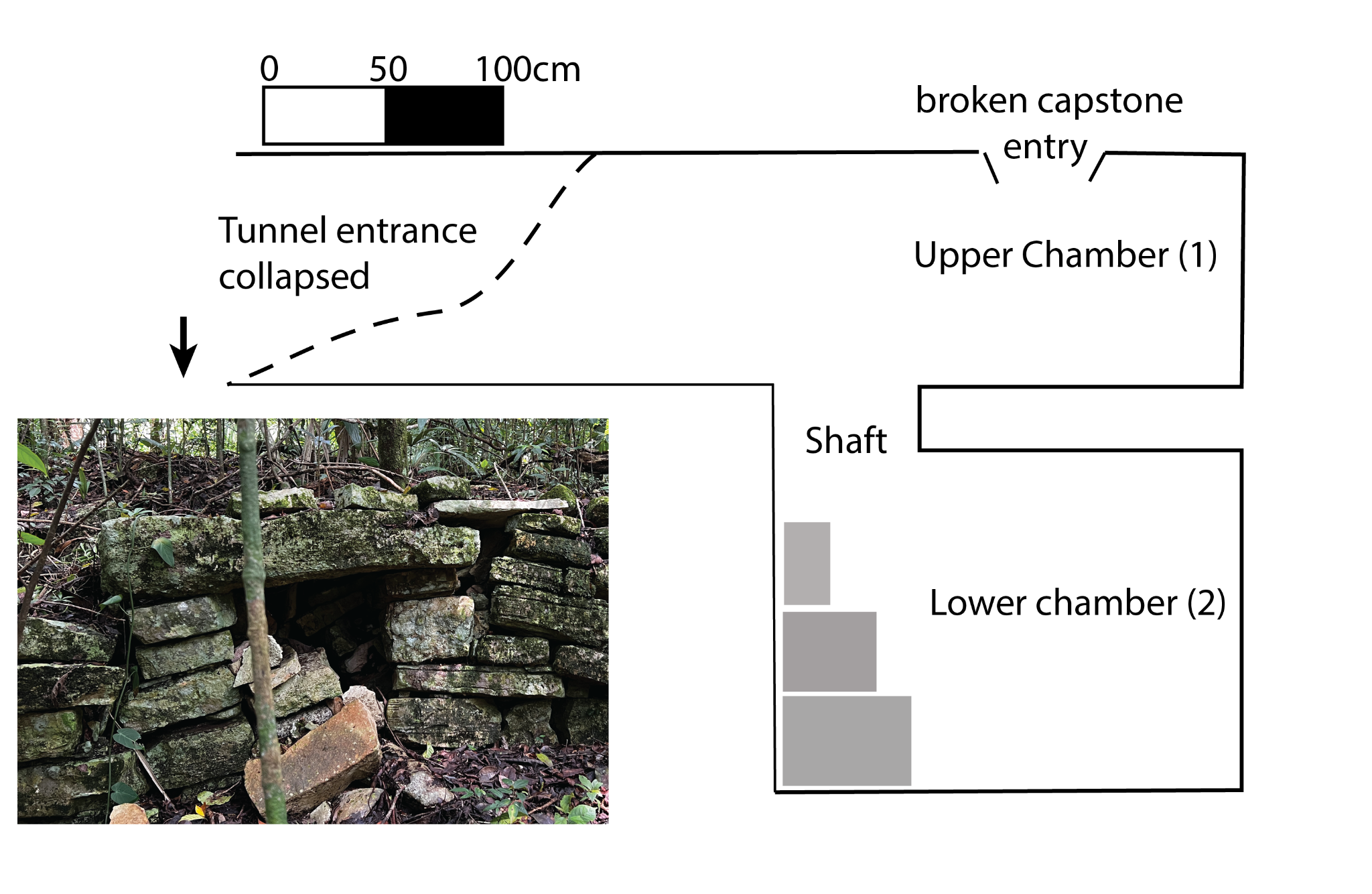

*Figure S1.7. Profile view of Tomb 1 with inset photograph of the lintel-capped entry tunnel.*

Human bones were found throughout Tomb 1 (Online Table 7). Available excavation documents describe excavations in both the upper and lower chambers, however they do not describe where the bones were located within the upper chamber. Notes on the lower chamber state that the majority of human bone was found in the center of the chamber. Bag labels for all skeletal elements from tomb 1 include “Chamber ?”, “Posthole”, “Chamber 2” (lower chamber), “Lower Chamber”, and “Shaft”. There are only minor macroscopic differences in taphonomy between these contexts. There are two teeth, mandibular molars, that were labelled “shaft”, indicating they were found in the passageway between chambers. “Posthole” refers to a series of circular holes dug in the lower chamber to support the ceiling, which is also partially the floor of the upper chamber. Field notes report human bone as well as greenstone recovered from the postholes.

The human remains recovered include seven isolated adult teeth and 57 identifiable bones, with numerous smaller bone fragments. Both the axial and appendicular skeleton are represented, including smaller bones of the hands and feet. There were very few bones from the cranium, thorax (ribs and sternum), or vertebral column. The total MNI from Tomb 1 is 11 (Table 1) based on seven isolated permanent teeth, three left mandibles, and one nonadult based on mixed dentition in alveolar bone (Online Table 7). A fragment of right maxilla was recovered with mixed dentition of a nonadult aged approximately 10-12 years at death^4^. There were no nonadult postcranial bones or epiphyses recovered. The elements present suggest that this was the primary place of interment for these individuals.

##### iii. West 5 Structure 36 tomb

W5 S36 is located in a non-elite settlement area 150 meters from the core of MKB (Figure S1.8). It was excavated in 1998 and 2000. This structure contained a tomb that was repeatedly used as an ossuary located in a non-elite settlement area <100 m meters from the core of MKB. Like tomb PT, W5 S36 follows a natural limestone fissure that the Maya artificially deepened by building low walls capped by large (20+ cm thick) limestone slab capstones (Figure S1.8). When first identified in 1998, most of the ceiling had collapsed, but showed evidence of being vaulted. Human remains were discovered immediately beneath the soil surface within the tomb. During the 1998 excavation, at least 3 individuals were identified based on fragmentary cranial deposits associated with disarticulated remains. In 1999, deeper excavations resulted in the recovery of over 1,000 mostly partial skeletal elements and an MNI of at least 20 based on dentition (Table 1). Artifacts recovered include ceramics, lithics, shell ear flares, carved bone, quartz crystals, hematite, and marine cowrie shells. The excavators note that several of the artifacts may be directly associated with specific individuals. The shell ear flares were found next to the articulated individual in the eastern half of the unit. There was also an upturned plate placed on the abdomen of this individual.

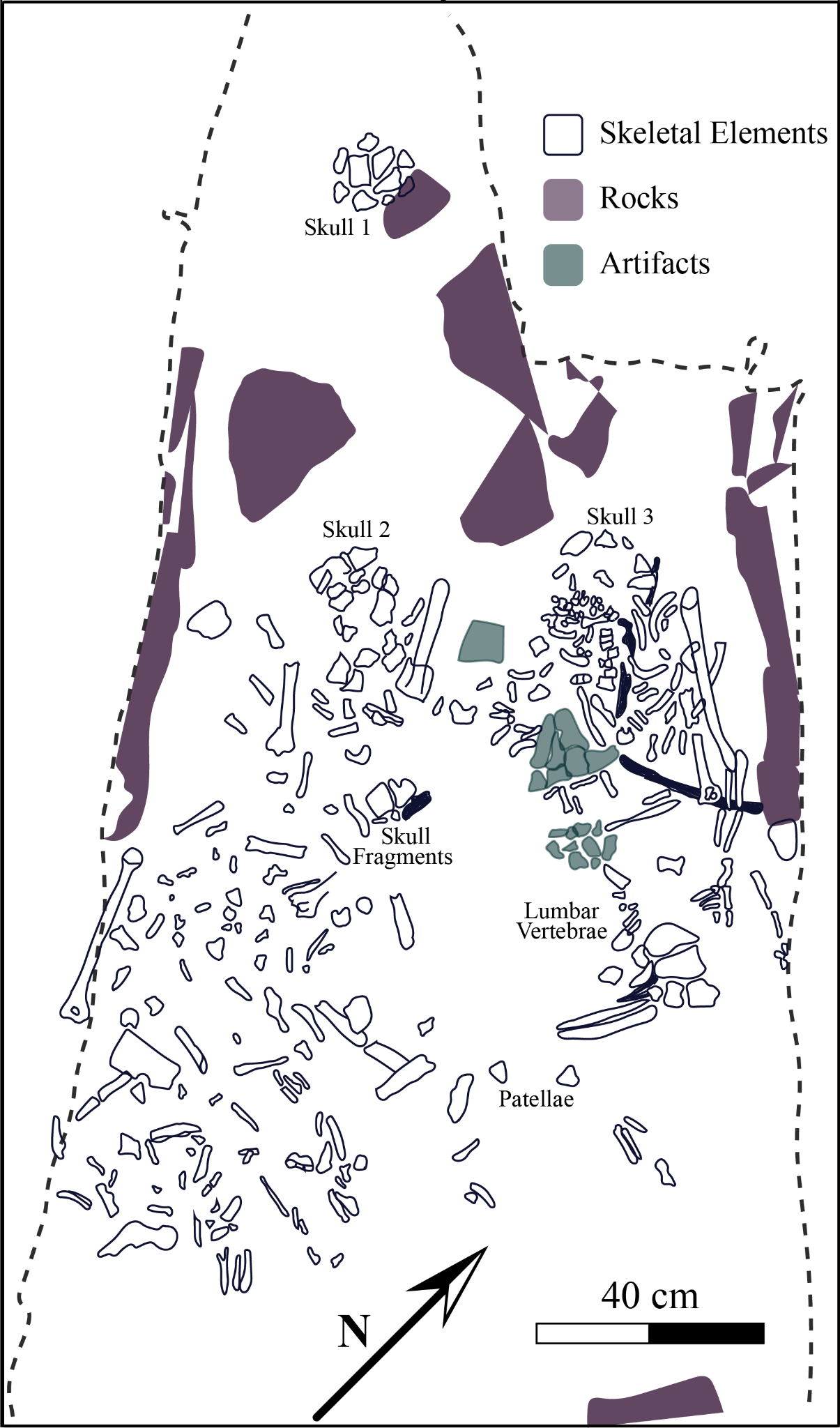

*Figure S1.8. Plan-view of the W5 S36 tomb*

Two concentrations of skull fragments were initially found in the northern and southern half of the tomb (Online Table 8). A third skull fragment was noted near the center and may have been the latest whole individual interred in this tomb since this was the only individual that was partially articulated. It is represented by the skull, vertebrae, ribs, and left arm. All other elements were disarticulated and comingled. It is possible that the MKB community revisited this tomb for additional interments, pushing aside existing remains in preparation for another individual.

Osteological analyses revealed that more than 1100 skeletal elements were included in W5 S36 (Online Table 8). The excavation notes indicate that clusters of bones were scattered throughout the tomb. Although all regions of the skeleton are represented, elements of the thorax (ribs, sternum), spine (vertebrae), and head (cranial vault, face, mandible) appeared with low frequency. A total of 373 teeth, including 2 deciduous teeth and 2 developing teeth, are present, representing at least 19 adults (based on replicate mandibular left canines) and 2 nonadults (based on replicate mandibular right canines in different stages of development: estimated 5-6 years and 10-12 years^4^, based on replicate mandibular left canines. All other possible permanent teeth in the dental arcade are present, with frequencies ranging between 3-18. Bones of the hand were the second most prevalent in W5 S36, totaling 415 elements. At least 12 individuals are represented based on 12 duplicates each of left scaphoids, left trapezia, right trapezoids, and right capitates. Most bones throughout the tomb had evidence of at least minor osteophytic lipping and/or remodeled periosteal reaction on the cortical surface, suggesting some individuals had arthritis, healed infections, or both.

##### iv. West 1 Structures 4 and 6

The West 1 Group consists of 21 structures which range from 0.5 to 2 meters in height spread across six plazas along a ridge (Figure S1.2). In total, two tombs with burials were excavated in structures 1 and 6. The tombs in W1 were oval and had less developed vaulting than those located in the site core but each contained low passages with linteled entrances, plaster floors, and ceramic offerings.

The W1 S4 tomb had partially collapsed when it was first documented in 1998 and excavations were completed in 1999. The original entrance to the tomb is located near the center of the northern basal edge of the mound (Figure S1.9). A thick, roughly rectangular limestone slab blocked the original tunnel-entrance that was still intact on the interior but the tunnel leading outward had collapsed. The entrance itself was rectangular and was capped with a limestone lintel. The tomb interior measures 222 cm NS x 192 cm EW and is oval with curved corners and has a simple vaulted roof. While the northern, western, and southern walls of the tomb are well-preserved, portions of the eastern wall had collapsed in antiquity, partially covering the original entrance to the tomb. Disarticulated skeletal material was recovered primarily from beneath this collapsed wall. Artifacts were excavated from all areas of the tomb including the top and side entrances. These include ceramic sherds, lithics, and faunal bone and shells. The structure was looted between the two seasons of excavation but much of the material seems to be *in situ* or was moved by natural taphonomic processes. Excavation notes point to several exotic items including a ceramic whistle, carved shell, obsidian blades, and a ceramic torch holder. Several unit stamped sherds or partial vessels were found throughout the excavation. In Southern Belize, unit stamping is assigned to the Late Classic as identified at the site of Lubaantun^5^.

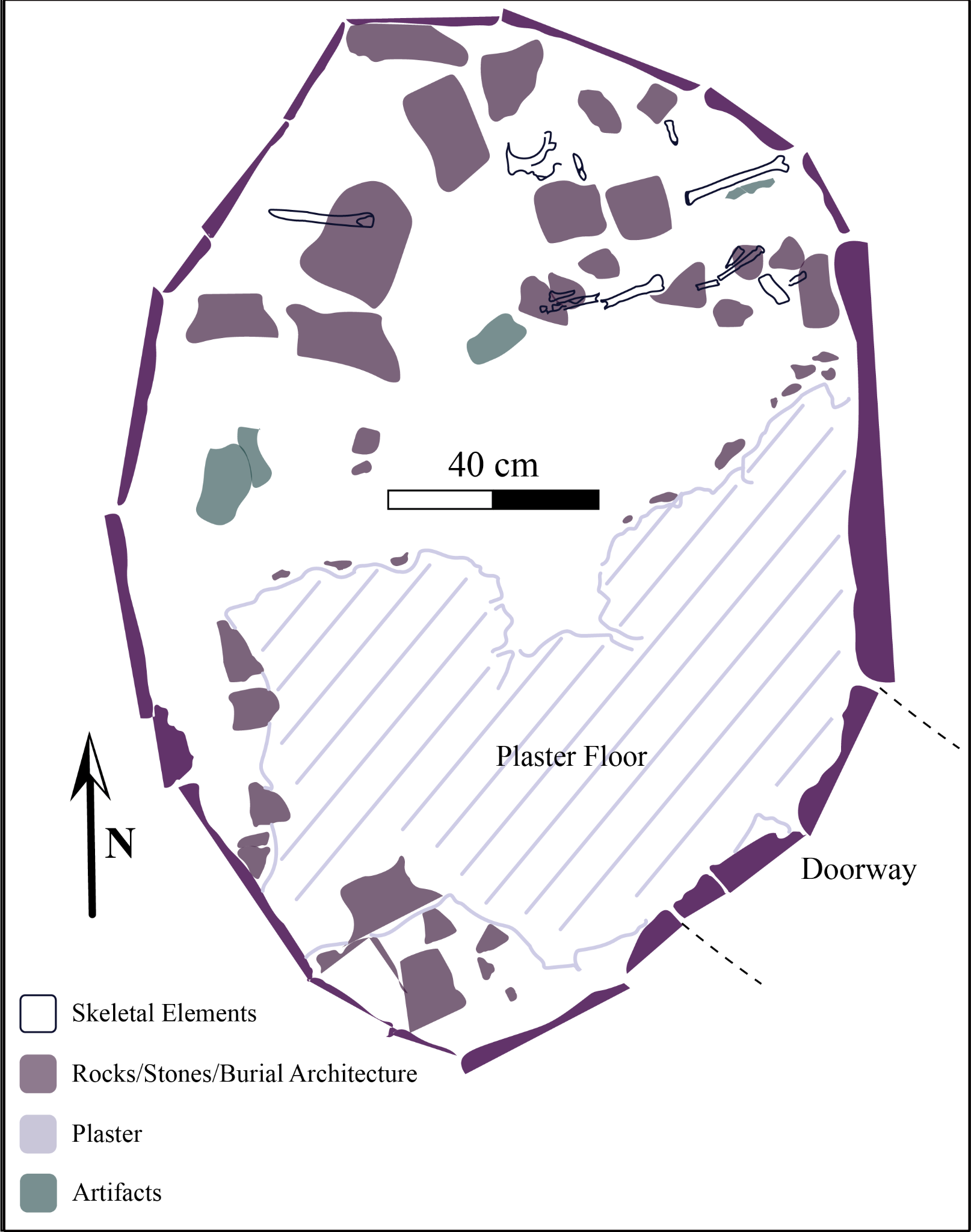

*Figure S1.9.Plan-view of the W1 S4 tomb*

W1 S4 is represented by a small, highly fragmentary assemblage recovered from multiple contexts (Online Table 9), likely reflecting post-depositional mixing, as skeletal elements refit across contexts and/or repeated disturbance. A minimum of two individuals is indicated by the duplication of humeral elements, right temporal bones, and occipitals across contexts, although each context (i.e., Skull 2, Skull 3, and Skull 4) independently suggests a minimum of one individual. A total of 58 skeletal elements were recorded in the inventory, including over 369 bone fragments.

The assemblage is dominated by fragmentary cranial bones, long bones, ribs, and small elements of the hands and feet, with preservation marked by soft, exfoliated cortical bone and extensive taphonomic damage consistent with burial in clay-rich soils and exposure to environmental disturbance. Field notes and excavation records indicate that some materials derive from collapse and secondary deposition within the tomb entrance and passageway, while others represent a partially articulated burial in the northern portion of the chamber or a dispersed bone scatter lacking clear articulation.

Several elements provide limited insight into the individuals' biological profiles. A relatively complete mandible (Skull 3) retains most of its dentition in occlusion, with the left lower first molar (LLM1) exhibiting complete alveolar resorption and the right lower third molar (LRM3) absent due to postmortem breakage between the body and ramus. The right molars exhibit substantial wear, likely reflecting compensatory mastication following abscessing of the left first molar, which obscures some morphological traits. Minimal caries and calculus are present, permitting limited morphological assessment. The mental eminence, the bony, triangular projection at the front-center of the mandible, is scored as 4^6^, suggesting a probable male. In contrast, cranial features associated with Skull 2, including an orbital margin score of 3 and a mastoid process score of 4, indicate an indeterminate sex. Overall, sex estimation is constrained by the absence of diagnostic os coxae features and the fragmentary nature of the assemblage.

All individuals are adults based on epiphyseal fusion and dental eruption. No additional skeletal features are sufficiently preserved to refine age estimates. Evidence of periosteal reaction is present on several long-bone fragments, however, its extent and etiology are difficult to assess due to poor preservation.

W1 S6 tomb measured 200 cm NS by 160 cm EW. This tomb was also oval, with no defined corners (Figure S1.10). The tomb would have been accessed via the northern wall of the structure, which contained a sill and doorjambs but no lintel. Human bone material was identified in the northern quadrant of the tomb interior. At least three distinct individuals were documented during the excavations. In the southwestern quadrant of the tomb, a tooth cache was identified that was not associated with any maxilla or mandible fragments.

Artifacts recovered include ceramics, freshwater shell, obsidian and quartz fragments but no chert lithics. A jade pendant depicting either a face with a feathered headdress or a monkey was found *in situ* in association with the primary burial. Drilled holes on the back of the pendant indicate that it was meant to be suspended without disrupting the carved design on the front. Another piece of jade was also found in close proximity with perforations though its exact function could not be determined. A ceramic whistle in the shape of a bird, a small marine shell, obsidian, quartz, and two polychrome sherds were also recovered from the base of the tomb.

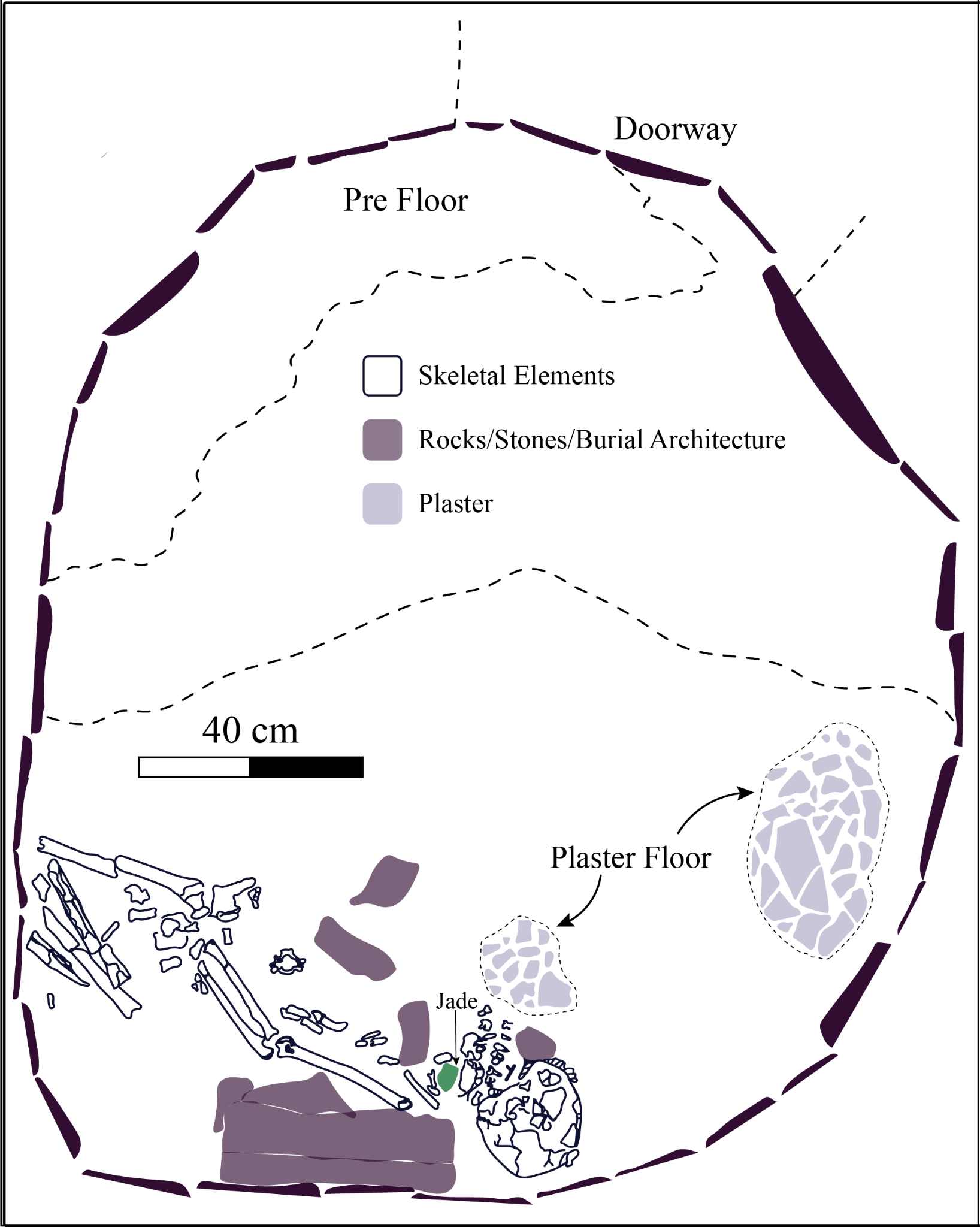

*Figure S1.10. Plan-view of the W1 S6 tomb*

The W1 S6 tomb was divided into two burial lots: MKB.98.6.2 and MKB.98.6.3a-b. MKB.96.6.2 consists of the remains of at least two individuals of unknown sex. The primary individual is a young to middle adult, and approximately 10% of their skeleton is present, although it is poorly preserved. All elements are fragmented and affected by insect/root damage. The outer layer of bones with thick cortical bone is soft (as seen in other contexts at Muklebal Tzul), making observations of pathology difficult. Remains of a commingled individual are present, in the form of charred long bone fragments (radius, possible metacarpal) that may belong to a subadult based on their size. For the primary individual, sex could not be estimated since neither the cranium nor much of the pelvis was preserved. However, partial reconstruction of an unsided tibial diaphysis showed that the individual was very gracile, which is possibly a feminine characteristic. Age was estimated using traditional methods^6^ and transition analysis^7^ using the right pubic symphysis. Age estimates (in years) range from 25+, 25-60, and 19.9-48.4, with median ages of 38, 35, and 29.1 respectively. Pathological changes were difficult to observe in many instances due to taphonomic damage. However, it was evident that the unsided tibial diaphyseal fragments (around midshaft) had health issues reflected in a porous periosteal reaction around the entire shaft. Similarly, both femora have healing, striated periosteal reaction on their diaphyses, which is also circumferential. The reaction becomes microporous on the proximal diaphyses Together these changes are possibly indicative of an infection.

This context consists of the remains of at least two adults of unknown sex. They are adults based on complete dental and skeletal development. Sex cannot be estimated since the pelves were not preserved, and the crania is too damaged by taphonomy. The original excavation notes document two separate crania that were recovered (with associated dentition), but the postcranial remains were comingled. It was not possible to separate the postcranial remains by individual, so they are denoted by lot as MKB.98.6.3, with the crania and teeth separated into individuals 3a and 3b. Between both individuals, approximately 25% of the skeleton is present, and is primarily represented by the two crania, long bones, and miscellaneous fragments. The remains are poorly preserved. Exfoliation and root/insect damage is so extensive that the bones are soft such that brush strokes from cleaning are evident in the cortical bone. Both crania were partially reconstructed to document evidence of cranial modification. The distal diaphyses of a left MC2 and an unnumbered metacarpal have well-healed, poorly aligned fractures of unknown types. For MKB.98.6.3b reconstruction of the cranial vault showed artificial cranial modification in the form of (parallelo-frontal-occipital) tabular modification with oblique pressure, causing bilobate expansion. Pressure was centered at the squamous portion of the occipital bone, with only one pad impression evident, which is at midline. There is depression at lamdba, but impressions of binding are not observed. The anterior aspect of the cranium was not preserved.

MKB.98.6.3a was described in the notes as a possible tooth cache, but may be the teeth of a single individual. Dental development is complete. Labial drilling is present on the ULC, and has a caries where the inlay would be. Individual 3a may have retained their ULM2 crown since this tooth has the same crown morphology as ULM1 but is significantly smaller, and all upper left permanent molars are accounted for. This tooth would have been in the place of ULP4 (which is absent in this individual). The maxillary anterior crowns are broken, rendering most pathological observations (wear, caries) unknown. Only one other caries is present in this dentition, in the interproximal surface of ULM3. A light amount of calculus is present on almost all teeth. Abscesses are unobservable since all teeth were found without associated alveolar bone. Teeth exhibit a moderate amount of attrition, with scores ranging from 2-4 for anterior teeth, and 2-6 for molar quadrants. The only enamel defect observable is a linear enamel hypoplasia on the LLC 2.5 mm from the CEJ, indicating developmental stress during tooth formation.

#### Bats’ub Cave

BS was initially documented as a salvage operation in 1995 after it was discovered by members of a British volunteer organization assisting the government of Belize with a forestry project. The entrance to the cave is 1.2 meters wide, 1.5 meters high and faces north (Figure S1.11). Entry requires a 1.5m vertical descent into the entry room which is 11 meters long and slightly over three meters at its widest point. The entry room periodically floods. Cultural materials were found inside an interior burial chamber that measures 4.3 meters by 2.5 meters and has a very low ceiling making it impossible for an adult to stand upright. The two rooms were separated by a floor-to-ceiling masonry wall that was plastered with a mix that contains fragments of bone. Throughout the interior chamber, charcoal and burnt wood fragments were collected. The ceiling and portions of the walls were coated with heavy layers of carbon, consistent with both burning of torches for light and the burning of incense. Radiocarbon dates of wood fragments from pieces of pine (*Pinus* caribaea) torches have a 2σ (95.4%) probability that the deposit dates to between 134-615 calCE, median probabilities within twenty-five years of 410 CE.

Limited testing in 1995 determined that there were subsurface human remains. In 1996, a team returned to the cave by helicopter for ten days to conduct excavations and to survey the surrounding area for signs of surface settlement. The team did not find any substantial settlement. Inside the cave, the excavations revealed a partially articulated sub-surface skeleton as well as associated grave goods. Removal of the thin surface soil revealed a degraded gravely layer beneath which a single person was interred. This adult burial was found in a shallow trench dug along the north wall of the inner chamber at a depth of approximately 20cm with the body placed on an unprepared clay floor. The sloping of the floor resulted in differential preservation of the skeletal material; the lower extremities were fairly well preserved but the upper portion of the body, including the pelvis, upper extremities, and vertebral column, was resting on moist clay, fostering decay. The cranium had been removed from the body in antiquity and in its place was a partial ceramic bowl containing a single jade bead. This may be analogous to Classic period Maya funerary practices in which jade beads were placed at the mouths of deceased elites^8^, and may be related to historical Maya beliefs that precious stones placed in burials of rulers capture the soul-breath of the deceased^9^.Cranial fragments (presumably from this individual) were found to the left of the medial plane of the body above the pelvis. No vertebrae were found near the cranium. The spine, which was resting on and slightly embedded in the moist clay floor, was complete but unrecoverable. Most of it had been replaced by clay leaving only a thin veneer of skeletal material. Caches of teeth were found both in the area of the pelvis and also at the top of the spinal column around the cervical vertebrae. In the area of the neck, seventeen beads, thirteen of which are jadeite and four of which are hematite, were recovered along with two carved shell disks. The disks are undecorated except for a scalloped border design. Loose soil and clay surrounded the burial. Above the burial was a thin layer of crushed limestone and when intially investigated bones were protruding from this surface. Embedded in this layer, just above the pelvis, was a small inverted bowl containing five cacao seeds. In the front room of the cave, at the base of the wall separating the two chambers, a location was identified atop a flat rock where travertine was crushed, presumably for mortar that was used to seal the wall separating the chambers. The wall itself was constructed of rough-cut and uncut limestone blocks neatly stacked. The exterior of this wall was sealed with a mixture of crushed travertine, mud, ash and, in at least one place, bone. Considerable care had been taken to ensure that all cracks between the blocks were thoroughly sealed, greatly aiding the long-term preservation of organic materials in the cave.

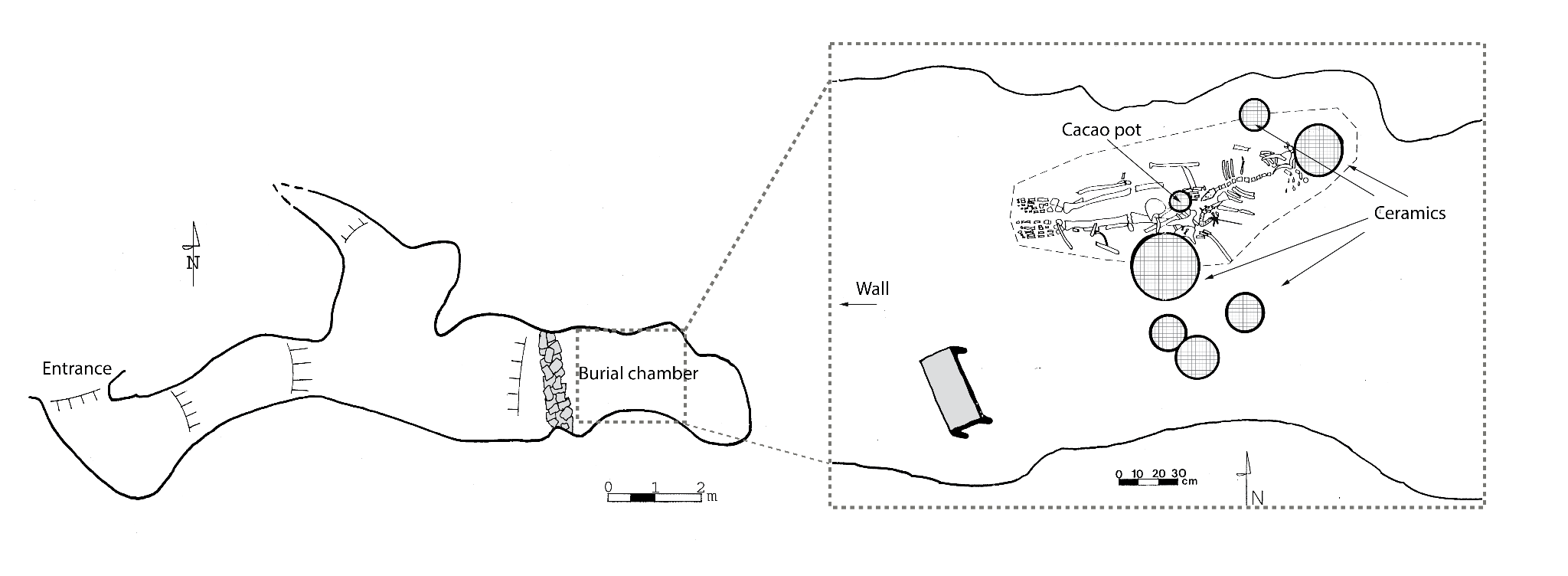

*Figure S1.11. Plan-view of Bats’ub Cave (left) with expanded view of the burial chamber (right).*

Sometime following the interment of the primary individual, grave goods were arranged around and on the burial surface. The elaborate funerary assemblage from BS indicates that the primary interment was an important individual in life and in death (Figure S1.12a-e). The ceramics are Early Classic and other parts of the funerary deposit also date to the Early Classic Period. The most elaborate is a black and red on orange bowl that had a characteristic basal-flange (Figure S1.12b), which was a feature of pottery known to date to 250-500 CE^10,11^, and depicted a scene from the Maya underworld. The exterior shows two individuals—or possibly the same individual repeated on both sides of the vessel—lying face up, with protruding tongues and eyes as narrow slits, details suggestive of a death pose. Either the head appears to be placed in the body’s lap or the character appears to be missing a torso. Their feet rest on what is likely a sacred bundle^12^ which is described elsewhere as bloodletter bundles, which could be a collection of stingray spines wrapped in cloth^13^. The interior surface of the vessel is decorated with a mythical feathered-serpent creature emerging from an object that could be a torch handle or a stingray spine (a bloodletting implement). Torches, stingray spines, and feathered serpents are all themes associated with an underworld setting. One vessel is a jar with a simple cacao bean applique (Figure S1.12a). Two of the remaining vessels depict unusual creatures (Figure S1.12c-d). One has an anthropomorphic face with a shape resembling a turtle body; the other either has an anthropomorphic or an animal face that cannot be confidently identified.

A small rosewood (*Dalbergia* stevensonii) stool is a unique piece of grave furniture (Figure S1.12e). If stools were placed in other burials in Mesoamerica, we have no evidence for them. The stool measures 35 × 17 × 8 cm and was carved from a single piece of wood. An AMS date from the stool places it between AD 85 and 410 (2σ) with a median probability of AD 243. If it was earlier than the primary burial it may indicate that the stool was an heirloom or that it was carved from the heartwood of a rosewood tree.

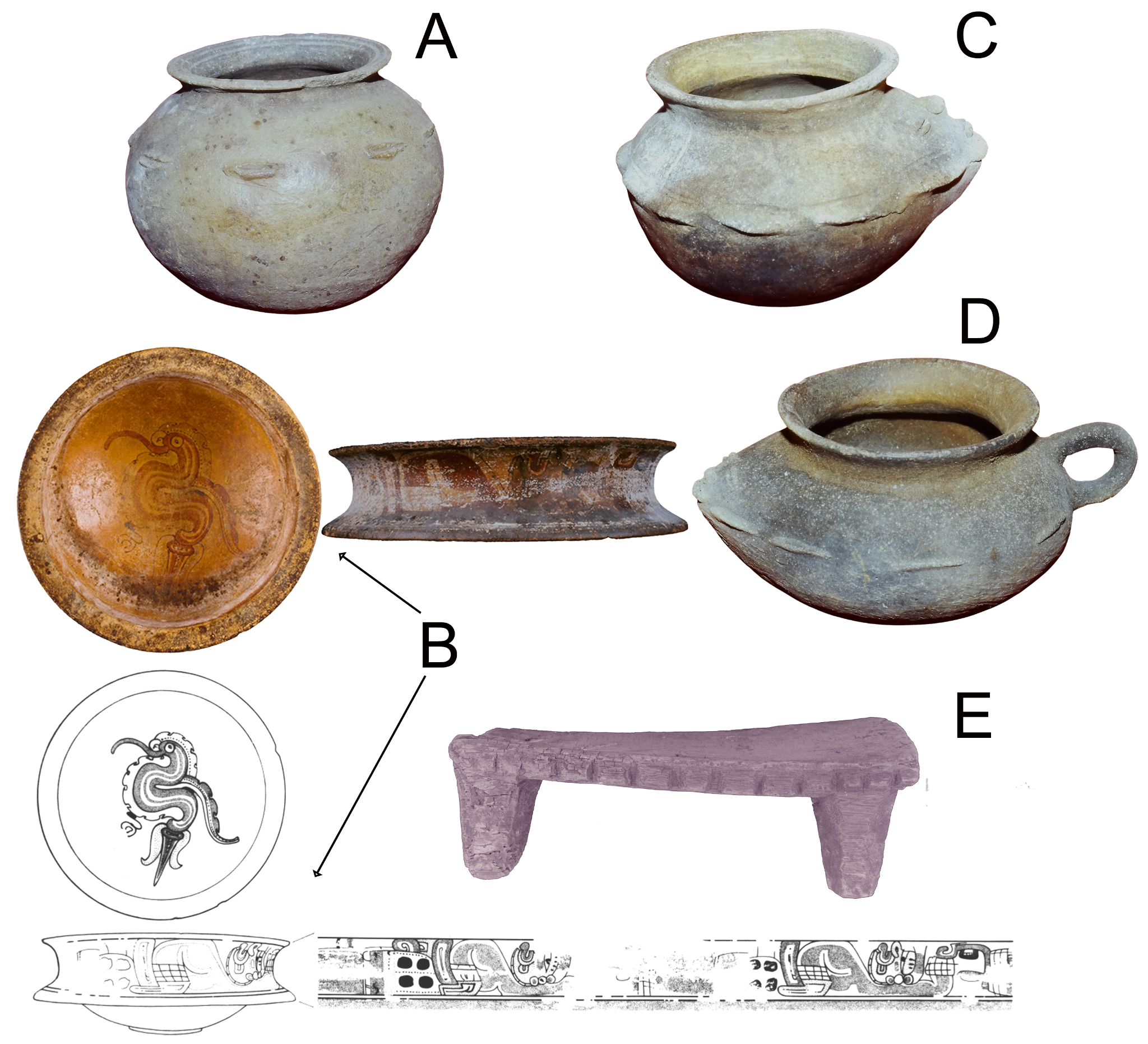

*Figure. S1.12. Artifact assemblage from Bats’ub Cave showing artifacts described in the text.*

BS contained the primary extended burial of an adult surrounded by a cache of approximately 430 commingled bones and teeth (Online Table 10). The natural clay surface on which the body was placed resulted in poor preservation such that almost all bones are broken or fragmentary, except for some complete hand and foot elements. Neither sex nor age could be estimated using osteological analyses of the commingled skeletal material, although no developing bones or teeth were found to suggest the presence of nonadults. A total of 226 permanent teeth were recovered from BS, representing an MNI of at least 15 individuals based on mandibular and maxillary left canines (Table 1). Canines are well represented compared to any other tooth class, which may reflect purposeful or preferential inclusion of specific teeth in this mortuary feature. Only three teeth exhibit dental modifications: circular drilling in the labial surfaces of two maxillary canines; mesial and distal filing on a maxillary incisor. All regions of the skeleton are at least partially represented. Pathological changes could not be observed due to taphonomic damage. Bones of the hands and feet as well as fragments of long bones with thick cortical bone (e.g. humerus, femur) were frequently recovered, but this may be due to their differential preservation (in the case of long bones) or numerous representations in the body (in the case of phalanges).

#### Saki Tzul

ST is a rockshelter formed below a sheer limestone cliff face that is located 70 m above the Ek Xux creek (Figure S1.1), a tributary of the Bladen Branch and less than 300 m away from the Classic Period Maya center of Ek Xux. ST is 145 m long and 8-to-15 m wide with ~ 1700 m^2^ of dry sediments inside the dripline, outside of which water falls. Seven years of excavations show that the site was used as a cemetery from 10,500 to 1,000 BP, a parallel chronological sequence to that of a second rockshelter, Mayahak Cab Pek (MHCP), located less than 3 km to the southwest in the neighboring valley. These two rockshelters have been critical for understanding the early population history of Central and South America^14^ as well as the origins of human diets in the region^14,15^. Classic Period burials in ST are found in the upper excavation levels. Most of the samples came from context 10 (ST.22.15.10) which is a complex commingled deposit of multiple crania and bone. The context 10 (Figure S1.13) includes at least two periods of activity based on the relationships between excavated matrices. These matrices were a complex, disturbed, and commingled jumble of redeposited bone that cut into the primary burial of two individuals interred in prone positions, both possibly interred during the same mortuary event based on their proximity. These two individuals were buried in opposing extended positions such that their legs were pointing in opposite cardinal directions, but their heads would have been within millimeters of each other, if not directly touching. One of these two individuals is I46277. Following those interments and in a later event additional burials were placed in the same location. This would have required digging into the ground surface, disturbing the earlier burials, and then the deposition of new burials. These later burials cut through at least two articulated individuals. Two others (I8564 and I19164), were excavated as isolated bone fragments in excavation fill in Classic period levels, possibly a disturbed component of context 10.

Artifacts recovered from the levels in which the burials were found include lithics, ceramics, faunal bone, and shell. Typeable ceramics date to the early Late Classic though there are a mix of ceramics that could possibly also date to the Terminal Preclassic. Lithics include obsidian blades and grinding implements including manos and metate fragments. Some special finds include a carved bone needle, bone scoop, and a jaguar tooth pendant. Due to the mixed context of the burials it is not possible to associate any of these artifacts with any particular individual.

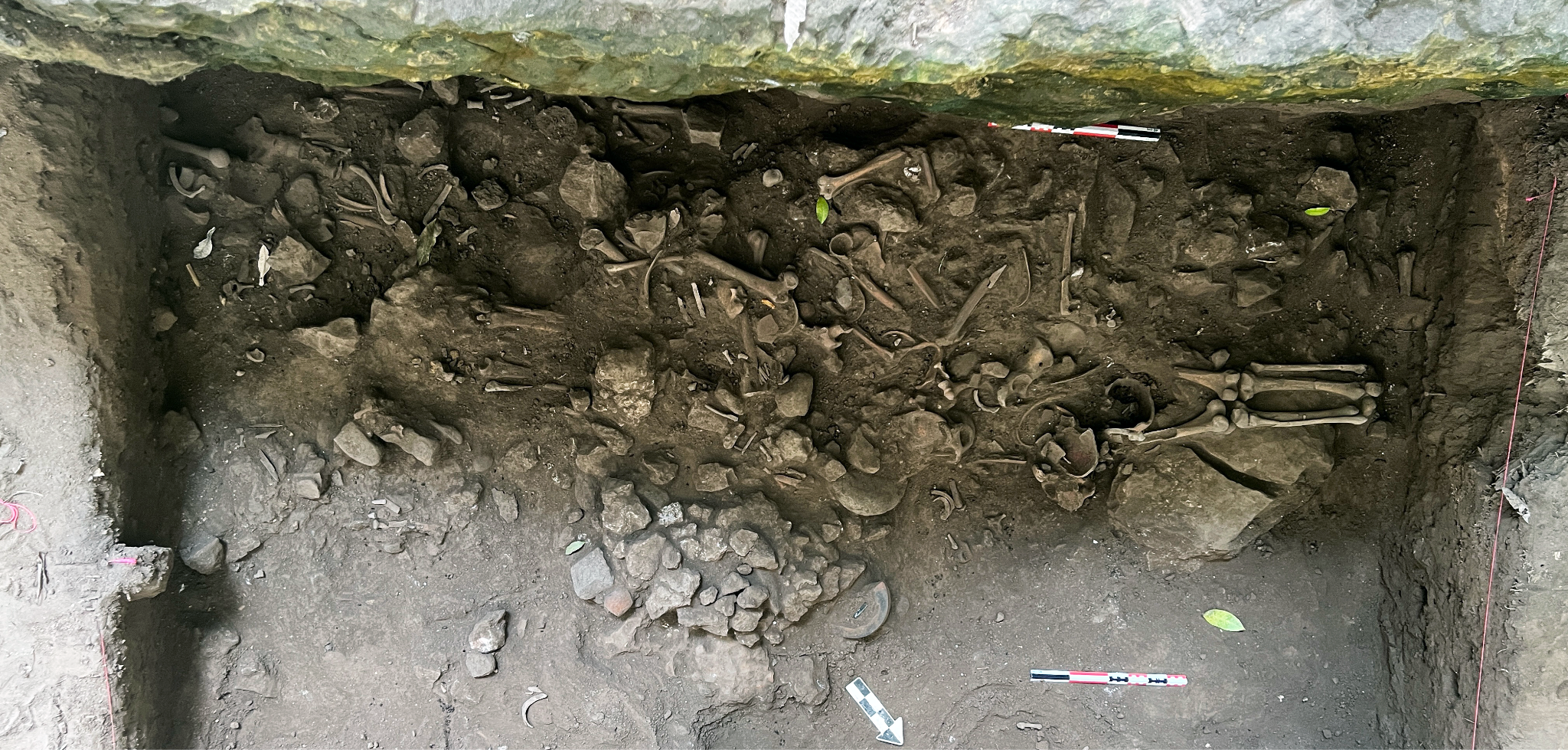

*Figure S1.13 Plan-view of the ossuary deposit in ST.22.15.10 (context 10).*

Osteological analysis of context ST.22.15.10 determined that at least six individuals are represented in the deposit: four adults, one nonadult, one neonate. MNI calculations are based on duplicated elements and the presence of bones in early stages of development. Most regions of the skeleton are equally represented, although few teeth are present. Adult MNI is based on the presence of four distinct mandibles and four frontal bones. Given the complexity of the deposit the sampling strategy for DNA analysis was broad across skeletal regions, favoring repeated elements including right temporal bones (3), right scaphoids (3), first proximal pedal phalanges (2), and mandibular teeth (3: canine, fourth premolar, first incisor).

Three of the adults were distinguishable with elements that could be matched based on similarities in size, shape, rearticulation of fragments, taphonomic changes, and pathological conditions. Some aspects of their bioprofiles could be estimated^6^, with notable pathological changes. Two are probable females: an older adult with diffuse active and remodeled periosteal reaction throughout the skeleton, remodeled porotic hyperostosis with diploic expansion is present on the cranial vault; a middle adult with remodeled porotic hyperostosis but no expansion. One individual is of indeterminate sex although not likely of advanced age based on the lack of alveolar remodeling in the maxilla and mandible in the tooth sockets. Almost all teeth are affected by postmortem tooth loss rather than antemortem, and there are no abscesses present to suggest advanced tooth decay. This individual has a small active, possibly neoplastic, lytic lesion near the right fronto-temporale landmark, with sharp margins surrounded by active microporosity. The lesion has perforated only the outer table and part of the diploe.

The nonadult individuals in ST.22.15.10 are represented by elements of a neonate and a young child. The neonatal individual is represented by bones of the pelvis and fragments of the cranium, ribs, and foot. Measurements of the pubis and ilium suggest this individual is between 36-38 fetal weeks^16^. The young child is represented by fragments of a humerus, radius, and ulna. The stage of development and maximum epiphyseal breadth of the distal humerus suggests this individual is approximately 3 years of age^17^.

#### Chabil Cab Pek

Located 1.7 km north of Batsub cave, Chabil Cab Pek (CCP - pretty stone house in Qeq’chi’ Maya) cave has a 5 m wide entrance that slopes downward into a small room (chamber A) with a narrow passage running to the south of this room and sloping upward to a masonry wall, approximately 10 m from the cave entrance (Figure S1.14). The exterior of the wall was sealed with a mud-plaster mix with traces of red pigment on the exterior. Behind the wall is a small 2 m x 1 m sealed chamber (chamber B) enclosed by a second plastered wall. This chamber has a small stone altar consisting of a tabular stone perched on two smaller stones like a table. Two ceramic vessels and skeletal material were found around the altar. One of the vessels is a basal flange bowl while the other is an unslipped jar with cacao bean applique around the shoulder. The basal flanged vessel was painted but the exterior is eroded. These two vessels are similar to the assemblage from BS cave. The human skeletal remains were found mostly between the two vessels and around the altar but there was no articulation. Just behind the second wall is chamber C which contained a large deposit of crushed limestone – possibly the material used to plaster the walls – but no evidence of burning. In a small depression next to the plaster three pieces of speleothem (natural cave formation, stalagmite or stalactite) were arranged in the shape of a cross, a likely reference to the cave-related symbolism of the quadripartite (four-part) world tree or Ceiba *pentandra* tree that links together the sky, surface world, and underworld in the ancient Maya Popol Vuh creation story^18^

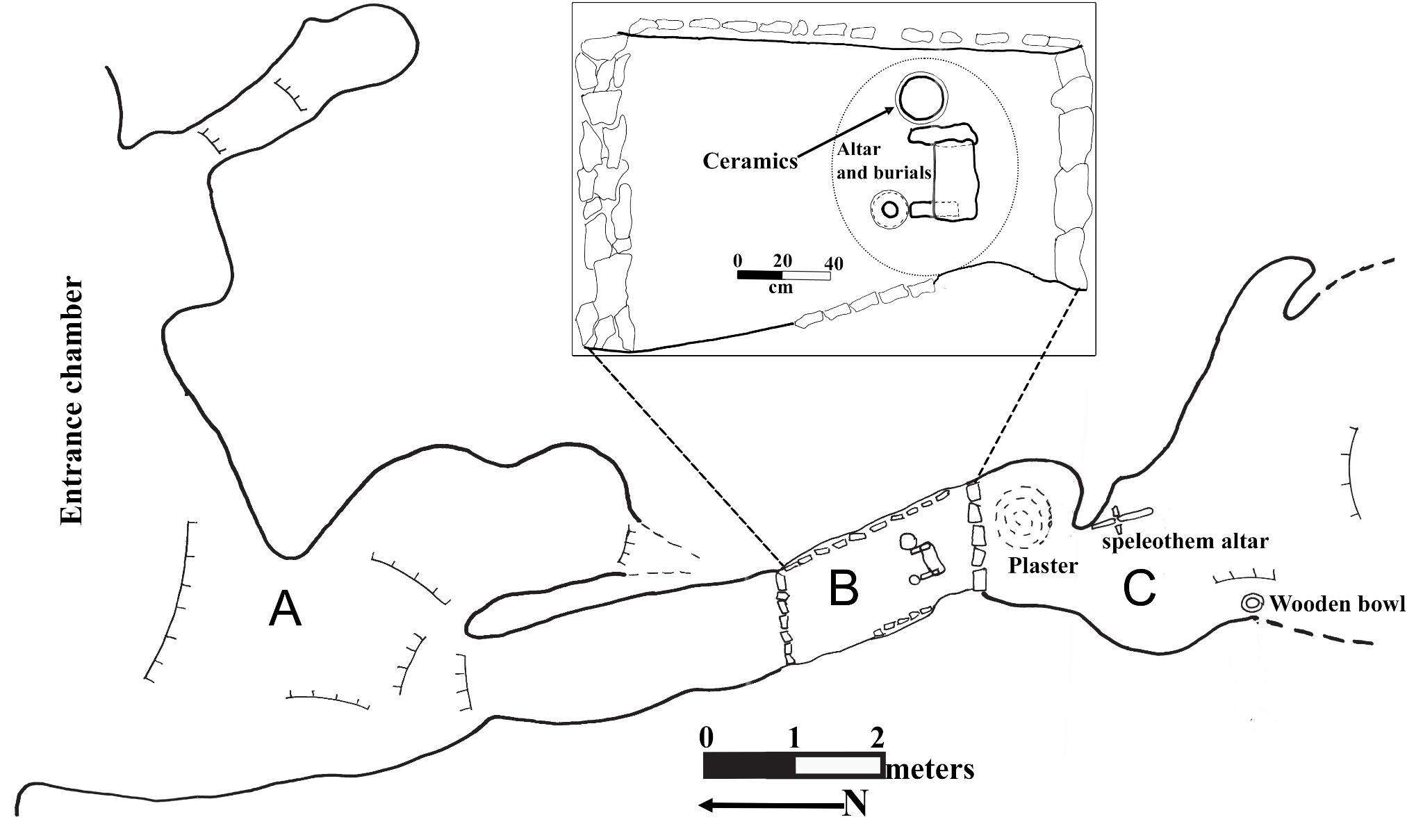

*Figure S1.14. Plan-view of Chabil Cab Pek cave with an expanded view of the burial chamber (above). The cave is divided into three chambers labeled A-C.*

CCP is represented by a small and highly fragmentary assemblage distributed across multiple excavation contexts within the cave tomb, which likely reflect arbitrary provenience designations rather than discrete mortuary events as the total area where human remains were found is very small. A total of 99 skeletal elements were recorded in the inventory, including a single permanent tooth.There was evidence of rodents gnawing on some skeletal elements. Osteological analysis indicates a minimum of two individuals based on skeletal elements, including two proximal right humeri and two right distal fibular epiphyses identified within the same context. In contrast, dental remains indicate a minimum of one individual, represented by a single lower right third premolar (LRP3), with no additional duplication observed.

Preservation across the assemblage is highly fragmentary, although certain elements are relatively well preserved, including portions of the scapular body and internal cranial structures such as the paranasal sinuses. The latter exhibit small spicules consistent with possible sinusitis. Long bones are poorly represented overall, with femora present only as fragmentary proximal and distal epiphyses. Some preserved elements appear relatively gracile, however, no estimates of stature or body proportions could be obtained due to the fragmentary nature of the assemblage.

Several elements provide limited insight into age and biological profile. A complete mandible exhibits extensive antemortem tooth loss with full resorption of the alveolar bone for all molars and the left premolar, and partial resorption affecting the canines and incisors, consistent with an older adult individual who had lost their teeth some time prior to death. A second left mandibular fragment also exhibits antemortem tooth loss. Additional indicators of advanced age include extensive lipping on a fragment of the auricular surface of the ilium and degenerative changes at the costal end of a first rib. No other skeletal features were sufficiently preserved to refine age estimates for the individuals represented. Sex estimation was limited by the absence of diagnostic os coxae features. However, the morphology of the mental eminence of the mandible and the mastoid process suggests a probable female individual, although it is unclear whether these elements derive from the same person.

### 2. Least Cost Path analysis

There are numerous approaches to calculating a Least Cost Path^19^. Since the focus of our study is not variations in paths, but understanding the general route and time it took to walk from MKB to BS, we used the basic Tobler’s hiking function. We did not account for differences in the human body (height / pace length, fitness level, etc), carrying load, or terrain type (dirt, grass, paved, etc.). In ArcGIS Pro, we ran several iterations of LCPs using different tools and took the average of their results. We ran the Distance Accumulation tool and the [deprecated] Path Distance tool, using MKB as the origin point. We included a table with the Tobler Hiking vertical factors^20^ to account for differences in hiking speed based on changes in the elevation. We also ran the Distance Accumulation tool using the “hiking” vertical factor built into ArcGIS Pro. The resulting raster files included a) an accumulated cost raster, wherein the value of each cell indicates the total time to move from the origin to that cell, and b) a backlink raster.

Using the Optimal Path as Raster tool and Cost path, we calculated Least Cost Paths for outputs from both the Distance Accumulation tool and Path Distance tool. We used BS as the destination. The resulting raster file was transformed into a polyline feature class and we calculated the total distance. If the inputs from the same Distance Accumulation output are used in the Optional Path as Raster Tool or Cost Path tool, there is no difference in the resulting Least Cost Path. Differences in LCPs stem from using the Distance Accumulation tool vs the Path Distance tool. Minor differences stem from using the built in ArcGIS Pro “hiking” vertical factor and the Tobler table vertical factor.

All in all, the LCP models allude to the likely path that was used to move from MKB to BS, which was between 26-27 km. Traveling from MKB to BS required substantial uphill climbs totaling nearly 800 m in elevation gain, particularly in the middle section of the hike just after Sachollil in the Snake Creek valley. While accumulated time calculations indicate that it would have taken minimally 6-7 hours to hike the distance from MKB to BS but this does not consider the rugged and difficult terrain in the Maya Mountains. We believe it would have taken more than one day when accounting for navigating the varied terrain of the neotropical rainforest and carrying heavy loads.

### 3. Building the lineages

Pairwise relatedness was evaluated for pairs of individuals using ancIBD^21^, READv2^22^, and reLD (an internal lab linkage disequilibrium based method). We provide three corresponding online data tables reporting the relevant pairwise outputs from each approach, including ancIBD nIBD>12 and sumIBD>12 values (Online Table 3), READv2 relationship calls (Online Table 4), and reLD likelihood-based relationship inferences (Online Table 5). We first built lineages using IBD between individuals with more than 700,000 SNPs, then performed a second pass including individuals with at least 200,000 SNPs to refine identities and, in rare cases, first-degree relationships. PMR analysis was restricted to individuals with more than 50,000 SNPs to find other lower coverage matches or first-degree relatives. Genetic matches were able to be confirmed for individuals even with fewer than 50,000 SNPs. reLD was run without a minimum SNP threshold, relying instead on its internal validation procedure, which required repeated likelihood estimates to converge on the same relationship with a 99^th^ percentile likelihood threshold.

### 4. Building the chronology

We established a high precision chronology based on AMS radiocarbon dates for 66 samples from all sites included in this study (Extended data table 2, Figure S4.1, and Table S4.2). These include direct dates on 62 individuals that bracket the sequence of human remains at MKB PT and T1, CCP, and BS between 45±49 CE and 734±40 CE. There are 4 additional dates on organic materials from Bats’ub cave. These include 2 fragments of pitch-pine torches, 1 charcoal fragment, and the wooden stool grave good. For those individuals with DNA, the I-ID is included, but all others are described as ‘Not processed’ or ‘Organic nonhuman’. All dates were calibrated using OxCal v4.6 and the IntCal 20 calibration curve^23^.

OxCal code containing uncalibrated ^14^C ages for directly dated individuals and materials:

R_Date("UCIAMS-14914 | Not processed | BS.96.B25.27_a",1975,35);

R_Date("ISGS-3527 | Organic nonhuman | BS pitch-pine torch",1920,110);

R_Date("UCIAMS-14913 | Not processed | BS.96.B25.2_c",1910,35);

R_Date("UCIAMS-14912 | Not processed | BS.95.B25.2_b",1800,35);

R_Date("UCIAMS-14915 | Not processed | BS.96.B25.11_b",1780,35);

R_Date("BETA-98152 | Organic nonhuman | BS Wooden bench",1780,60);

R_Date("UCIAMS-14922 | Not processed | BS.95.B25.2_b",1765,35);

R_Date("NOSAMS-204278 | I43789 | 5747 (PT.E96-4.LLI2-1)",1751,18);

R_Date("UCIAMS-14924 | Not processed | BS.95.B25.5_c",1745,35);

R_Date("NOSAMS-203735 | Primary BS | BS.95.B25.4_d",1733,23);

R_Date("UCIAMS-280706 | I37590 | 4406 (CBCP.96.2.A_a)",1725,15);

R_Date("ISGS-3509 | Organic nonhuman | BS pitch-pine torch",1680,70);

R_Date("NOSAMS-204290 | I43801 | 5759 (PT.96-MZ7.URI2-9)",1639,18);

R_Date("UCIAMS-228030 | I19164 | 1304 (ST.17.6.7b)",1630,25);

R_Date("ISGS-3476 | Organic nonhuman | BS carbonized wood",1610,70);

R_Date("NOSAMS-204500 | I43802 | 5760 (PT.96-MZ-8.URI1-5)",1603,18);

R_Date("NOSAMS-204268 | I43013 | 5757 (PT.96-MZ-7.L?I1-1)",1596,19);

R_Date("UCIAMS-14930 | Not processed | JI.96-MZ-5.URI1-1",1555,30);

R_Date("NOSAMS-204509 | I49214 | 5928 (PT.96-MZ-7.LLI2-4)",1554,21);

R_Date("NOSAMS-204276 | I49348 | 6226 (PT.96-MZ-7.URP3-4)",1538,18);

R_Date("UCIAMS-14931 | Not processed | JI.96-MZ-5.URI1-2",1535,30);

R_Date("NOSAMS-204281 | I42564 | 5711 (PT.96-MZ-7.URI2-6)",1527,18);

R_Date("NOSAMS-204279 | I43235 | 5905 (PT.96-MZ-7.LLI2-8)",1519,18);

R_Date("NOSAMS-204295 | I43309 | 6627 (BS.95.B25.8_an)",1501,19);

R_Date("NOSAMS-204267 | I28568 | 5697 (PT.96-MZ-7)",1500,20);

R_Date("NOSAMS-203734 | I28597 | 6524 (BS.95.B25.8_CF)",1497,17);

R_Date("PSUAMS-8383 | Not processed | BS.95.B25.8a",1495,20);

R_Date("NOSAMS-203733 | I28570 | 6620 (BS.95.B25.8_v)",1492,18);

R_Date("NOSAMS-204277 | I28570 | 5938 (T1.E96-139.LRI2-1)",1491,22);

R_Date("NOSAMS-203731 | I39918 | 4709 (MKB.99.36.1_aen)",1486,17);

R_Date("NOSAMS-204287 | I43774 | 5732 (PT.E96-88.URI1-1)",1481,17);

R_Date("PSUAMS-2147 | I8564 | UNM 860 ST.16.2.a2",1480,20);

R_Date("NOSAMS-204266 | I43727 | 5724 (PT.96-MZ-7.LLI2-5)",1474,18);

R_Date("NOSAMS-204292 | I28597 | 2632 (MKB.96.PT.P7m)",1470,17);

R_Date("NOSAMS-204280 | I28575 | 5751 (PT.96-MZ-7.LRI2-4)",1459,20);

R_Date("NOSAMS-204270 | I43319 | 5738 (PT.96-MZ-7.LLI1-5)",1455,20);

R_Date("NOSAMS-204294 | I28581 | 5753 (PT.E96-23.ULI1-1)",1453,18);

R_Date("NOSAMS-204269 | I28522 | 5745 (PT.96-MZ-7.URI2-3)",1450,20);

R_Date("NOSAMS-204264 | I43770 | 5754 (PT.96-MZ-7.LLI2-7)",1440,46);

R_Date("NOSAMS-204271 | I28517 | 5691 (PT.96-MZ-7.URI1-1)",1437,28);

R_Date("NOSAMS-204263 | I42992 | 5722 (PT.96-MZ-7.Root-1)",1435,23);

R_Date("NOSAMS-204293 | I28578 | 6572 (BS.95.B25.8_cp)",1432,20);

R_Date("NOSAMS-204282 | I49757 | 6346 (PT.E96-88.LLP4-2)",1431,17);

R_Date("NOSAMS-203732 | I39924 | 4728 (MKB.99.36.1_su)",1382,18);

R_Date("NOSAMS-204289 | I49127 | 6047 (PT.96-MZ-8.LRC-1)",1366,18);

R_Date("NOSAMS-204508 | I43797 | 5755 (PT.96-MZ-8.ULI2-1)",1361,20);

R_Date("PSUAMS 12894 | Not processed | MKB.99.36.1a",1355,20);

R_Date("UCIAMS 280699 | Not processed | MKB.99.36.1_ub",1350,15);

R_Date("NOSAMS-204288 | I48494 | 6026 (PT.96-MZ-7.LLC-2)",1345,18);

R_Date("UCIAMS 280700 | Not processed | MKB.99.36.1_dw",1345,15);

R_Date("UCIAMS 280697 | Not processed | MKB.99.36.1_vo",1345,15);

R_Date("NOSAMS-204507 | I42563 | 5736 (PT.96-MZ-8.LLI1-2)",1341,24);

R_Date("PSUAMS-8702 | Not processed | MKB.M.32.1. E95-11",1340,30);

R_Date("UCIAMS 280696 | Not processed | MKB.99.36.1_uh",1340,15);

R_Date("UCIAMS 280701 | Not processed | MKB.99.36.1_cd",1335,15);

R_Date("UCIAMS 280702 | Not processed | MKB.99.36.1_fl",1325,15);

R_Date("NOSAMS-204283 | I43012 | 5909 (PT.96-MZ-7.URI1-3)",1320,28);

R_Date("UCIAMS 280698 | Not processed | MKB.99.36.1_ui",1320,15);

R_Date("PSUAMS-8368 | Not processed | BS.95.B25.8",1315,20);

R_Date("NOSAMS-204499 | I42053 | 5737 (PT.96-MZ-8.LLI2-1)",1311,18);

R_Date("UCIAMS 280704 | Not processed | MKB.99.36.1_ay",1310,15);

R_Date("NOSAMS-204291 | I39917 | 4707 (MKB.99.36.1_qd)",1304,21);

R_Date("UCIAMS 280705 | Not processed | MKB.99.36.1_as",1300,15);

R_Date("NOSAMS-204286 | I28521 | 6501 (BS.95.B25.8_dy)",1292,18);

R_Date("UCIAMS 280703 | Not processed | MKB.99.36.1_akv",1290,15);

R_Date("NOSAMS-204502 | I46063 | 5927 (PT.95-MZ-5.LLI2-1)",1257,18);

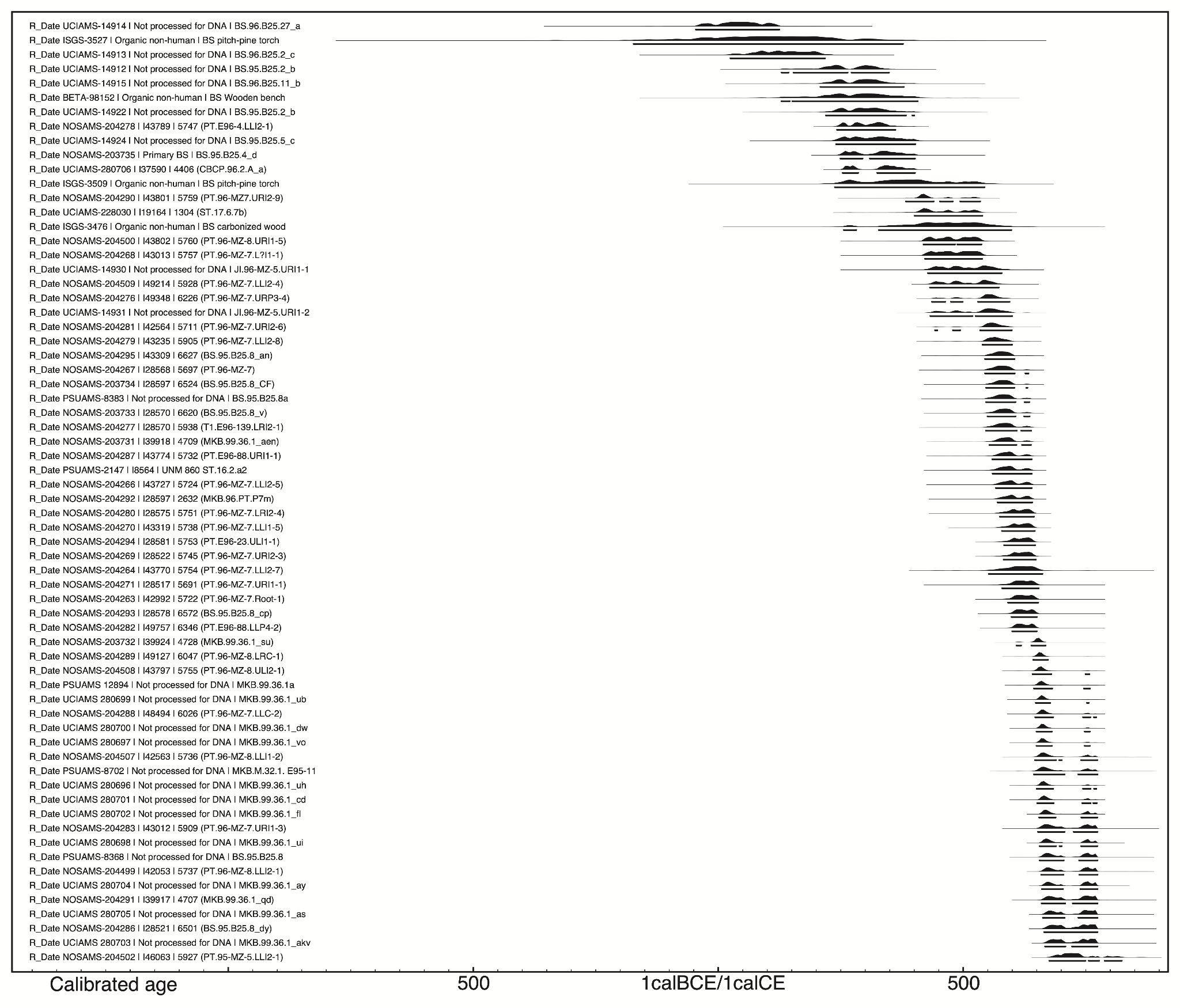

Figure S4.1. Calibrated ages of directly dated individuals and materials.

Table S4.2. Calibrated ages of directly dated individuals and materials.

| Name | Median (BCE/CE) | Unmodelled  (BCE/CE) | |
| --- | --- | --- | --- |
|  |  | From 95.4% CI | To 95.4% CI |
| R_Date UCIAMS-14914 \| Not processed for DNA \| BS.96.B25.27_a | 45 | -46 | 125 |
| R_Date ISGS-3527 \| Organic non-human \| BS pitch-pine torch | 105 | -173 | 378 |
| R_Date UCIAMS-14913 \| Not processed for DNA \| BS.96.B25.2_c | 131 | 26 | 219 |
| R_Date UCIAMS-14912 \| Not processed for DNA \| BS.95.B25.2_b | 258 | 130 | 349 |
| R_Date UCIAMS-14915 \| Not processed for DNA \| BS.96.B25.11_b | 294 | 209 | 379 |
| R_Date BETA-98152 \| Organic non-human \| BS Wooden bench | 287 | 130 | 408 |
| R_Date UCIAMS-14922 \| Not processed for DNA \| BS.95.B25.2_b | 301 | 221 | 401 |
| R_Date NOSAMS-204278 \| I43789 \| 5747 (PT.E96-4.LLI2-1) | 306 | 243 | 362 |
| R_Date UCIAMS-14924 \| Not processed for DNA \| BS.95.B25.5_c | 317 | 241 | 403 |
| R_Date NOSAMS-203735 \| Primary BS \| BS.95.B25.4_d | 331 | 250 | 402 |
| R_Date UCIAMS-280706 \| I37590 \| 4406 (CBCP.96.2.A_a) | 341 | 255 | 402 |
| R_Date ISGS-3509 \| Organic non-human \| BS pitch-pine torch | 381 | 239 | 544 |
| R_Date NOSAMS-204290 \| I43801 \| 5759 (PT.96-MZ7.URI2-9) | 425 | 384 | 535 |
| R_Date UCIAMS-228030 \| I19164 \| 1304 (ST.17.6.7b) | 462 | 402 | 540 |
| R_Date ISGS-3476 \| Organic non-human \| BS carbonized wood | 466 | 257 | 599 |
| R_Date NOSAMS-204500 \| I43802 \| 5760 (PT.96-MZ-8.URI1-5) | 478 | 420 | 538 |
| R_Date NOSAMS-204268 \| I43013 \| 5757 (PT.96-MZ-7.L?I1-1) | 481 | 423 | 539 |
| R_Date UCIAMS-14930 \| Not processed for DNA \| JI.96-MZ-5.URI1-1 | 503 | 429 | 579 |
| R_Date NOSAMS-204509 \| I49214 \| 5928 (PT.96-MZ-7.LLI2-4) | 506 | 433 | 573 |
| R_Date NOSAMS-204276 \| I49348 \| 6226 (PT.96-MZ-7.URP3-4) | 552 | 437 | 595 |
| R_Date UCIAMS-14931 \| Not processed for DNA \| JI.96-MZ-5.URI1-2 | 547 | 434 | 601 |
| R_Date NOSAMS-204281 \| I42564 \| 5711 (PT.96-MZ-7.URI2-6) | 560 | 443 | 600 |
| R_Date NOSAMS-204279 \| I43235 \| 5905 (PT.96-MZ-7.LLI2-8) | 566 | 540 | 600 |
| R_Date NOSAMS-204295 \| I43309 \| 6627 (BS.95.B25.8_an) | 579 | 545 | 605 |
| R_Date NOSAMS-204267 \| I28568 \| 5697 (PT.96-MZ-7) | 579 | 545 | 634 |
| R_Date NOSAMS-203734 \| I28597 \| 6524 (BS.95.B25.8_CF) | 581 | 547 | 632 |
| R_Date PSUAMS-8383 \| Not processed for DNA \| BS.95.B25.8a | 582 | 547 | 636 |
| R_Date NOSAMS-203733 \| I28570 \| 6620 (BS.95.B25.8_v) | 584 | 550 | 636 |
| R_Date NOSAMS-204277 \| I28570 \| 5938 (T1.E96-139.LRI2-1) | 585 | 547 | 639 |
| R_Date NOSAMS-203731 \| I39918 \| 4709 (MKB.99.36.1_aen) | 588 | 554 | 639 |
| R_Date NOSAMS-204287 \| I43774 \| 5732 (PT.E96-88.URI1-1) | 592 | 560 | 640 |
| R_Date PSUAMS-2147 \| I8564 \| UNM 860 ST.16.2.a2 | 593 | 561 | 641 |
| R_Date NOSAMS-204266 \| I43727 \| 5724 (PT.96-MZ-7.LLI2-5) | 598 | 567 | 641 |
| R_Date NOSAMS-204292 \| I28597 \| 2632 (MKB.96.PT.P7m) | 602 | 571 | 642 |
| R_Date NOSAMS-204280 \| I28575 \| 5751 (PT.96-MZ-7.LRI2-4) | 614 | 575 | 645 |
| R_Date NOSAMS-204270 \| I43319 \| 5738 (PT.96-MZ-7.LLI1-5) | 618 | 580 | 647 |
| R_Date NOSAMS-204294 \| I28581 \| 5753 (PT.E96-23.ULI1-1) | 619 | 584 | 647 |
| R_Date NOSAMS-204269 \| I28522 \| 5745 (PT.96-MZ-7.URI2-3) | 620 | 583 | 649 |
| R_Date NOSAMS-204264 \| I43770 \| 5754 (PT.96-MZ-7.LLI2-7) | 616 | 553 | 662 |
| R_Date NOSAMS-204271 \| I28517 \| 5691 (PT.96-MZ-7.URI1-1) | 622 | 580 | 655 |
| R_Date NOSAMS-204263 \| I42992 \| 5722 (PT.96-MZ-7.Root-1) | 623 | 592 | 654 |
| R_Date NOSAMS-204293 \| I28578 \| 6572 (BS.95.B25.8_cp) | 624 | 599 | 651 |
| R_Date NOSAMS-204282 \| I49757 \| 6346 (PT.E96-88.LLP4-2) | 624 | 601 | 651 |
| R_Date NOSAMS-203732 \| I39924 \| 4728 (MKB.99.36.1_su) | 653 | 610 | 669 |
| R_Date NOSAMS-204289 \| I49127 \| 6047 (PT.96-MZ-8.LRC-1) | 658 | 644 | 674 |
| R_Date NOSAMS-204508 \| I43797 \| 5755 (PT.96-MZ-8.ULI2-1) | 659 | 642 | 758 |
| R_Date PSUAMS 12894 \| Not processed for DNA \| MKB.99.36.1a | 661 | 645 | 759 |
| R_Date UCIAMS 280699 \| Not processed for DNA \| MKB.99.36.1_ub | 662 | 649 | 757 |
| R_Date NOSAMS-204288 \| I48494 \| 6026 (PT.96-MZ-7.LLC-2) | 665 | 648 | 772 |
| R_Date UCIAMS 280700 \| Not processed for DNA \| MKB.99.36.1_dw | 664 | 650 | 759 |
| R_Date UCIAMS 280697 \| Not processed for DNA \| MKB.99.36.1_vo | 664 | 650 | 759 |
| R_Date NOSAMS-204507 \| I42563 \| 5736 (PT.96-MZ-8.LLI1-2) | 671 | 647 | 774 |
| R_Date PSUAMS-8702 \| Not processed for DNA \| MKB.M.32.1. E95-11 | 677 | 645 | 775 |
| R_Date UCIAMS 280696 \| Not processed for DNA \| MKB.99.36.1_uh | 666 | 651 | 772 |
| R_Date UCIAMS 280701 \| Not processed for DNA \| MKB.99.36.1_cd | 669 | 652 | 773 |
| R_Date UCIAMS 280702 \| Not processed for DNA \| MKB.99.36.1_fl | 677 | 656 | 774 |
| R_Date NOSAMS-204283 \| I43012 \| 5909 (PT.96-MZ-7.URI1-3) | 701 | 653 | 775 |
| R_Date UCIAMS 280698 \| Not processed for DNA \| MKB.99.36.1_ui | 685 | 657 | 774 |
| R_Date PSUAMS-8368 \| Not processed for DNA \| BS.95.B25.8 | 703 | 657 | 775 |
| R_Date NOSAMS-204499 \| I42053 \| 5737 (PT.96-MZ-8.LLI2-1) | 727 | 660 | 775 |
| R_Date UCIAMS 280704 \| Not processed for DNA \| MKB.99.36.1_ay | 740 | 661 | 775 |
| R_Date NOSAMS-204291 \| I39917 \| 4707 (MKB.99.36.1_qd) | 728 | 661 | 775 |
| R_Date UCIAMS 280705 \| Not processed for DNA \| MKB.99.36.1_as | 740 | 663 | 775 |
| R_Date NOSAMS-204286 \| I28521 \| 6501 (BS.95.B25.8_dy) | 729 | 666 | 775 |
| R_Date UCIAMS 280703 \| Not processed for DNA \| MKB.99.36.1_akv | 730 | 668 | 775 |
| R_Date NOSAMS-204502 \| I46063 \| 5927 (PT.95-MZ-5.LLI2-1) | 724 | 676 | 824 |

We used OxCal^24^ to model and constrain lineage age ranges and to calculate the ages for dated and undated lineage members. This model was constrained by direct dates and pairwise relationships of lineage members. Most modeled ages were constrained to 95.4% credible intervals spanning only a few decades, although some boundaries and broader spans remain less precise.

```
 Sequence()
 {
  Boundary("Start L1b");
  Phase ("L1b: Generation 1")
  {
   R_Date("I43801",1639,18);
   Date("I49519");
  };
  Interval(N(28,10));
  Phase("L1b: Generation 2 and below split")
  {
   Sequence()
   {
    R_Date("I49348",1538,18);
    Interval(N(28,10));
    Date("L1b: Generation 3 main");
    Interval(N(28,10));
    Date("I49967");
    Interval(N(28,10));
    Boundary("Start L1b Generation 5 main");
    Phase("L1b: Generation 5 main")
    {
     Sequence()
     {
      R_Combine("I28570_R_Combine")
      {
       R_Date("NOSAMS-203733 | I28570 | 6620 (BS.95.B25.8_v)", 1492, 18);
       R_Date("NOSAMS-204277 | I28570 | 5938 (T1.E96-139.LRI2-1)", 1491, 22);
      };
      Interval(N(28,10));
      Date("I28520");
     };
     R_Date("NOSAMS-204294 | I28581 | 5753 (PT.E96-23.ULI1-1)", 1453, 18);
    };
    Boundary("End L1b Generation 5 main");
   };
   Sequence()
   {
    Date("L1b: Generation 2 offshoot");
    Interval(N(28,10));
    Date("L1b: Generation 3 offshoot");
    Interval(N(28,10));
    Boundary("limit cousins time span start");
    Phase("ST cousins constraint")
    {
     R_Combine("I28597_R_Combine")
     {
      R_Date("NOSAMS-203734 | I28597 | 6524 (BS.95.B25.8_CF)",1497,17);
      R_Date("NOSAMS-204292 | I28597 | 2632 (MKB.96.PT.P7m)",1470,17);
     };
     R_Date("UCIAMS-228030 | I19164 | 1304 (ST.17.6.7b)",1630,25);
     Span(U(0,40));
    };
    Boundary("limit cousins time span end");
   };
  };
  Span("Span of L1b dates");
  Boundary("End L1b");
 };
 Sequence()
 {
  Boundary("Start L2");
  R_Date("NOSAMS-204281 | I42564 | 5711 (PT.96-MZ-7.URI2-6)",1527,18);
  Interval(N(28,10));
  Date("L2: Generation 2");
  Interval(N(28,10));
  Date("L2: Generation 3");
  Interval(N(28,10));
  Phase("L2: Generation 4 and below")
  {
   Sequence()
   {
    Boundary("Start L2a Generation 4 to end");
    Boundary("Start L2a Generation 4");
    Phase("L2a: Generation 4a")
    {
     R_Date("NOSAMS-204279 | I43235 | 5905 (PT.96-MZ-7.LLI2-8)",1519,18);
     R_Date("NOSAMS-204509 | I49214 | 5928 (PT.96-MZ-7.LLI2-4)",1554,21);
     Interval(N(28,10));
    };
    Boundary("End L2a Generation 4");
    Boundary("Start L2a Generation 5");
    Phase("L2a: Generation 5")
    {
     R_Date("NOSAMS-204287 | I43774 | 5732 (PT.E96-88.URI1-1)",1481,17);
     Date("I43244");
     Interval(N(28,10));
    };
    Boundary("End L2a Generation 5");
    Boundary("Start L2a Generations 6-7");
    Phase("L2a: Generation 6 and below")
    {
     R_Date("NOSAMS-204264 | I43770 | 5754 (PT.96-MZ-7.LLI2-7)",1440,46);
     R_Date("NOSAMS-204288 | I48494 | 6026 (PT.96-MZ-7.LLC-2)",1345,18);
     Sequence()
     {
      R_Date("NOSAMS-204508 | I43797 | 5755 (PT.96-MZ-8.ULI2-1)",1361,20);
      Interval(N(28,10));
      R_Date("NOSAMS-204507 | I42563 | 5736 (PT.96-MZ-8.LLI1-2)",1341,24);
     };
    };
    Boundary("End L2a Generations 6-7");
    Boundary("End L2a Generation 4 to end");
   };
   Sequence()
   {
    Boundary("Start L2b Generation 4 to end");
    Date("I49489");
    Interval(N(28,10));
    Boundary("Start L2b Generation 5 and below");
    Phase("L2b: Generation 5 and below split")
    {
     Sequence()
     {
      Date("L2b: Generation 5 offshoot father");
      Interval(N(28,10));
      R_Date("NOSAMS-204269 | I28522 | 5745 (PT.96-MZ-7.URI2-3)",1450,20);
     };
     Sequence()
     {
      Date("L2b: Generation 5 main father");
      Interval(N(28,10));
      R_Date("NOSAMS-204270 | I43319 | 5738 (PT.96-MZ-7.LLI1-5)",1455,20);
      Interval(N(28,10));
      Boundary("Start L2b Generation 7-8");
      Phase("L2b: Generation 7 and below main")
      {
       R_Date("NOSAMS-204271 | I28517 | 5691 (PT.96-MZ-7.URI1-1)",1437,28);
       R_Date("NOSAMS-204295 | I43309 | 6627 (BS.95.B25.8_an)",1501,19);
       Sequence()
       {
        R_Date("NOSAMS-204293 | I28578 | 6572 (BS.95.B25.8_cp)",1432,20);
        Interval(N(28,10));
        R_Date("NOSAMS-204282 | I49757 | 6346 (PT.E96-88.LLP4-2)",1431,17);
       };
      };
      Boundary("End L2b Generation 7-8");
     };
    };
    Boundary("End L2b Generation 5 and below");
    Boundary("End L2b Generation 4 to end");
   };
  };
  Span("Span of L2 dates");
  Boundary("End L2");
 };
 Sequence()
 {
  Boundary("Start L3");
  R_Date("NOSAMS-203731 | I39918 | 4709 (MKB.99.36.1_aen)",1486,17);
  Interval(N(28,10));
  Date("L3: Generation 2");
  Interval(N(28,10));
  Date("I39921");
  Interval(N(28,10));
  R_Date("NOSAMS-203732 | I39924 | 4728 (MKB.99.36.1_su)",1382,18);
  Span("Span of L3 dates");
  Boundary("End L3");
 };
```

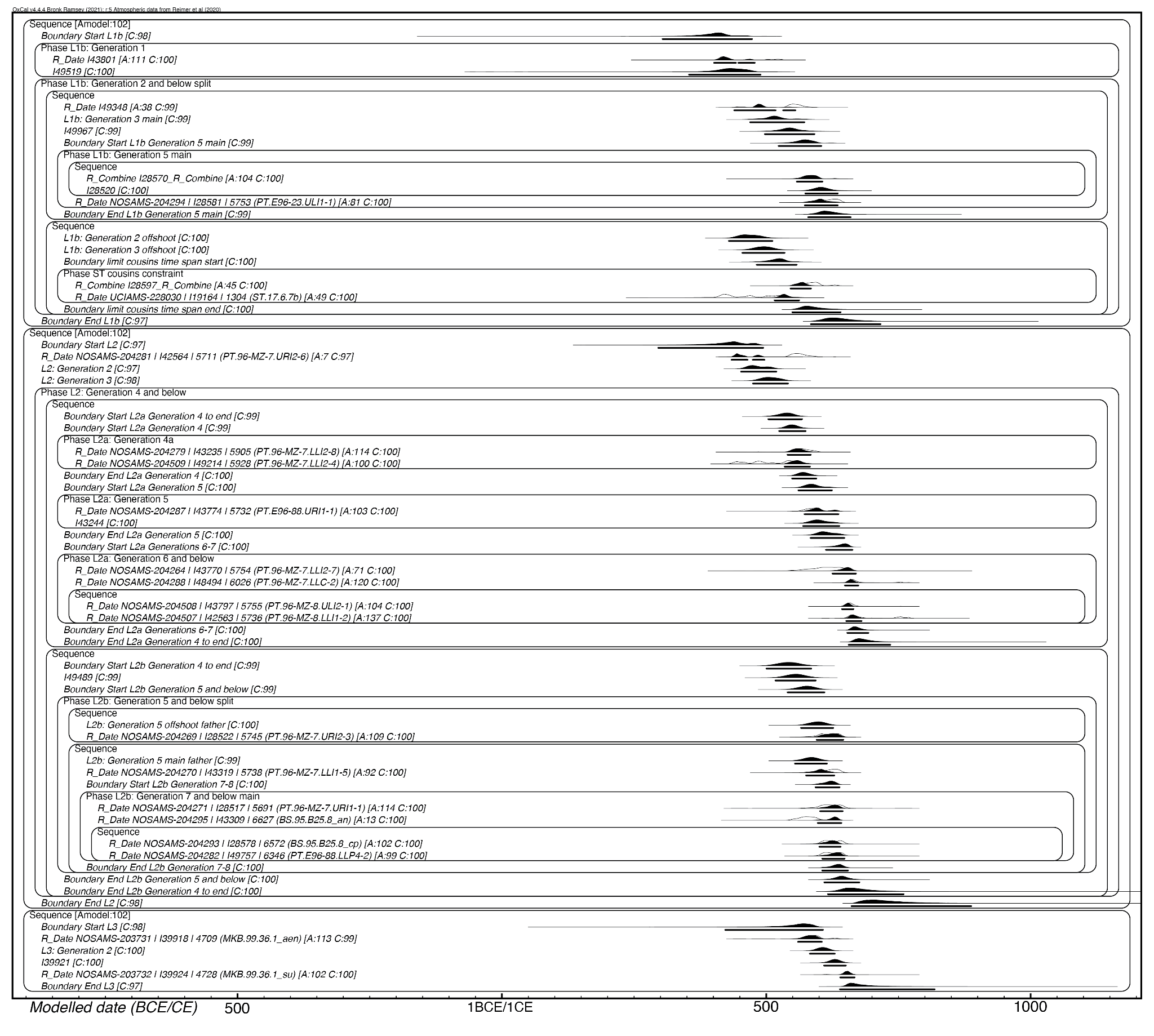

Figure S4.3. OxCal output for modeling and constraining pedigree age ranges.

Table S4.4. OxCal output for modeling and constraining pedigree age ranges.

| ***Name*** | **Unmodelled (BCE/CE)** | | **Modelled (BCE/CE)** | | **Indices** | |
| --- | --- | --- | --- | --- | --- | --- |
|  | **from_95_4** | **to_95_4** | **from_95_4** | **to_95_4** | **A** | **C** |
| *Sequence* |  |  |  |  |  |  |
| *Boundary Start L1b* |  |  | 305 | 474 |  | 98.2 |
| *Phase L1b: Generation 1* |  |  |  |  |  |  |
| *R_Date I43801* | 384 | 535 | 402 | 479 | 111.4 | 99.8 |
| *I49519* |  |  | 355 | 490 |  | 99.6 |
| *Interval* | 8 | 48 | 5 | 44 | 97.3 | 99.8 |
| *N(28,10)* | 8 | 48 |  |  |  |  |
| *Phase L1b: Generation 2 and below split* |  |  |  |  |  |  |
| *Sequence* |  |  |  |  |  |  |
| *R_Date I49348* | 437 | 595 | 440 | 556 | 37.7 | 99.1 |
| *Interval* | 8 | 48 | 7 | 47 | 99.5 | 99.6 |
| *N(28,10)* | 8 | 48 |  |  |  |  |
| *L1b: Generation 3 main* |  |  | 470 | 573 |  | 99.1 |
| *Interval* | 8 | 48 | 7 | 47 | 99.4 | 99.6 |
| *N(28,10)* | 8 | 48 |  |  |  |  |
| *I49967* |  |  | 498 | 592 |  | 99.3 |
| *Interval* | 8 | 48 | 7 | 47 | 99.3 | 99.6 |
| *N(28,10)* | 8 | 48 |  |  |  |  |
| *Boundary Start L1b Generation 5 main* |  |  | 524 | 605 |  | 99.4 |
| *Phase L1b: Generation 5 main* |  |  |  |  |  |  |
| *Sequence* |  |  |  |  |  |  |
| *R_Combine I28570_R_Combine* | 553 | 632 | 559 | 607 | 104 | 99.8 |
| *Interval* | 8 | 48 | 0 | 38 | 82.9 | 99 |
| *N(28,10)* | 8 | 48 |  |  |  |  |
| *I28520* |  |  | 575 | 636 |  | 99.7 |
| *R_Date NOSAMS-204294 \| I28581 \| 5753 (PT.E96-23.ULI1-1)* | 584 | 647 | 574 | 636 | 81.2 | 99.8 |
| *Boundary End L1b Generation 5 main* |  |  | 580 | 660 |  | 99.3 |
| *Sequence* |  |  |  |  |  |  |
| *L1b: Generation 2 offshoot* |  |  | 430 | 512 |  | 99.7 |
| *Interval* | 8 | 48 | 6 | 45 | 99.1 | 99.7 |
| *N(28,10)* | 8 | 48 |  |  |  |  |
| *L1b: Generation 3 offshoot* |  |  | 456 | 535 |  | 99.8 |
| *Interval* | 8 | 48 | 6 | 45 | 99.3 | 99.8 |
| *N(28,10)* | 8 | 48 |  |  |  |  |
| *Boundary limit cousins time span start* |  |  | 483 | 558 |  | 99.8 |
| *Phase ST cousins constraint* |  |  |  |  |  |  |
| *R_Combine I28597_R_Combine* | 563 | 637 | 547 | 585 | 44.9 | 99.9 |
| *R_Date UCIAMS-228030 \| I19164 \| 1304 (ST.17.6.7b)* | 402 | 540 | 517 | 563 | 49.3 | 99.7 |
| *Span* | 0 | 40 | 0 | 40 | 100 | 99.4 |
| *U(0,40)* | 7.98E-17 | 40 |  |  |  |  |
| *Boundary limit cousins time span end* |  |  | 550 | 641 |  | 99.9 |
| *Span Span of L1b dates* |  |  | 133 | 376 |  | 97.6 |
| *Boundary End L1b* |  |  | 585 | 717 |  | 97.4 |
| *Sequence* |  |  |  |  |  |  |
| *Boundary Start L2* |  |  | 297 | 495 |  | 97 |
| *R_Date NOSAMS-204281 \| I42564 \| 5711 (PT.96-MZ-7.URI2-6)* | 443 | 600 | 435 | 497 | 7.3 | 96.8 |
| *Interval* | 8 | 48 | 4 | 42 | 96 | 99.6 |
| *N(28,10)* | 8 | 48 |  |  |  |  |
| *L2: Generation 2* |  |  | 453 | 520 |  | 96.9 |
| *Interval* | 8 | 48 | 4 | 42 | 95.9 | 99.5 |
| *N(28,10)* | 8 | 48 |  |  |  |  |
| *L2: Generation 3* |  |  | 476 | 542 |  | 97.8 |
| *Interval* | 8 | 48 | 4 | 42 | 95.9 | 99.6 |
| *N(28,10)* | 8 | 48 |  |  |  |  |
| *Phase L2: Generation 4 and below* |  |  |  |  |  |  |
| *Sequence* |  |  |  |  |  |  |
| *Boundary Start L2a Generation 4 to end* |  |  | 505 | 568 |  | 98.7 |
| *Boundary Start L2a Generation 4* |  |  | 525 | 575 |  | 99.3 |
| *Phase L2a: Generation 4a* |  |  |  |  |  |  |
| *R_Date NOSAMS-204279 \| I43235 \| 5905 (PT.96-MZ-7.LLI2-8)* | 540 | 600 | 541 | 585 | 113.9 | 99.6 |
| *R_Date NOSAMS-204509 \| I49214 \| 5928 (PT.96-MZ-7.LLI2-4)* | 433 | 573 | 536 | 583 | 100.2 | 99.5 |
| *Interval* | 8 | 48 | 3 | 40 | 93.6 | 99.7 |
| *N(28,10)* | 8 | 48 |  |  |  |  |
| *Boundary End L2a Generation 4* |  |  | 550 | 595 |  | 99.6 |
| *Boundary Start L2a Generation 5* |  |  | 562 | 625 |  | 99.8 |
| *Phase L2a: Generation 5* |  |  |  |  |  |  |
| *R_Date NOSAMS-204287 \| I43774 \| 5732 (PT.E96-88.URI1-1)* | *560* | *640* | *573* | *637* | *103.4* | *99.8* |
| *I43244* |  |  | 571 | 639 |  | 99.8 |
| *Interval* | 8 | 48 | 6 | 44 | 99.6 | 99.9 |
| *N(28,10)* | 8 | 48 |  |  |  |  |
| *Boundary End L2a Generation 5* |  |  | 585 | 648 |  | 99.7 |
| *Boundary Start L2a Generations 6-7* |  |  | 614 | 664 |  | 99.6 |
| *Phase L2a: Generation 6 and below* |  |  |  |  |  |  |
| *R_Date NOSAMS-204264 \| I43770 \| 5754 (PT.96-MZ-7.LLI2-7)* | 553 | 662 | 626 | 670 | 71.2 | 99.7 |
| *R_Date NOSAMS-204288 \| I48494 \| 6026 (PT.96-MZ-7.LLC-2)* | 648 | 772 | 650 | 675 | 119.5 | 99.9 |
| *Sequence* |  |  |  |  |  |  |
| *R_Date NOSAMS-204508 \| I43797 \| 5755 (PT.96-MZ-8.ULI2-1)* | 642 | 758 | 644 | 665 | 104 | 100 |
| *Interval* | 8 | 48 | 0 | 24 | 42.4 | 99.2 |
| *N(28,10)* | 8 | 48 |  |  |  |  |
| *R_Date NOSAMS-204507 \| I42563 \| 5736 (PT.96-MZ-8.LLI1-2)* | 647 | 774 | 652 | 681 | 137.1 | 99.6 |
| *Boundary End L2a Generations 6-7* |  |  | 654 | 694 |  | 99.5 |
| *Boundary End L2a Generation 4 to end* |  |  | 657 | 735 |  | 99.5 |
| *Sequence* |  |  |  |  |  |  |
| *Boundary Start L2b Generation 4 to end* |  |  | 501 | 586 |  | 98.9 |
| *I49489* |  |  | 518 | 594 |  | 99.2 |
| *Interval* | 8 | 48 | 0 | 38 | 85.3 | 99.7 |
| *N(28,10)* | 8 | 48 |  |  |  |  |
| *Boundary Start L2b Generation 5 and below* |  |  | 541 | 611 |  | 99.2 |
| *Phase L2b: Generation 5 and below split* |  |  |  |  |  |  |
| *Sequence* |  |  |  |  |  |  |
| *L2b: Generation 5 offshoot father* |  |  | 567 | 627 |  | 99.8 |
| *Interval* | 8 | 48 | 6 | 44 | 99.5 | 99.8 |
| *N(28,10)* | 8 | 48 |  |  |  |  |
| *R_Date NOSAMS-204269 \| I28522 \| 5745 (PT.96-MZ-7.URI2-3)* | 583 | 649 | 596 | 646 | 109 | 99.9 |
| *Sequence* |  |  |  |  |  |  |
| *L2b: Generation 5 main father* |  |  | 555 | 615 |  | 99.4 |
| *Interval* | 8 | 48 | 0 | 34 | 77.2 | 99.8 |
| *N(28,10)* | 8 | 48 |  |  |  |  |
| *R_Date NOSAMS-204270 \| I43319 \| 5738 (PT.96-MZ-7.LLI1-5)* | 580 | 647 | 576 | 629 | 92.3 | 99.6 |
| *Interval* | 8 | 48 | 0 | 30 | 67.1 | 99.8 |
| *N(28,10)* | 8 | 48 |  |  |  |  |
| *Boundary Start L2b Generation 7-8* |  |  | 594 | 639 |  | 99.7 |
| *Phase L2b: Generation 7 and below main* |  |  |  |  |  |  |
| *R_Date NOSAMS-204271 \| I28517 \| 5691 (PT.96-MZ-7.URI1-1)* | 580 | 655 | 603 | 645 | 114.4 | 99.9 |
| *R_Date NOSAMS-204295 \| I43309 \| 6627 (BS.95.B25.8_an)* | 545 | 605 | 598 | 643 | 13.2 | 99.9 |
| *Sequence* |  |  |  |  |  |  |
| *R_Date NOSAMS-204293 \| I28578 \| 6572 (BS.95.B25.8_cp)* | 599 | 651 | 601 | 641 | 101.9 | 99.8 |
| *Interval* | 8 | 48 | 0 | 23 | 29.1 | 99.7 |
| *N(28,10)* | 8 | 48 |  |  |  |  |
| *R_Date NOSAMS-204282 \| I49757 \| 6346 (PT.E96-88.LLP4-2)* | 601 | 651 | 607 | 649 | 98.7 | 99.9 |
| *Boundary End L2b Generation 7-8* |  |  | 607 | 655 |  | 99.8 |
| *Boundary End L2b Generation 5 and below* |  |  | 611 | 677 |  | 99.8 |
| *Boundary End L2b Generation 4 to end* |  |  | 618 | 760 |  | 99.6 |
| *Span Span of L2 dates* |  |  | 186 | 535 |  | 97.1 |
| *Boundary End L2* |  |  | 662 | 888 |  | 98.3 |
| *Sequence* |  |  |  |  |  |  |
| *Boundary Start L3* |  |  | 423 | 608 |  | 98.1 |
| *R_Date NOSAMS-203731 \| I39918 \| 4709 (MKB.99.36.1_aen)* | 554 | 639 | 561 | 606 | 112.6 | 99.3 |
| *Interval* | 8 | 48 | 5 | 41 | 98.3 | 99.6 |
| *N(28,10)* | 8 | 48 |  |  |  |  |
| *L3: Generation 2* |  |  | 583 | 630 |  | 99.8 |
| *Interval* | 8 | 48 | 6 | 41 | 98.5 | 99.5 |
| *N(28,10)* | 8 | 48 |  |  |  |  |
| *I39921* |  |  | 610 | 651 |  | 99.8 |
| *Interval* | 8 | 48 | 5 | 41 | 98.4 | 99.5 |
| *N(28,10)* | 8 | 48 |  |  |  |  |
| *R_Date NOSAMS-203732 \| I39924 \| 4728 (MKB.99.36.1_su)* | 610 | 669 | 641 | 668 | 102.2 | 99.7 |
| *Span Span of L3 dates* |  |  | 46 | 343 |  | 97.4 |
| *Boundary End L3* |  |  | 640 | 819 |  | 97.3 |

### 5. Significance of replicate interments by lineage or site

We tested whether the proportion of cross-site replicates differs across reconstructed lineages and mortuary contexts (Table S5.1). For pairwise comparisons between two lineage families or contexts, we used two-sided Fisher exact tests. For multi-group comparisons, we used χ^2^ tests of independence.

A global χ^2^ test across L1-b, L2-a, L2-b, and L3 supports heterogeneity in the proportion of replicates across lineages (χ^2^ *p* = 0.05, Table S5.2). However, most pairwise lineage comparisons have limited power because sample sizes are small, especially for L3 (Figure S5.3). The pairwise results are most informative for L2-b, which has the most extreme proportion of cross-site replicates. This is apparent in that the lineage-level χ^2^ signal is driven largely by L2-b, and weakens when L2-a and L2-b are merged into L2 with the additional root ancestor, yielding no evidence of heterogeneity across L1-b, L2, and L3 (χ^2^ *p* = 0.26, Table S5.2).

The proportion of cross-site replicates differs significantly between the elite lineage L2-b and the non-elite lineage L3 (Fisher *p* = 0.03, Table S5.2). The proportion of cross-site replicates is also higher in L2-b than L2-a (two-sided Fisher *p* = 0.06, Table S5.2; one-sided Fisher *p* = 0.04, not shown). The power analysis in Figures S5.3–S5.4 shows that several lineage contrasts, and notably L2-b vs L2-a, are underpowered at current n, but that the null and alternative *p*-value distributions separate clearly, with the alternative hypothesis *p*-values showing a distinctively non-uniform distribution, skewed toward zero. Taken together, this is consistent with treating L2-a and L2-b as distinct sub-lineages for analysis.

In contrast to lineage-level tests, mortuary context-level comparisons have larger sample sizes and show strong heterogeneity in proportions of cross-site replicates. A global χ^2^ test across PT, T1, BS, CCP, ST, W5, and W1 is significant (χ^2^ *p* = 2.6×10⁻⁷, Table S5.2), indicating that cross-site replication is concentrated in specific contexts rather than being broadly distributed across the mortuary landscape. Elite tomb contexts contain cross-site replicates, whereas the sampled non-elite tomb contexts do not. Among 89 individuals from PT and T1, 25 are cross-site replicates, while among 18 individuals from W5 and W1, none are cross-site replicates (two-sided Fisher *p* = 0.006; χ^2^ *p* = 0.01, Table S5.2). The same pattern holds for PT versus W5 alone (25 of 88 versus 0 of 17, two-sided Fisher *p* = 0.01). BS is also enriched for cross-site replicates compared to ST (24 of 30 versus 0 of 4, two-sided Fisher *p* = 0.005).

Because contexts differ in how many skeletal elements yielded genome-wide data, we additionally tested whether the observed BS–elite-tomb overlap could be explained by sampling depth alone. We downsampled the elite element pool (all genotyped elements from PT) to match the number of genotyped elements from the pooled non-elite tomb contexts (W5+W1). For each downsample, we collapsed sampled elements to unique individuals and recomputed overlap with BS (and separately with BS+CCP). Under this downsampled null, the expected overlap is much smaller than observed, and in our simulations the number of BS individuals also represented in PT never approached the observed value. Thus, unequal numbers of genotyped elements across contexts cannot explain the strong BS–elite-tomb overlap.

| Lineage family or mortuary context (tomb, cave, rockshelter) | No. cross-site replicates | Total no. individuals | Proportion of cross-site replicates |
| --- | --- | --- | --- |
| L1-b | 3 | 7 | 0.43 |
| L2-a | 4 | 12 | 0.33 |
| L2-b | 6 | 7 | 0.86 |
| L2 | 10 | 20 | 0.53 |
| Elite lineages (L1 and L2) | 13 | 27 | 0.5 |
| L3 | 0 | 3 | 0 |
| PT | 25 | 88 | 0.28 |
| T1 | 2 | 3 | 0.67 |
| BS | 24 | 30 | 0.8 |
| CCP | 1 | 2 | 0.5 |
| ST | 0 | 4 | 0 |
| W5 | 0 | 17 | 0 |
| W1 | 0 | 1 | 0 |

*Table S5.1. Numbers of individuals classified as cross-site replicates and total individuals within each lineage or context. The proportion of cross-site replicates is replicates divided by total individuals.*

| test | label | p-value |
| --- | --- | --- |
| fisher | L1b vs L3 | 0.48 |
| fisher | L2a vs L3 | 0.52 |
| fisher | L2b vs L3 | 0.03 |
| fisher | L2 (all) vs L3 | 0.23 |
| fisher | Elite lineages (L1b+L2) vs non-elite lineage L3 | 0.24 |
| fisher | L1b vs L2a | 1.00 |
| fisher | L1b vs L2b | 0.27 |
| fisher | L2a vs L2b | 0.06 |
| fisher | L1b vs L2 (all) | 1.00 |
| chi2 | L1b, L2a, L2b, L3 | 0.05 |
| chi2 | L1b, L2(all), L3 | 0.26 |
| fisher | BS vs ST | 0.005 |
| fisher | T1 vs PT | 0.21 |
| fisher | PT vs W5 | 0.01 |
| fisher | T1 vs W1 | 1.00 |
| fisher | PT vs W1 | 1.00 |
| fisher | T1 vs W5 | 0.02 |
| fisher | Elite tombs (PT+T1) vs non-elite tombs (W5+W1) | 0.01 |
| chi2 | PT, T1, BS, CCP, ST, W5, W1 | 2.6E-07 |
| chi2 | Elite tombs (PT+T1) vs non-elite tombs (W5+W1) | 0.01 |

*Table S5.2. Pairwise and global tests of differences in proportions of cross-site replicates across lineages and mortuary contexts. Fisher exact tests are used for pairwise 2×2 comparisons and χ^2^ tests for multi-group r×2 comparisons.*

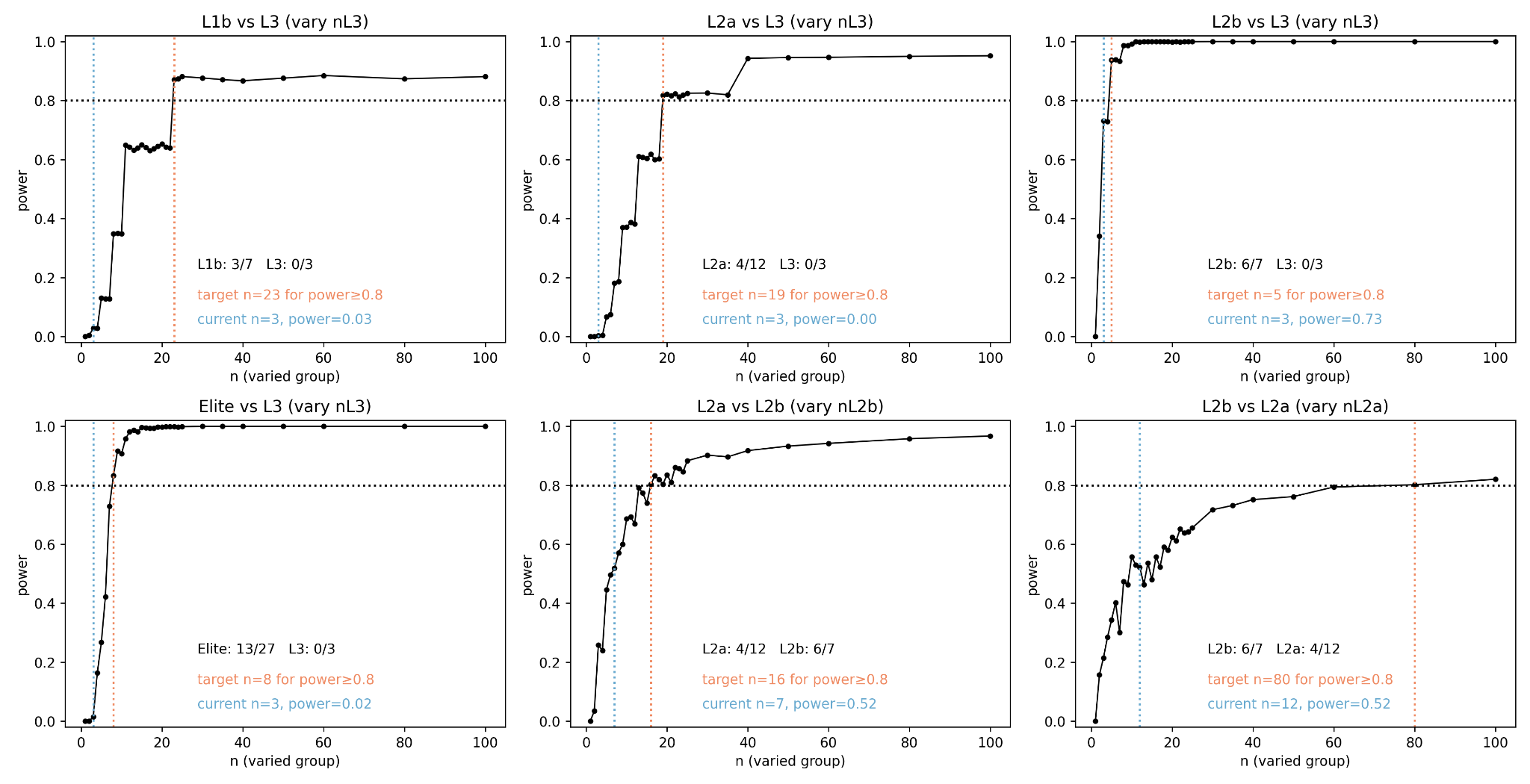

*Figure S5.3. Simulation-based power curves for Fisher exact tests of cross-site replicate proportion differences between lineages. Each panel shows estimated power at α = 0.05 as a function of the sample size of the varied group size, with other parameters fixed (other group size and both group replication proportions). Vertical dotted lines mark the observed sample size for the varied group and the smallest sample size needed to achieve a power of 0.8.*

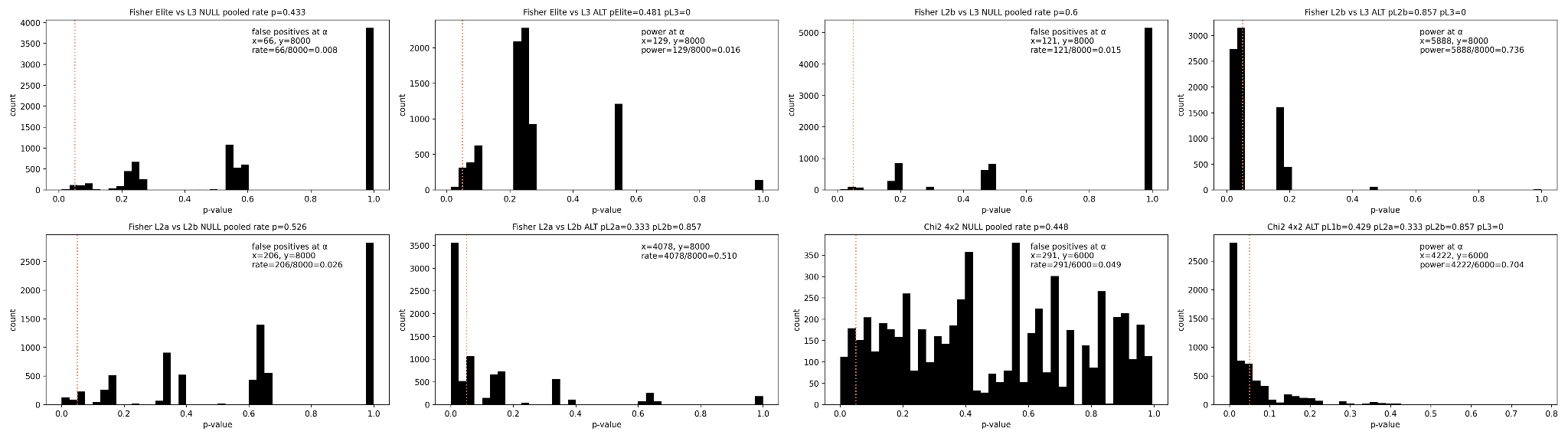

*Figure S5.4. Simulated p-value distributions under null and alternative models for selected lineage Fisher exact tests. Null simulations use a pooled proportion of cross-site replicates (total cross-site replicates across both groups divided by the total individuals across both groups). Alternative simulations use the specific observed proportions of cross-site replicates. Vertical lines mark p = 0.05.*

### 6. Patrilineality in the elite tomb lineages

We tested whether molecular sex is associated with occupying an internal lineage position versus a marrying-in lineage position in the reconstructed elite lineages L1-b and L2. Internal lineage members are individuals with both an ancestor and a descendant within the reconstructed lineage. Marrying-in lineage members are individuals with descendants but no ancestor within the lineage and who are not in the root generation. Root founders and terminal leaves were excluded from this analysis because they do not inform whether lineage continuity passes through males or females.

Among fourteen individuals meeting these criteria (Figure S6.1) and with molecular sex determined, all three marrying-in individuals are female and none are male, whereas ten of eleven internal lineage members are male (Table S6.2). We evaluated the association between molecular sex and lineage role using Fisher exact test on the resulting 2×2 contingency table [[1, 3], [10, 0]], where rows are molecular sex (female, male) and columns are lineage role (internal lineage member, marrying-in). The association is significant (two-sided *p* = 0.01). Because the observed table is at the boundary of what is possible given the fixed margins, the two-sided *p* equals the one-sided lower-tail *p* in the direction of male enrichment among internal lineage members. This indicates a sex bias in how individuals occupy internal pedigree positions, with males more often appearing as internal lineage members and females more often entering the pedigree through marriage.

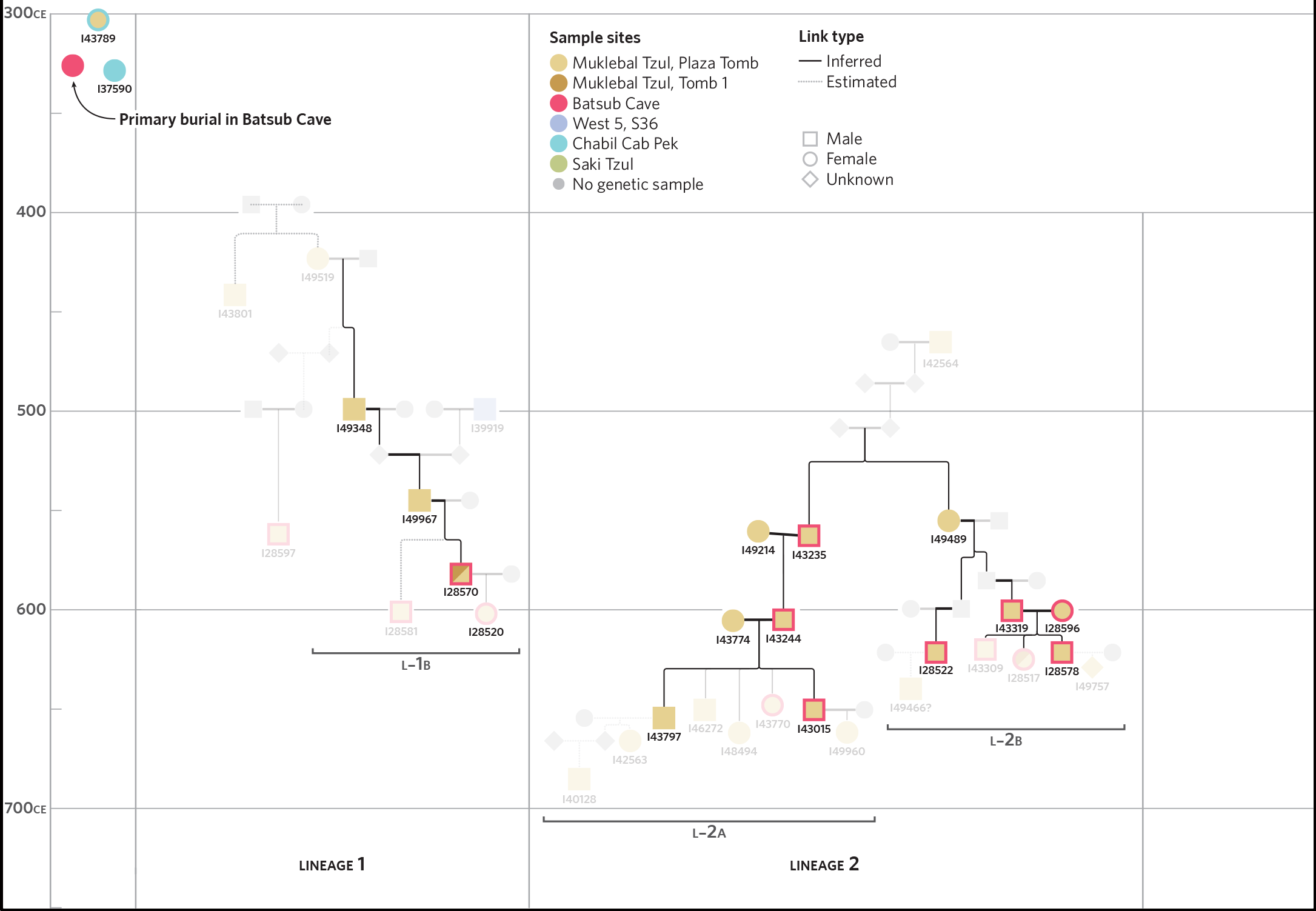

*Figure S6.1. Internal and marrying-in lineage members used in the Fisher exact test for sex-bias in lineality. The pedigree structure is similar to the main lineage figure, but only the fourteen individuals included in the analysis are shown at full opacity. All other lineage members are displayed at 20% opacity for context.*

*Table S6.2. Lineage roles and molecular sex for individuals shown in Figure S6.1. Listed are lineage family assignment, individual ID, molecular sex, pedigree node type (root, internal, leaf, or marry_in), and status used to define internal lineage members (TRUE) and marrying-in (FALSE) lineage members for the Fisher exact test.*

| Family | I-ID | M. Sex | Node type | IsInternal |
| --- | --- | --- | --- | --- |
| L1b | I28520 | F | leaf | NaN |
| L1b | I28570 | M | internal | TRUE |
| L1b | I28581 | M | leaf | NaN |
| L1b | I39919 | M | root | NaN |
| L1b | I49348 | M | internal | TRUE |
| L1b | I49519 | F | root | NaN |
| L1b | I49967 | M | internal | TRUE |
| L2a | I40128 | M | leaf | NaN |
| L2a | I42563 | F | leaf | NaN |
| L2a | I42564 | M | root | NaN |
| L2a | I43015 | M | internal | TRUE |
| L2a | I43235 | M | internal | TRUE |
| L2a | I43244 | M | internal | TRUE |
| L2a | I43770 | F | leaf | NaN |
| L2a | I43774 | F | marry_in | FALSE |
| L2a | I43797 | M | internal | TRUE |
| L2a | I46272 | M | leaf | NaN |
| L2a | I49214 | F | leaf | NaN |
| L2a | I49359 | F | marry_in | FALSE |
| L2a | I49960 | F | leaf | NaN |
| L2b | I28517 | F | leaf | NaN |
| L2b | I28522 | M | internal | TRUE |
| L2b | I28578 | M | internal | TRUE |
| L2b | I28596 | F | marry_in | FALSE |
| L2b | I43319 | M | internal | TRUE |
| L2b | I43555 | M | leaf | NaN |
| L2b | I49489 | F | internal | TRUE |

We also tested for sex-biased mortuary patterns within the elite lineages by asking whether molecular sex among individuals in elite lineages is associated with presence in BS, having any replicate, or having any cross-site replicate. For each comparison, we constructed a 2x2 contingency table and applied both a 𝜒^2^ test of independence and a Fisher exact test. We find no evidence of association between sex and any of these other variables. For presence in BS, the table was [[4 females in elite lineages that are interred in BS, 7 females in elite lineages that are not interred in BS],[9 males in elite lineages that are interred in BS, 7 males in elite lineages that are not interred in BS]] (𝜒^2^ *p* = 0.3, Fisher *p* = 0.4). For having any replicate, the table was [[10 females in elite lineages that have replicates, 1 female in elite lineages that does not have a replicate],[14 males in elite lineages that have replicates, 2 males in elite lineages that do not have replicates]] (𝜒^2^ *p* = 0.8, Fisher *p* = 1). For having any cross-site replicate, the table was [[4 females in elite lineages that have cross-site replicates, 7 females in elite lineages that do not have cross-site replicates], [9 males in elite lineages that have cross-site replicates, 7 males in elite lineages that do not have cross-site replicates]] (𝜒^2^ *p* = 0.3, Fisher *p* = 0.4). Because all the cross-site replicates within these lineages occur at BS, the BS and cross-site replicate contingency tables are identical.

### 7. Pairwise association of mortuary and relationship variables

We performed χ^2^ tests of independence across pairs of variables capturing lineage and family membership, pedigree role, cross site replication status, site presence and per site element counts, molecular sex, Y haplogroup, mtDNA haplogroup, and SNP coverage. For each pair of variables, we constructed a contingency table using pd.crosstab in pandas in python and computed a χ^2^ test of independence using chi2_contingency in scipy.stats in python. We report the χ^2^ statistic, degrees of freedom, and *p* value in Table S7.1. Degrees of freedom depend on the number of distinct values observed for each variable, or 22 in the case of max SNPs, which was binned into equal widths.

Table S7.1:

| **Category 1** | **Category 2** | **Num bins 1** | **Num bins 2** | **Chi-squared** | **DOF** | **P-value** |
| --- | --- | --- | --- | --- | --- | --- |
| Family | Family_withNaNs_assosciatedasfamily | 17 | 10 | 828 | 144 | 1.45E-96 |
| number_duplicates | MKB PT | 14 | 12 | 468 | 143 | 3.69E-36 |
| number_duplicates | BS | 14 | 6 | 226 | 65 | 1.13E-19 |
| EliteLineageFamiliesMKB | NonLineageNonFamily | 2 | 2 | 77 | 1 | 1.68E-18 |
| Ifpresent_BS_CCP | BS | 2 | 6 | 88 | 5 | 2.23E-17 |
| Family_withNaNs_assosciatedasfamily | EliteLineageFamiliesMKB | 10 | 2 | 92 | 9 | 6.46E-16 |
| Family_withNaNs_assosciatedasfamily | NonLineageNonFamily | 10 | 2 | 92 | 9 | 6.46E-16 |
| Family_withNaNs_assosciatedasfamily | NonEliteLineage | 10 | 2 | 92 | 9 | 6.46E-16 |
| **if_cross_site_duplicate** | **Ifpresent_BS_CCP** | **2** | **2** | **63** | **1** | **2.50E-15** |
| Ifpresent_PT_T1 | MKB PT | 2 | 12 | 87 | 11 | 5.75E-14 |
| if_cross_site_duplicate | BS | 2 | 6 | 68 | 5 | 2.59E-13 |
| M. Sex | Y hg | 2 | 7 | 70 | 6 | 4.23E-13 |
| Family | NonLineageNonFamily | 17 | 2 | 92 | 16 | 1.07E-12 |
| Family | EliteLineageFamiliesMKB | 17 | 2 | 92 | 16 | 1.07E-12 |
| Family | NonEliteLineage | 17 | 2 | 92 | 16 | 1.07E-12 |
| Ifpresent_PT_T1 | MKB W5 S36 | 2 | 3 | 49 | 2 | 2.11E-11 |
| BS | MKB T1 | 6 | 3 | 68 | 10 | 1.11E-10 |
| Family | ST | 17 | 3 | 102 | 32 | 2.89E-09 |
| number_duplicates | MKB T1 | 14 | 3 | 91 | 26 | 3.34E-09 |
| Family_withNaNs_assosciatedasfamily | mtDNA hg | 10 | 29 | 404 | 252 | 3.56E-09 |
| if_cross_site_duplicate | number_duplicates | 2 | 14 | 65 | 13 | 6.96E-09 |
| NonEliteLineage | mtDNA hg | 2 | 29 | 92 | 28 | 9.63E-09 |
| CCP | mtDNA hg | 2 | 29 | 92 | 28 | 9.63E-09 |
| **Family** | **mtDNA hg** | **17** | **29** | **633** | **448** | **1.71E-08** |
| Family_withNaNs_assosciatedasfamily | ST | 10 | 3 | 72 | 18 | 2.11E-08 |
| if_cross_site_duplicate | MKB PT | 2 | 12 | 55 | 11 | 6.44E-08 |
| Family | if_cross_site_duplicate | 17 | 2 | 56 | 16 | 2.06E-06 |
| BS | MKB PT | 6 | 12 | 117 | 55 | 2.12E-06 |
| Family | MKB PT | 17 | 12 | 276 | 176 | 2.14E-06 |
| number_duplicates | Ifpresent_BS_CCP | 14 | 2 | 49 | 13 | 3.94E-06 |
| **Ifpresent_BS_CCP** | **MKB PT** | **2** | **12** | **40** | **11** | **3.98E-05** |
| Y hg | mtDNA hg | 7 | 29 | 250 | 168 | 4.05E-05 |
| Y hg | max SNPs | 7 | 22 | 195 | 126 | 7.77E-05 |
| Family | number_duplicates | 17 | 14 | 291 | 208 | 0.0001 |
| Family | Ifpresent_BS_CCP | 17 | 2 | 44 | 16 | 0.0002 |
| MKB W5 S36 | NonEliteLineage | 3 | 2 | 16 | 2 | 0.0003 |
| **if_cross_site_duplicate** | **Ifpresent_PT_T1** | **2** | **2** | **12** | **1** | **0.0004** |
| MKB T1 | mtDNA hg | 3 | 29 | 96 | 56 | 0.0007 |
| Family_withNaNs_assosciatedasfamily | MKB PT | 10 | 12 | 149 | 99 | 0.0008 |
| Family | max SNPs | 17 | 22 | 421 | 336 | 0.0010 |
| MKB PT | MKB W5 S36 | 12 | 3 | 47 | 22 | 0.0016 |
| Family | Ifpresent_PT_T1 | 17 | 2 | 38 | 16 | 0.0017 |
| **number_duplicates** | **EliteLineageFamiliesMKB** | **14** | **2** | **32** | **13** | **0.0028** |
| Ifpresent_PT_T1 | ST | 2 | 3 | 11 | 2 | 0.0050 |
| Family_withNaNs_assosciatedasfamily | Ifpresent_PT_T1 | 10 | 2 | 23 | 9 | 0.0073 |
| MKB W5 S36 | mtDNA hg | 3 | 29 | 85 | 56 | 0.0075 |
| MKB W5 S36 | max SNPs | 3 | 22 | 67 | 42 | 0.0088 |
| **Ifpresent_BS_CCP** | **MKB W5 S36** | **2** | **3** | **9** | **2** | **0.0092** |
| if_cross_site_duplicate | Y hg | 2 | 7 | 17 | 6 | 0.0097 |
| MKB PT | EliteLineageFamiliesMKB | 12 | 2 | 24 | 11 | 0.0113 |
| number_duplicates | NonLineageNonFamily | 14 | 2 | 27 | 13 | 0.0136 |
| M. Sex | max SNPs | 2 | 22 | 37 | 21 | 0.0158 |
| MKB T1 | MKB PT | 3 | 12 | 37 | 22 | 0.0209 |
| **if_cross_site_duplicate** | **MKB W5 S36** | **2** | **3** | **8** | **2** | **0.0220** |
| **Family_withNaNs_assosciatedasfamily** | **number_duplicates** | **10** | **14** | **150** | **117** | **0.0225** |
| NonLineageNonFamily | max SNPs | 2 | 22 | 36 | 21 | 0.0244 |
| NonEliteLineage | Y hg | 2 | 7 | 14 | 6 | 0.0260 |
| Ifpresent_PT_T1 | NonEliteLineage | 2 | 2 | 5 | 1 | 0.0312 |
| Family | BS | 17 | 6 | 105 | 80 | 0.0337 |
| MKB PT | M. Sex | 12 | 2 | 21 | 11 | 0.0357 |
| EliteLineageFamiliesMKB | max SNPs | 2 | 22 | 34 | 21 | 0.0363 |
| MKB PT | NonLineageNonFamily | 12 | 2 | 21 | 11 | 0.0365 |
| EliteLineageFamiliesMKB | Y hg | 2 | 7 | 13 | 6 | 0.0472 |
| if_cross_site_duplicate | max SNPs | 2 | 22 | 33 | 21 | 0.0473 |
| Ifpresent_PT_T1 | EliteLineageFamiliesMKB | 2 | 2 | 4 | 1 | 0.0508 |
| number_duplicates | Ifpresent_PT_T1 | 14 | 2 | 22 | 13 | 0.0564 |
| Family_withNaNs_assosciatedasfamily | M. Sex | 10 | 2 | 17 | 9 | 0.0568 |
| NonLineageNonFamily | Y hg | 2 | 7 | 12 | 6 | 0.0630 |
| Ifpresent_BS_CCP | Y hg | 2 | 7 | 12 | 6 | 0.0664 |
| M. Sex | mtDNA hg | 2 | 29 | 39 | 28 | 0.0755 |
| NonLineageNonFamily | M. Sex | 2 | 2 | 3 | 1 | 0.0795 |
| if_cross_site_duplicate | EliteLineageFamiliesMKB | 2 | 2 | 3 | 1 | 0.0798 |
| EliteLineageFamiliesMKB | mtDNA hg | 2 | 29 | 39 | 28 | 0.0805 |
| NonLineageNonFamily | mtDNA hg | 2 | 29 | 39 | 28 | 0.0817 |
| Ifpresent_PT_T1 | max SNPs | 2 | 22 | 30 | 21 | 0.0864 |
| number_duplicates | M. Sex | 14 | 2 | 20 | 13 | 0.0932 |
| MKB W5 S36 | EliteLineageFamiliesMKB | 3 | 2 | 5 | 2 | 0.0954 |
| **if_cross_site_duplicate** | **M. Sex** | **2** | **2** | **3** | **1** | **0.1032** |
| Ifpresent_PT_T1 | Y hg | 2 | 7 | 10 | 6 | 0.1092 |
| ST | EliteLineageFamiliesMKB | 3 | 2 | 4 | 2 | 0.1126 |
| Ifpresent_PT_T1 | BS | 2 | 6 | 9 | 5 | 0.1279 |
| Ifpresent_BS_CCP | Ifpresent_PT_T1 | 2 | 2 | 2 | 1 | 0.1413 |
| Ifpresent_BS_CCP | max SNPs | 2 | 22 | 28 | 21 | 0.1423 |
| ST | NonLineageNonFamily | 3 | 2 | 4 | 2 | 0.1471 |
| MKB W5 S36 | M. Sex | 3 | 2 | 4 | 2 | 0.1480 |
| EliteLineageFamiliesMKB | M. Sex | 2 | 2 | 2 | 1 | 0.1484 |
| Family | M. Sex | 17 | 2 | 22 | 16 | 0.1574 |
| Family_withNaNs_assosciatedasfamily | if_cross_site_duplicate | 10 | 2 | 13 | 9 | 0.1620 |
| MKB PT | mtDNA hg | 12 | 29 | 332 | 308 | 0.1632 |
| MKB PT | Y hg | 12 | 7 | 77 | 66 | 0.1646 |
| if_cross_site_duplicate | NonLineageNonFamily | 2 | 2 | 2 | 1 | 0.1737 |
| NonEliteLineage | max SNPs | 2 | 22 | 26 | 21 | 0.2081 |
| **Ifpresent_BS_CCP** | **M. Sex** | **2** | **2** | **2** | **1** | **0.2105** |
| if_cross_site_duplicate | MKB T1 | 2 | 3 | 3 | 2 | 0.2146 |
| MKB W5 S36 | Y hg | 3 | 7 | 15 | 12 | 0.2334 |
| EliteLineageFamiliesMKB | NonEliteLineage | 2 | 2 | 1 | 1 | 0.2560 |
| NonEliteLineage | NonLineageNonFamily | 2 | 2 | 1 | 1 | 0.2720 |
| Family_withNaNs_assosciatedasfamily | Y hg | 10 | 7 | 59 | 54 | 0.2928 |
| Ifpresent_BS_CCP | MKB T1 | 2 | 3 | 2 | 2 | 0.3019 |
| MKB PT | max SNPs | 12 | 22 | 241 | 231 | 0.3140 |
| Family_withNaNs_assosciatedasfamily | MKB W5 S36 | 10 | 3 | 20 | 18 | 0.3230 |
| Ifpresent_PT_T1 | NonLineageNonFamily | 2 | 2 | 1 | 1 | 0.3384 |
| Family_withNaNs_assosciatedasfamily | Ifpresent_BS_CCP | 10 | 2 | 10 | 9 | 0.3482 |
| Ifpresent_BS_CCP | ST | 2 | 3 | 2 | 2 | 0.3636 |
| Ifpresent_PT_T1 | mtDNA hg | 2 | 29 | 30 | 28 | 0.3770 |
| MKB W1 | max SNPs | 2 | 22 | 22 | 21 | 0.3857 |
| BS | M. Sex | 6 | 2 | 5 | 5 | 0.4069 |
| if_cross_site_duplicate | mtDNA hg | 2 | 29 | 29 | 28 | 0.4076 |
| Ifpresent_BS_CCP | EliteLineageFamiliesMKB | 2 | 2 | 1 | 1 | 0.4165 |
| if_cross_site_duplicate | ST | 2 | 3 | 2 | 2 | 0.4388 |
| BS | EliteLineageFamiliesMKB | 6 | 2 | 5 | 5 | 0.4706 |
| ST | M. Sex | 3 | 2 | 1 | 2 | 0.4809 |
| Ifpresent_PT_T1 | M. Sex | 2 | 2 | 0 | 1 | 0.4885 |
| MKB T1 | M. Sex | 3 | 2 | 1 | 2 | 0.4890 |
| Family | MKB W5 S36 | 17 | 3 | 32 | 32 | 0.4906 |
| MKB W5 S36 | NonLineageNonFamily | 3 | 2 | 1 | 2 | 0.5043 |
| number_duplicates | MKB W5 S36 | 14 | 3 | 25 | 26 | 0.5370 |
| BS | MKB W5 S36 | 6 | 3 | 9 | 10 | 0.5401 |
| Ifpresent_PT_T1 | MKB T1 | 2 | 3 | 1 | 2 | 0.5429 |
| Ifpresent_BS_CCP | NonEliteLineage | 2 | 2 | 0 | 1 | 0.5493 |
| number_duplicates | max SNPs | 14 | 22 | 267 | 273 | 0.5883 |
| MKB T1 | EliteLineageFamiliesMKB | 3 | 2 | 1 | 2 | 0.5892 |
| BS | NonLineageNonFamily | 6 | 2 | 4 | 5 | 0.5960 |
| mtDNA hg | max SNPs | 29 | 22 | 578 | 588 | 0.6098 |
| if_cross_site_duplicate | CCP | 2 | 2 | 0 | 1 | 0.6274 |
| Ifpresent_PT_T1 | MKB W1 | 2 | 2 | 0 | 1 | 0.6274 |
| MKB T1 | NonLineageNonFamily | 3 | 2 | 1 | 2 | 0.6285 |
| number_duplicates | CCP | 14 | 2 | 11 | 13 | 0.6430 |
| if_cross_site_duplicate | NonEliteLineage | 2 | 2 | 0 | 1 | 0.6502 |
| Family_withNaNs_assosciatedasfamily | BS | 10 | 6 | 41 | 45 | 0.6534 |
| Ifpresent_BS_CCP | mtDNA hg | 2 | 29 | 24 | 28 | 0.6640 |
| Ifpresent_BS_CCP | NonLineageNonFamily | 2 | 2 | 0 | 1 | 0.7058 |
| Ifpresent_BS_CCP | CCP | 2 | 2 | 0 | 1 | 0.7091 |
| Family_withNaNs_assosciatedasfamily | max SNPs | 10 | 22 | 178 | 189 | 0.7110 |
| number_duplicates | mtDNA hg | 14 | 29 | 348 | 364 | 0.7205 |
| MKB PT | NonEliteLineage | 12 | 2 | 7 | 11 | 0.7602 |
| number_duplicates | Y hg | 14 | 7 | 69 | 78 | 0.7617 |
| NonEliteLineage | M. Sex | 2 | 2 | 0 | 1 | 0.7863 |
| ST | mtDNA hg | 3 | 29 | 46 | 56 | 0.8217 |
| Family | CCP | 17 | 2 | 11 | 16 | 0.8326 |
| Family | Y hg | 17 | 7 | 82 | 96 | 0.8436 |
| BS | Y hg | 6 | 7 | 22 | 30 | 0.8613 |
| MKB W5 S36 | MKB W1 | 3 | 2 | 0 | 2 | 0.8990 |
| MKB W5 S36 | CCP | 3 | 2 | 0 | 2 | 0.8990 |
| ST | Y hg | 3 | 7 | 6 | 12 | 0.9182 |
| BS | NonEliteLineage | 6 | 2 | 1 | 5 | 0.9213 |
| MKB W5 S36 | ST | 3 | 3 | 1 | 4 | 0.9274 |
| ST | NonEliteLineage | 3 | 2 | 0 | 2 | 0.9319 |
| ST | max SNPs | 3 | 22 | 29 | 42 | 0.9432 |
| MKB T1 | Y hg | 3 | 7 | 5 | 12 | 0.9489 |
| MKB T1 | NonEliteLineage | 3 | 2 | 0 | 2 | 0.9491 |
| Family_withNaNs_assosciatedasfamily | MKB T1 | 10 | 3 | 9 | 18 | 0.9495 |
| MKB T1 | MKB W5 S36 | 3 | 3 | 1 | 4 | 0.9570 |
| MKB W1 | NonLineageNonFamily | 2 | 2 | 0 | 1 | 0.9651 |
| CCP | NonLineageNonFamily | 2 | 2 | 0 | 1 | 0.9651 |
| Family | MKB T1 | 17 | 3 | 18 | 32 | 0.9758 |
| MKB W1 | ST | 2 | 3 | 0 | 2 | 0.9773 |
| ST | CCP | 3 | 2 | 0 | 2 | 0.9773 |
| MKB T1 | MKB W1 | 3 | 2 | 0 | 2 | 0.9831 |
| MKB T1 | CCP | 3 | 2 | 0 | 2 | 0.9831 |
| number_duplicates | ST | 14 | 3 | 13 | 26 | 0.9836 |
| MKB PT | ST | 12 | 3 | 10 | 22 | 0.9857 |
| BS | MKB W1 | 6 | 2 | 0 | 5 | 0.9933 |
| BS | CCP | 6 | 2 | 0 | 5 | 0.9933 |
| BS | mtDNA hg | 6 | 29 | 100 | 140 | 0.9958 |
| MKB PT | MKB W1 | 12 | 2 | 2 | 11 | 0.9963 |
| BS | max SNPs | 6 | 22 | 70 | 105 | 0.9964 |
| BS | ST | 6 | 3 | 2 | 10 | 0.9969 |
| MKB W1 | Y hg | 2 | 7 | 1 | 6 | 0.9969 |
| CCP | Y hg | 2 | 7 | 1 | 6 | 0.9969 |
| MKB PT | CCP | 12 | 2 | 2 | 11 | 0.9971 |
| MKB T1 | ST | 3 | 3 | 0 | 4 | 0.9976 |
| Family_withNaNs_assosciatedasfamily | CCP | 10 | 2 | 1 | 9 | 0.9992 |
| Family_withNaNs_assosciatedasfamily | MKB W1 | 10 | 2 | 1 | 9 | 0.9992 |
| number_duplicates | NonEliteLineage | 14 | 2 | 2 | 13 | 0.9992 |
| MKB W1 | mtDNA hg | 2 | 29 | 8 | 28 | 0.9999 |
| CCP | max SNPs | 2 | 22 | 4 | 21 | 1.0000 |
| Family | MKB W1 | 17 | 2 | 2 | 16 | 1.0000 |
| number_duplicates | MKB W1 | 14 | 2 | 1 | 13 | 1.0000 |
| MKB T1 | max SNPs | 3 | 22 | 7 | 42 | 1.0000 |
| CCP | EliteLineageFamiliesMKB | 2 | 2 | 0 | 1 | 1.0000 |
| MKB W1 | NonEliteLineage | 2 | 2 | 0 | 1 | 1.0000 |
| Ifpresent_BS_CCP | MKB W1 | 2 | 2 | 0 | 1 | 1.0000 |
| MKB W1 | CCP | 2 | 2 | 0 | 1 | 1.0000 |
| MKB W1 | M. Sex | 2 | 2 | 0 | 1 | 1.0000 |
| CCP | M. Sex | 2 | 2 | 0 | 1 | 1.0000 |
| MKB W1 | EliteLineageFamiliesMKB | 2 | 2 | 0 | 1 | 1.0000 |
| CCP | NonEliteLineage | 2 | 2 | 0 | 1 | 1.0000 |
| Ifpresent_PT_T1 | CCP | 2 | 2 | 0 | 1 | 1.0000 |
| if_cross_site_duplicate | MKB W1 | 2 | 2 | 0 | 1 | 1.0000 |

### 8. Asymmetrical BS-PT replication

All analyses in this section are restricted to individuals with genome-wide data from at least one skeletal element yielding ≥50,000 SNPs.

Overlap between BS and PT is highly asymmetric. We summarized the individual-level overlap using the 2×2 contingency table [24, 6; 24, 64] (Table S8.1), where rows indicate context (BS or PT) and columns indicate whether an individual is also represented in the other context. A Fisher exact test on this table is significant ($p=5\cdot{10}^{-7}$) and confirms strong deviation from independence. Directionally, individuals found in BS are disproportionately also found in PT, whereas individuals found in PT are disproportionately not also found in BS.

*Table S8.1. BS–PT overlap among individuals with ≥50,000 SNPs.*

| **Context (row)** | **Also found in the other context** | **Not found in the other context** | **Row total** |
| --- | --- | --- | --- |
| BS | 24 | 6 | 30 |
| PT | 24 | 64 | 88 |
| Column Total | 48 | 70 | 118 |

The Fisher test treats each individual as a single per-individuals observation with two options (presence of cross-site replication / absence of cross-site replication) and therefore does not explicitly model the fact that individuals contribute different numbers of genotyped elements. To verify that the directional asymmetry of the Fisher test is not an artifact of sampling depth, we performed a multivariate-hypergeometric simulation where we pool all elements across BS and PT, and then randomly redistribute the “BS” and “PT” labels among the pooled elements. The total number of samples and the total number of BS and PT samples in the simulation is kept the same as the observed totals. The total number of skeletal elements for each individual in the simulation is also kept the same as the observed totals. In each of the 50,000 simulations, we randomly assigned the skeletal elements for each individual “BS” or “PT” labels using the hygernd function that is used to allocate a fixed number of “BS” labels across the skeletal elements of individuals, each with a fixed number of skeletal elements. We then collapsed back to individuals and computed three statistics: (i) the number of individuals found in both BS and PT, (ii) the proportion of BS individuals also present in PT, and (iii) the proportion of PT individuals also present in BS.

```
% Multivariate hypergeometric null:
% Fix each individual's total number of samples (s_i).
% Fix the total number of BS (Ncave) and elite (S - Ncave) samples (pooled
% accross all individuals), where S is the total number of samples across
% all individuals.
% Randomly allocate exactly Ncave cave labels across the S elements for
% interment in BS. The rest are PT.
% Then count how many individuals have both cave and elite elements and
% compare the simulated proportions to the observed proportions.

% Fisher exact test for assymetric BS-PT overlap (Table S8.1).
[~,p,~]=fishertest([24, 6; 24, 64]);

T = readtable("iid_site_counts_minsnps.tsv","FileType","text","Delimiter","\t");

% Adjust these if your file differs:
BS  = T{:,7};
T1  = T{:,8};
PT  = T{:,9};

% Replace NaN with 0
BS(isnan(BS)) = 0;
T1(isnan(T1)) = 0;
PT(isnan(PT)) = 0;

Cave  = BS;
Elite = PT;

s = Cave + Elite;
keep = s > 0;
Cave = Cave(keep);
Elite = Elite(keep);
s = s(keep);

Ncave = sum(Cave);
S = sum(s);

obs_n_cave   = sum(Cave > 0);
obs_n_elite  = sum(Elite > 0);
obs_overlap = sum((Cave > 0) & (Elite > 0));
obs_p_elite_given_cave = obs_overlap / obs_n_cave;
obs_p_cave_given_elite = obs_overlap / obs_n_elite;

nsim = 50000;
rng(1);
sim_overlap = zeros(1, nsim);
sim_n_cave  = zeros(1, nsim);
sim_n_elite = zeros(1, nsim);
sim_p_elite_given_cave = zeros(1, nsim);
sim_p_cave_given_elite = zeros(1, nsim);

for r = 1:nsim
    K = Ncave;
    N = S;
    CaveSim = zeros(size(s));

    % sequential hypergeometric allocation across individuals
    for i = 1:numel(s)
        n_i = s(i);
        if i == numel(s)
            x = K;  % whatever remains must go to last category
        else
            % draw x ~ Hypergeom(successes=K in population N, draws=n_i)
            x = hygernd(N, K, n_i);
        end
        CaveSim(i) = x;
        K = K - x;
        N = N - n_i;
    end

    EliteSim = s - CaveSim;

    isC = CaveSim > 0;
    isE = EliteSim > 0;

    ov = sum(isC & isE);
    nc = sum(isC);
    ne = sum(isE);

    sim_overlap(r) = ov;
    sim_n_cave(r)  = nc;
    sim_n_elite(r) = ne;

    sim_p_elite_given_cave(r) = ov / nc;
    sim_p_cave_given_elite(r) = ov / ne;
end

% Empirical one-sided p-values (enrichment: simulated >= observed)
p_ov_right  = (1 + sum(sim_overlap >= obs_overlap)) / (1 + nsim); % overlap of 24 individuals (not contingent on being in BS or in PT) is not higher than expected
p_ov_left  = (1 + sum(sim_overlap <= obs_overlap)) / (1 + nsim); % overlap or 24 individuals (not contingent on being in BS or in PT) is actually lower than expected
p_pEC_right = (1 + sum(sim_p_elite_given_cave >= obs_p_elite_given_cave)) / (1 + nsim); % BS to PT: observed proportion 0.80 is higher than expected
p_pEC_left = (1 + sum(sim_p_elite_given_cave <= obs_p_elite_given_cave)) / (1 + nsim); % BS to PT: observed proportion 0.80 is not lower than expected
p_pCE_right = (1 + sum(sim_p_cave_given_elite >= obs_p_cave_given_elite)) / (1 + nsim); % PT to BS: observed proportion 0.27 is not higher than expected
p_pCE_left = (1 + sum(sim_p_cave_given_elite <= obs_p_cave_given_elite)) / (1 + nsim); % PT to BS: observed proportion 0.27 is not higher than expected

fprintf("S=%d total elements, Kcave=%d fixed cave elements, nsim=%d\n", S, Ncave, nsim);
fprintf("Observed: n_cave=%d, n_elite=%d, overlap=%d\n", obs_n_cave, obs_n_elite, obs_overlap);
fprintf("Observed: P(elite|cave)=%.4f, P(cave|elite)=%.4f\n", obs_p_elite_given_cave, obs_p_cave_given_elite);
fprintf("Empirical: p (>=obs) overlap=%.6g, p (<=obs): overlap=%.6g, p (>=obs) (elite|cave)=%.6g, p (<=obs) (elite|cave)=%.6g, p (>=obs) (cave|elite)=%.6g, p (<=obs) (cave|elite)=%.6g\n", p_ov_right, p_ov_left, p_pEC_right, p_pEC_left, p_pCE_right, p_pCE_left);

% ---- Plots ----
figure; tiledlayout(1,3, "Padding","compact", "TileSpacing","compact");

nexttile;
histogram(sim_overlap, 'BinMethod','integers');
xline(obs_overlap, ':', 'LineWidth', 2);
xlabel('overlap count'); ylabel('count');
title('Null: overlap');

nexttile;
histogram(sim_p_elite_given_cave, 40);
xline(obs_p_elite_given_cave, ':', 'LineWidth', 2);
xlabel('P(elite | cave)'); ylabel('count');
title('Null: P(elite|cave)');

nexttile;
histogram(sim_p_cave_given_elite, 40);
xline(obs_p_cave_given_elite, ':', 'LineWidth', 2);
xlabel('P(cave | elite)'); ylabel('count');
title('Null: P(cave|elite)');

set(gcf,'Color','w');
exportgraphics(gcf,'mvhypergeom_3tests.pdf');
```

Under this null model, the observed fraction of BS individuals also present in PT is 24/30 (0.80), which is higher than expected by chance (Figure S8.2, right-tailed empirical p ≈ 0.04). In contrast, the observed fraction of PT individuals also present in BS is 24/88 (0.27), which is lower than expected by chance (left-tailed empirical *p* ≈ 3.6×10⁻⁴). Together, these results reinforce an asymmetric relationship in which BS draws selectively from a non-random subset of PT individuals, rather than reflecting symmetric overlap driven by sampling depth.

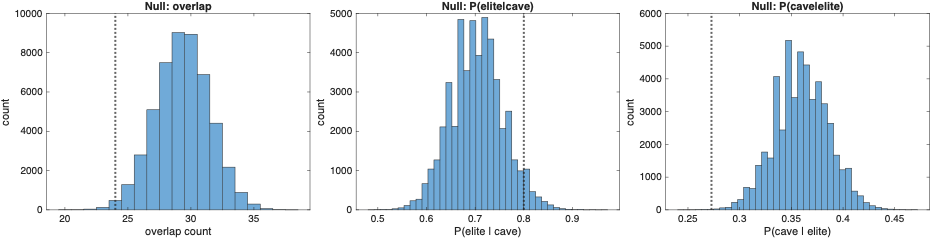

*Figure S8.2. Null distributions from the multivariate-hypergeometric simulation for (i) the number of individuals represented in both BS and PT, (ii) P(PT|BS), and (iii) P(BS|PT). Vertical dotted lines mark the observed values.*

Given that individuals found at BS are from a non-random subset of PT individuals and given the apparent increase in PT–BS cross-site interment in later generations (Figure 2), we tested whether overlap is uniform through time. We split dated PT individuals into those before versus after ~520 CE and tabulated whether they are also represented at BS (Table S8.3). Fisher exact testing indicates a significant shift toward greater BS overlap in later PT individuals (two-sided *p* = 0.0408; one-sided *p* = 0.0345 in the direction of enrichment after ~520 CE).

*Table S8.3. Temporal shift in PT to BS overlap (before vs after ~520 CE)*

| **PT date bin** | **Also found in BS** | **Not found in BS** | **Row total** |
| --- | --- | --- | --- |
| Before ~520 CE | 1 | 6 | 7 |
| After ~520 CE | 21 | 14 | 35 |
| Total | 22 | 20 | 42 |
